# Variation in near-surface soil temperature drives plant assemblage insurance potential

**DOI:** 10.1101/2022.11.21.517364

**Authors:** Elizabeth G. Simpson, Ian Fraser, Hillary Woolf, William D. Pearse

## Abstract

1. Studying how assemblages vary across environmental gradients provides a baseline for how assemblages may respond to climate change. Per the biological insurance hypothesis, assemblages with more variation in functional diversity will maintain ecosystem functions when species are lost. In complement, environmental heterogeneity supports landscape-scale ecosystem functionality (*i*.*e*. spatial insurance), when that variation includes environments with more abundant resources.
2. We use the relationship between vascular plant functional diversity and microenvironment to identify where assemblages are most likely to maintain functionality in a mountainous fieldsite in northeastern Utah, USA. We assessed how life history strategies and information about phylogenetic differences affect these diversity-environment relationships.
3. We found less functionally dispersed assemblages, that were shorter and more resource-conservative on hotter, more variable, south-facing slopes. In contrast, we found more functionally dispersed assemblages, that were taller and more resource-acquisitive on cooler, less variable, north-facing slopes. Herbaceous and woody perennials drove these trends. Additionally, including information about phylogenetic differences in a dispersion metric indicated that phylogeny accounts for traits we did not measure.
4. *Synthesis*. At our fieldsite, soil temperature acts as an environmental filter across aspect. If soil temperature increases and becomes more variable, the function of north- vs. south-facing assemblages may be at risk for contrasting reasons. On south-facing slopes, assemblages may not have the variance in functional diversity needed to respond to more intense, stressful conditions. Conversely, assemblages on north-facing slopes may not have the resource-conservative strategies needed to persist if temperatures become hotter and more variable. We suggest that studying dispersal traits, especially of perennial species, will provide additional insight into whether this landscape will maintain function as climate changes.

## 1 Introduction

As Earth’s climate warms and becomes more variable (IPCC 2018), ecological assemblages face new environmental conditions that cause species loss. Biotic and abiotic factors influence whether species loss affects overall ecosystem functionality. The biological insurance hypothesis (Loreau et al. 2021; Yachi and Loreau 1999) proposes that an assemblage with more variation in species’ functional responses to environmental stressors better maintains ecological function, even if few species perform a given function (Mori et al. 2013; Suding et al. 2008; Elmqvist et al. 2003). In complement, environmental heterogeneity supports landscape-scale ecosystem functionality when patches with less stressful conditions and more abundant resources support species that would otherwise go locally extinct (Greiser et al. 2020; Maclean et al. 2015). Quantifying how functional diversity varies across environmental gradients identifies the environmental conditions where assemblages are best equipped to maintain functionality. These relationships provide a foundation for monitoring and management decisions that protect ecosystem functioning (Díaz et al. 2018; Cardinale et al. 2012).

Species’ characteristics, interactions, and dispersal rates respond to resource gradients in a spatial context (Leibold et al. 2004). In line with this framework, functional traits reflect how plants acquire resources and affect the ecosystem around them (Reich 2014; Mason and de Bello 2013; Suding et al. 2008; Lavorel and Garnier 2002). For example, the leaf economic spectrum describes how plants invest resources into their leaves (Díaz et al. 2016; Wright et al. 2004). In resource-poor, variable environments plants tend to invest resources into leaves that last longer and produce more photosynthate over longer timescales, a conservative strategy. In contrast, plants in resource-rich environments tend to invest fewer resources into leaves that will not last as long but produce more photosynthate in a shorter time span, an acquisitive strategy.

The mean and variance in functional diversity metrics calculated from functional traits provide complementary information about the current and future functionality of assemblages. Individual traits, summarized at the assemblage level as the community weighted mean (CWM) of that trait (Lavorel et al. 2008), represent the strategies plants most commonly use in an environment. The CWM of traits often shifts as the environment changes, as a result of phenotypic plasticity and/or species turnover. For example, as summer temperatures warmed in the Arctic, assemblages grew taller as a result of immigration by taller, but still local, species (Bjorkman et al. 2018). However, leaf traits only responded at wetter locations, where species invested fewer resources into leaves, potentially allocating those resources to higher growth rates. The variance in all measured functional traits (*e*.*g*. functional dispersion; Laliberté and Legendre 2010) provides insight into the environmental conditions where species are relatively more or less likely to maintain function as conditions change in the future. For example, in high alpine meadows in Colorado, assemblages had more variance in plant functional strategies in response to an increase in spatially variable environmental conditions (Stark et al. 2017).

An assemblage well-equipped to maintain ecological function contains species with more variation in functional response traits (Lavorel and Garnier 2002; Díaz et al. 2013). However, these traits must not be strongly correlated with the effect traits that provide ecosystem services (Díaz et al. 2013). For example, per this hypothesis, an assemblage with more variation in resource investment strategies (measured as response traits) will maintain more consistent productivity (measured by effect traits) in response to variation in water availability, as long as resource investment and productivity traits are not correlated. However, a global aggregation of local studies found that, broadly, plant assemblages contain less variation in functional traits than expected by chance (Bruelheide et al. 2018). This finding indicates a need to understand the environmental conditions where plant assemblages have more biological insurance potential locally.

While environmental changes (*e*.*g*., increased moisture) may result in more resources for species, others (*e*.*g*., increased temperature) may cause more stress that plants need to adapt to. We hypothesize that the mean and variance of plant traits summarized at the assemblage level will reflect current differences in resource availability and environmental stress. To test this hypothesis, we look at how the mean and variance in functional traits vary across microenvironment — near-surface soil temperature and soil texture — at a fieldsite in northeastern Utah (Figure 1). We identify which strategies plants most commonly use in different environments using the CWM of functional traits that provide insight into the tradeoff between resource abundance and environmental stress. In complement, we identify where assemblages will likely maintain function in the future by assessing the variance in all measured functional traits. Since life history strategies typically encompass similar functional strategies (Díaz et al. 2016), we test whether particular life history groups drive these diversity-environment relationships. If a life history group drives these diversity-environment relationships, this group may be a good candidate for continued monitoring and management. Finally, we assess whether including information about phylogenetic differences between species changes the relationship between function and microenvironment. Taken together, the results of this assessment guide management actions by determining where and how plant assemblages will be able to maintain functions and ecosystem services.

**Figure 1:**
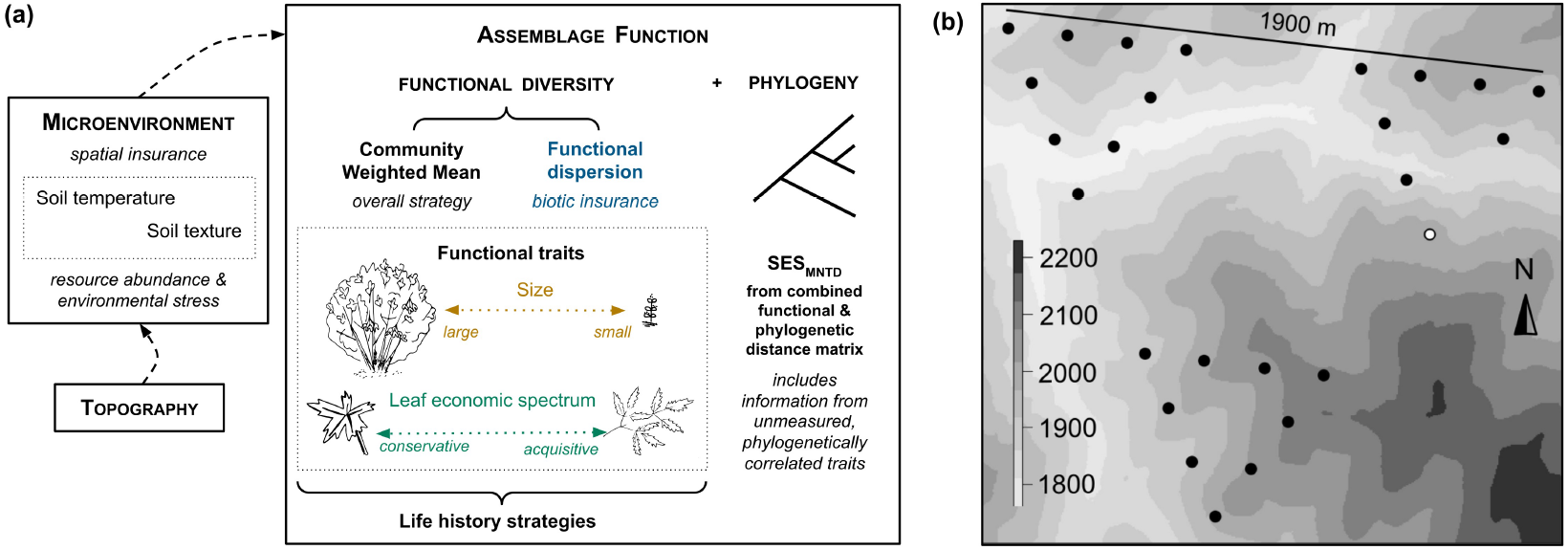
Assessing changes in functional diversity across microenvironment identifies where assemblages have the highest potential of maintaining function in response to changing conditions. (a) Environmental heterogeneity, often driven by differences in topography, provides spatial insurance when microenvironments with more resources and/or less stressful conditions support species that would otherwise be lost as climate changes. Per the biological insurance hypothesis, assemblages with more variation in function (*e*.*g*. functional dispersion, Laliberté and Legendre 2010) can better respond and maintain function as climate changes. The CWM of individual functional traits (Lavorel et al. 2008) provides insight into the overall functional strategy individuals in an assemblage utilize. Life history strategies potentially affect the relationships between these metrics and microenvironments. Additionally, *SES*_*MNTD*_, calculated from the combined functional and phylogenetic distances between the most closely related pairs of species (Cadotte et al. 2013), shows whether phylogeny represents differences between species that were not accounted for in the traits we measured. Color coding of the functional metric text used throughout the figures. (b) We assessed these relationships at twenty-six 1-m^2^ plots along the Right Hand Fork of the Logan River in northeast Utah (Simpson and Pearse 2021). One plot was excluded from the final analysis because the temperature sensor was removed by wildlife disturbance (in white). Background grayscale shows elevation based on a five-meter digital elevation model in meters (Utah Automated Geographic Reference Center 2007).

## 2 Materials & Methods

We assessed how the functional strategies and insurance potential of assemblages change across microenvironments at twenty-six 1-m^2^ fractally-arranged vegetation plots in a longterm study site along the Right Hand Fork of the Logan River in Cache National Forest, UT (41°46’12”N, 111°35’30”W; Figure 1). At this site, the spatial arrangement of these plots and sampling intensity effectively captured environmental variation and responses across the landscape (Simpson and Pearse 2021). We assessed the relationship between plant functions and microenvironments in a spatial context, *i*.*e*. we do not intend to assess the direct temporal response of function to the environment. Additionally, we looked at whether information about life history strategies and phylogenetic differences changed the understanding of ecological differences we assessed from functional diversity. Data processing and analyses were performed in R (v. 4.2.1; R Core Team 2022) and all software packages *in italics* below are R packages, unless otherwise noted. All data collected and code to reproduce analyses will be openly released.

### 2.1 Vegetation cover assessment & functional trait collection and processing

We measured the total canopy cover of vascular plant species in each 1-m^2^ plot during June–July 2018. We included cover from species rooted outside of the plot because this best represents the total functionality of the assemblage for abundance-weighted functional diversity measures. To standardize cover assessment, we used a quadrat divided into four 0.25-m^2^ quadrants and assessed percent cover with a 10 × 10 grid of 0.025-m^2^ grid cells. We identified plants using local herbarium resources and field guides and standardized taxa names using The World Flora Online (*<*http://www.worldfloraonline.org/*>*).

We collected functional traits based on their representation of the two main axes of variation in aboveground plant traits at both the species (Díaz et al. 2016) and assemblage (Bruelheide et al. 2018) levels. Plant size (the mean and maximum height) and leaf traits [specific leaf area (SLA) and leaf area (LA)] quantify contrasting functional strategies a plant uses to access to light and integrate resources, via competition or facilitation with neighboring individuals (Reich 2014). We also chose these traits because they both respond to environmental conditions and affect ecosystem functions (Lavorel and Garnier 2002). To focus on the functional consequences of potentially losing response diversity, we assumed that current variation in functional trait strategies across environment represents a unified functional strategy of response and effect traits, without directly measuring ecosystem function (Reich 2014; Suding et al. 2008; Elmqvist et al. 2003). All functional trait data were taken in or near the twenty-six plots during June–July 2018 and June–July 2019 and were collected, processed, and analyzed following Perez-Harguindeguy et al. (2016). We focused on interspecific variation in all traits.

#### Height traits

Globally, plant height represents the overall strategy of how a species lives (*e*.*g*., its lifespan), grows (*e*.*g*., time to maturity), and reproduces (*e*.*g*., seed mass and the number of seeds it produces; Díaz et al. 2016; Moles et al. 2009). In cold, dry places, like the overall climate at the fieldsite, a wide range of height strategies typically succeed compared to warm, wet environments, where tall species dominate. Because many of the species in the plots are graminoids and forbs, which can be very variable in height, we aimed to measure the height (cm) of up to twenty-five randomly selected individuals of each species within each 1m^2^ plot. If there were less than ten individuals in a plot, we continued measuring individuals from within ten meters of the plot. We measured from the ground to the top of the main photosynthetic tissue, not including inflorescences, seeds, or fruits if those extended beyond the tallest leaves. Drooping foliage was measured as-is to assess the general canopy height of the plant. Across all plots, for each species, we calculated one measure of average plant height and one measure of the maximum plant height achieved by that species. Calculating both height measures allowed us to look at an average measure of how height responds to environmental conditions across the site (mean height) compared to the maximum height that species achieved in all of the plots.

#### Leaf traits

We calculated the specific leaf area (SLA), the total fresh leaf area (LA; mm^2^) divided by its oven-dry mass (g), to quantify the resource acquisition strategy of each species. We aimed to collect at least five leaves from five individuals for each species within twenty meters of each plot. We adjusted the number of leaves based on size; from three leaves for large-leaved species to twenty leaves for small-leaved species. We collected leaves from each individual randomly, and when possible, chose fully-developed sun leaves that were undamaged by herbivory or pathogens. We placed the leaves from each individual in a sealed, plastic bag, to retain their moisture, and kept them flat using cardboard that was tied together for transport back to the lab. The same day, we scanned the leaves with a high-resolution flatbed scanner. Then, we dried them at 70 °C for 72 hours and weighed them to determine their oven-dried leaf mass (g).

We used an automated, threshold-based pipeline (‘stalkless’ Pearse et al. 2018) to calculate the leaves’ surface areas (mm^2^). This workflow relied on thresholding the contrast between dark and light pixels in an image to separate the leaves, or darker areas of the image, from the lighter background. As a baseline, we set the threshold to the mean intensity of each scan plus two times the standard deviation of each scan’s intensity. The program identified all regions of the scan greater than the threshold as leaves and calculated LA by counting the pixels in all of the regions larger than the mean region size plus two standard deviations as processed LA. We checked all of the processed images from the scans and adjusted the threshold to capture the correct shape of each leaf. To focus on interspecific variation in the leaf traits and avoid poor scans, we chose the best sample of leaves from a species, if multiple samples were collected. To choose the best sample, we prioritized fully-developed, undamaged leaves, followed by those collected in 2019 when we used a scanner that produced more precise images, and finally, all else equal, focused on samples taken from environments where the species was relatively abundant and the topography was most consistent with ‘average’ topography at the site.

### 2.2 Quantifying microenvironment

#### Near-surface soil temperature

Local seasonal temperature variation directly affects both ecosystem and individual plant functions and relates to other important microclimate conditions, like the consistency of snow cover (Lembrechts et al. 2020). To measure near-surface soil temperature, we buried a HOBO 8K Pendant®Temperature/Alarm Data Logger (UA-001-08) in a ten-centimeter deep hole at each of the plots. We anchored the logger into the sides of the hole with metal landscaping pins attached to the logger with zip ties. Then, we covered the logger with soil and rocks, to match the surrounding landscape and protect the sensor from disturbance by wildlife. Each logger was set to record the temperature every 90 minutes and start logging at midnight the following day using HOBOware software (*<*https://www.onsetcomp.com/hoboware-free-download/*>*). We downloaded temperature data during September 2018 and September 2019 to get a full year of temperature data. One sensor was lost in 2018, because of substantial wildlife disturbance (it appeared to be pulled out by a grazer or dug up by a rodent, Figure 1, in white), resulting in temperature data at twenty-five plots. To summarize inter-annual temperature variables, we converted the temperature readings to °Celcius using *weathermetrics* (Anderson et al. 2013) and manipulated the date and time to be able to assess the first and last month and day temperature readings recorded at each plot using *lubridate* (Grolemund and Wickham 2011) and *dplyr* (Wickham et al. 2021). Then, we subset the time frame to a year of temperature data, from September 28, 2017, at 00:00 (Mountain Standard Time, MST) to September 28, 2018, at 00:00 MST, and calculated the annual mean, maximum, minimum, and standard deviation in temperature at each plot.

Soil texture. We used the hydrometer method [following procedure and calculations in Ashworth et al. (2001), based on Bouyoucos (1927)] to assess soil texture from soil samples collected at the twenty-six core plots in mid-summer 2018. At the same position about half a meter from each plot, we removed the organic matter and collected soil from a ten-centimeter deep by eight-centimeter wide hole and transferred it back to the lab to be aired dried for further processing. We physically broke up the soil clumps so that particles would disperse by sieving the soils to two millimeters and further grinding them with a mortar and pestle. We chemically dispersed the soil using a 50 g/L sodium hexametaphospate solution and finished dispersing the solution by inverting the cylinder several times. We took hydrometer measurements at forty seconds and two hours to determine the amount of sand, silt, and clay in each soil sample. Finally, we used the package *soiltexture* to classify these percentages into soil classes based on the USDA soil texture triangle (Moeys 2018).

### 2.3 Statistical Analysis

We aimed to quantify how biological variation, as measured by functional diversity and phylogeny, relates to environmental heterogeneity, as measured by microenvironment (see the framework in Figure 1). First, we looked at whether differences in topography (aspect, elevation, and slope) predict differences in microenvironment (near-surface soil temperature and soil texture). Then, we assessed whether topography-predicted microenvironmental conditions predicted functional diversity — mean and variation. We analyzed whether these relationships differ depending on plant life history strategy, *i*.*e*. whether a plant is an annual/biennial, herbaceous perennial, or woody perennial because these groupings tend to have more similar traits, compared to all plant species. Finally, we incorporated information about ecological differences from both phylogenetic and functional differences to determine if phylogeny quantifies differences between species that were not represented by the traits we measured.

Microenvironment-topography relationships. We looked at how microenvironment varies across topography to quantify how near-surface soil temperature and soil texture spatially vary across the fieldsite. Since our analysis is based on twenty-five plots, we aimed to isolate relationships between one microenvironmental variable and one topographic variable to properly estimate coefficients. Soil texture can affect soil temperature (Akter et al. 2015), so we assessed whether each temperature variable correlated with the components of soil texture (percentage of sand, silt, and clay) by calculating the correlation coefficient, Pearson’s *r*. Then, we looked at how each microenvironmental variable — the mean, standard deviation, maximum, and minimum temperature and amount of sand, silt, and clay — correlated with three topographic variables — aspect, elevation, and slope, because drainage patterns can affect soil particle distribution (Brown et al. 2004).

Functional diversity-microenvironment relationships. We quantified the functional strategies in each assemblage using functional dispersion (FDis; Laliberté and Legendre 2010) and the community weighted mean (CWM, Lavorel et al. 2008) of each of four traits — specific leaf area (SLA), leaf area (LA), mean height and maximum height. We calculated FDis, the mean distance of all species’ traits to the weighted centroid of the assemblage in multivariate trait space, using *FD::*dbFD (Laliberté et al. 2014). We calculated the abundance-weighted version of this metric to assess how the prevalence of species contributes to that assemblage’s potential to maintain function in the future. First, we generated a speciesby-species distance matrix from (weighted) functional traits using the Gower (dis)similarity coefficient (Gower 1971). Because of large differences in the units of the different traits we measured, we standardized each trait to have a mean zero and a unit variance. Then, we performed a principal coordinate analysis (PCoA) on this uncorrected species-species distance matrix to generate PCoA axes that were used as ‘traits’; all four PCoA axes were maintained. To verify that the closely related leaf and height traits were not over-inflating FDis, we also calculated FDis with just two traits — maximum height and SLA. Again, we used *FD::dbFD* to calculate the abundance-weighted CWM of the four traits for each assemblage (Lavorel et al. 2008). This provided more detailed information about the functional composition of each assemblage; in the case of the traits we chose, about overall plant size and leaf economic strategies.

To ensure we did not over-fit our data, we needed to be selective in choosing environmental predictors of functional diversity at our twenty-five plots. So, we assessed which temperature and texture variable each functional diversity metric temperature correlated most strongly with using Pearson’s *r*. This resulted in one temperature and one texture explanatory variable in each additive linear model of functional diversity across microenvironment. We modeled the CWM of leaf traits, LA and SLA, and FDis as a function of mean soil temperature and the amount of clay in the soil. We modeled the CWM of height traits, mean and maximum, as a function of the interannual variation [standard deviation (SD)] in soil temperature and the amount of sand in the soil. FDis calculated with two traits, maximum height and SLA, was modeled as a function of mean soil temperature and the amount of sand in the soil. We logged all functional diversity metrics to improve normality and used ANOVA to test whether both, either, or none of the environmental variables best predicted our diversity metrics.

#### Effect of life history strategies on functional diversity-environment relationships

We determined whether each species assessed was a woody perennial, herbaceous perennial, or annual/biennial using a local flora (Shaw et al. 1989), and subset the species in each assemblage into these groups. Then, we calculated all five functional metrics for each of these subsets, as described above. Across the entire site, we calculated the overall FDis and CWM of LA, SLA, and maximum and mean height for each life-history strategy. Then, at the assemblage-level, we used model-averaging (using *MuMIn::dredge*; Bartoń 2022) to statistically test whether each life-history strategy’s diversity and changes across environmental gradients, differed from one another. All predictor variables were z-transformed to make their resulting coefficients a measure of the relative importance of each explanatory variable (Grueber et al. 2011).

Effect of phylogenetic differences on understanding ecological differences. Species’ phylogenetic relationships may represent ecological differences not captured by the functional differences we measured. We assessed whether or not phylogeny added information about ecological differences using the mean nearest taxon distance (*SES*_*MNTD*_). This metric averages the distance between nearest neighbors for all species in the assemblage and compares that to a randomized, null assemblage drawn from the wider source pool (*n* = 999, Pearse et al. 2015; Kembel et al. 2010; Kembel 2009; Webb 2000). We calculated *SES*_*MNTD*_ from the combined functional and phylogenetic distances between the most closely related pairs of species using the phylogenetic weighting parameter *a*. This ‘traitgram’ approach (Cadotte et al. 2013) means that, when *a* = 0, *SES*_*MNTD*_ reflects only functional differences, while when *a* = 1, *SES*_*MNTD*_ is generated from a distance matrix of only phylogenetic differences. Importantly, when *a* is intermediate between the two, it reflects both phylogeny and traits (when *a* = 0.5 it reflects both equally), and so the relative contributions of both can be assessed.

For phylogenetic distances, we used the phylogenetic tree for vascular land plants from Zanne et al. (2014) and added missing species using *pez::congeneric*.*merge* (Pearse et al. 2015). We set the phylogenetic weighting parameter to calculate abundance-weighted *SES*_*MNTD*_ eleven times (*a* = 0, 0.1, 0.2, …0.9, 1), using *pez::*.*ses*.*mntd*, to see whether phylogenetic and functional information are revealing related, or complementary, information about our system. Finally, we looked at both how *SES*_*MNTD*_ varied overall, and how the relationship between

*SES*_*MNTD*_ and microenvironment changed, as the amount of difference from functional and phylogenetic information varied.

## 3 Results

### 3.1 Microenvironment-topography relationships

There were no significant correlations between the two types of microenvironment variables, soil temperature and texture (Supplementary information). Near-surface soil temperature and soil texture varied across different elements of topography, soil temperature across aspect and soil texture across elevation (Figure 2). Overall, near-surface soil temperature variables —the mean, maximum, and standard deviation in temperature — were higher on south-facing slopes than north-facing ones. On average, south-facing slopes had a 6.7 °C warmer mean temperature, 21 °C warmer maximum temperature, and 4.6 °C more variation in temperature than north-facing slopes. The average minimum temperature at each plot (−1.7 ^+^/-0.39 °C) did not vary across aspect.

**Figure 2:**
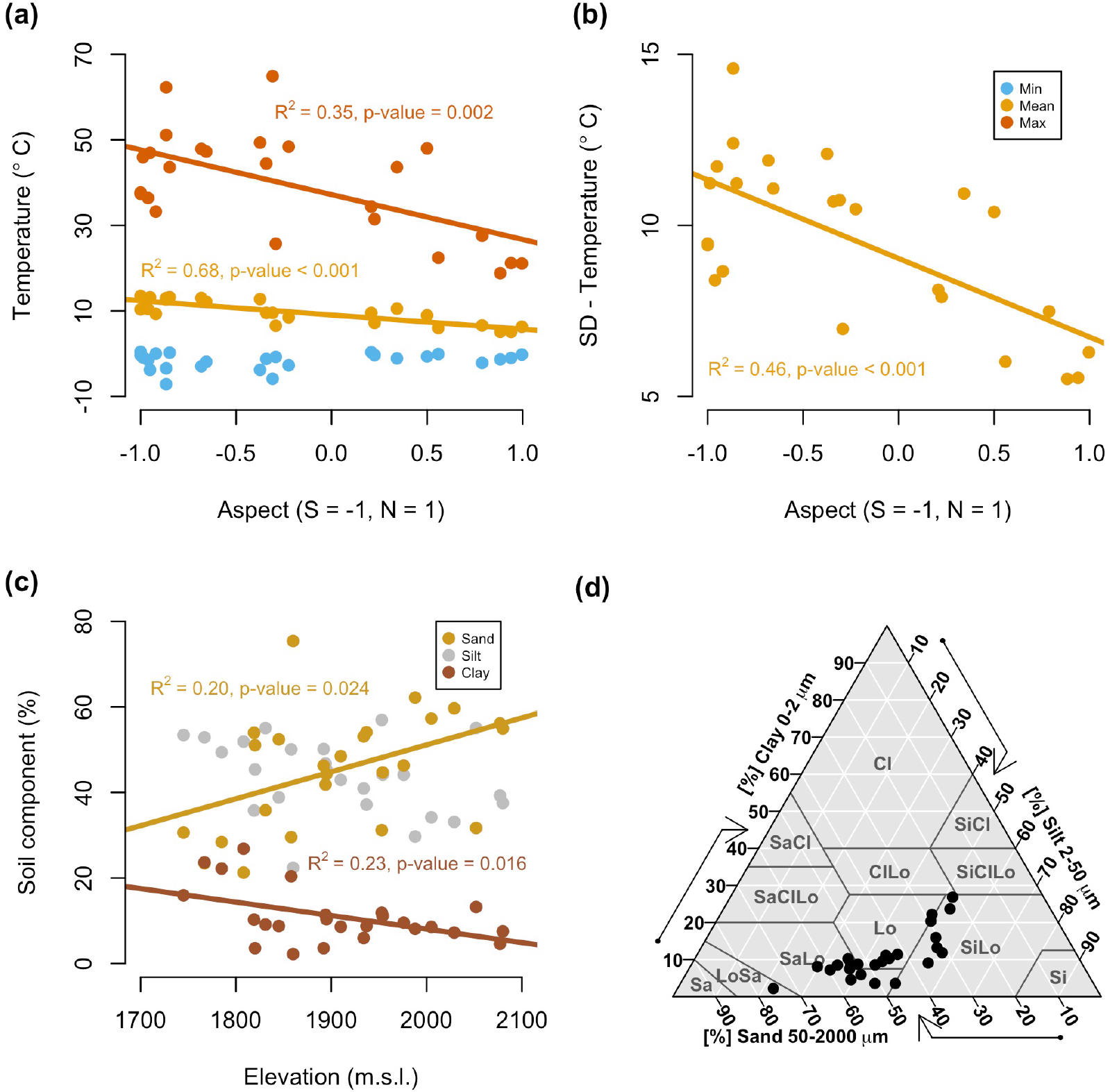
Near-surface soil temperature and soil texture vary across topography at Right Hand Fork. (a) The mean (yellow, slope = -3.33, *F*_1,23_ = 49.59) and maximum (red, slope = -10.49, *F*_1,23_ = 12.61) near-surface soil temperatures are significantly higher in more south- than north-facing plots. The minimum temperature does not significantly change across aspect (light blue, average = -1.69 °C). (b) The variance in temperature at each plot also significantly decreased from south- to north-facing plots (slope = -2.30, *F*_1,23_ = 19.47). (c) The percent of sand and clay vary inversely across elevation with lower amounts of sand (yellow, slope = 0.063, *F*_1,23_ = 5.85) and higher amounts of clay (brick red, slope = -0.032, *F*_1,23_ = 6.79) at lower elevations. The amount of silt in the soil did not significantly change across aspect (gray, average = 43.70 %). (d) All of the soils at Right Hand Fork are loams with lower amounts of clay (0 - 30 %), moderate amounts of silt (20 - 60 %), and moderate to high amounts of sand (20 - 80%).

The soil texture at all plots was loamy, including nine sandy loams, eight silty loams, seven loams, and one loamy sand (Figure 2). The texture of these soils are all low in clay (10-30%), with moderate amounts of silt (20-60%) and the highest range in the amount of sand (20-80%). Both the percentage of sand and clay significantly varied across elevation. Elevation predicted lower amounts of sand (about 35%) and higher amounts of clay (about 15%) at the lowest elevation plots (1745 m.s.l.) and higher amounts of sand (about 55%) and lower amounts of clay (about 5%) at the highest elevation plots (2080 m.s.l, Figure 2). The amount of silt in the soil did not significantly vary across elevation (average = 43.7%).

### 3.2 Functional diversity-microenvironment relationships

We obtained four traits for 84 out of the total 100 species identified. We scanned and weighed a total of 7,213 leaves and obtained LA from the stalkless pipeline for 6,613 of those leaves. Prioritizing the best sample of leaves for each species resulted in 3,454 leaves that were used to generate the leaf traits in the analysis presented here. We measured the height of 1,831 individuals, all of which were used to calculate the mean and maximum height variables. Microenvironments with the lowest mean soil temperatures, which tended to be on north-facing slopes, supported assemblages with larger leaves, more acquisitive strategies, and more functional dispersion (Figure 3, Supplementary information). Conversely, plots with less inter-annual variation in soil temperature, also found on north-facing slopes, predicted taller assemblages, whether measured as the mean or maximum. When FDis was calculated with two traits, the relationship between functional dispersion and mean temperature was very similar [FDis (4 traits) slope = -0.316, FDis (2 traits) slope = -0.321, Supplementary information]. The relationship between height and the inter-annual variation in temperature was also similar whether it was calculated as the maximum height a species achieved or the mean height of the species across the site (maximum height slope = -0.218, mean height slope = -0.233, Supplementary information).

**Figure 3:**
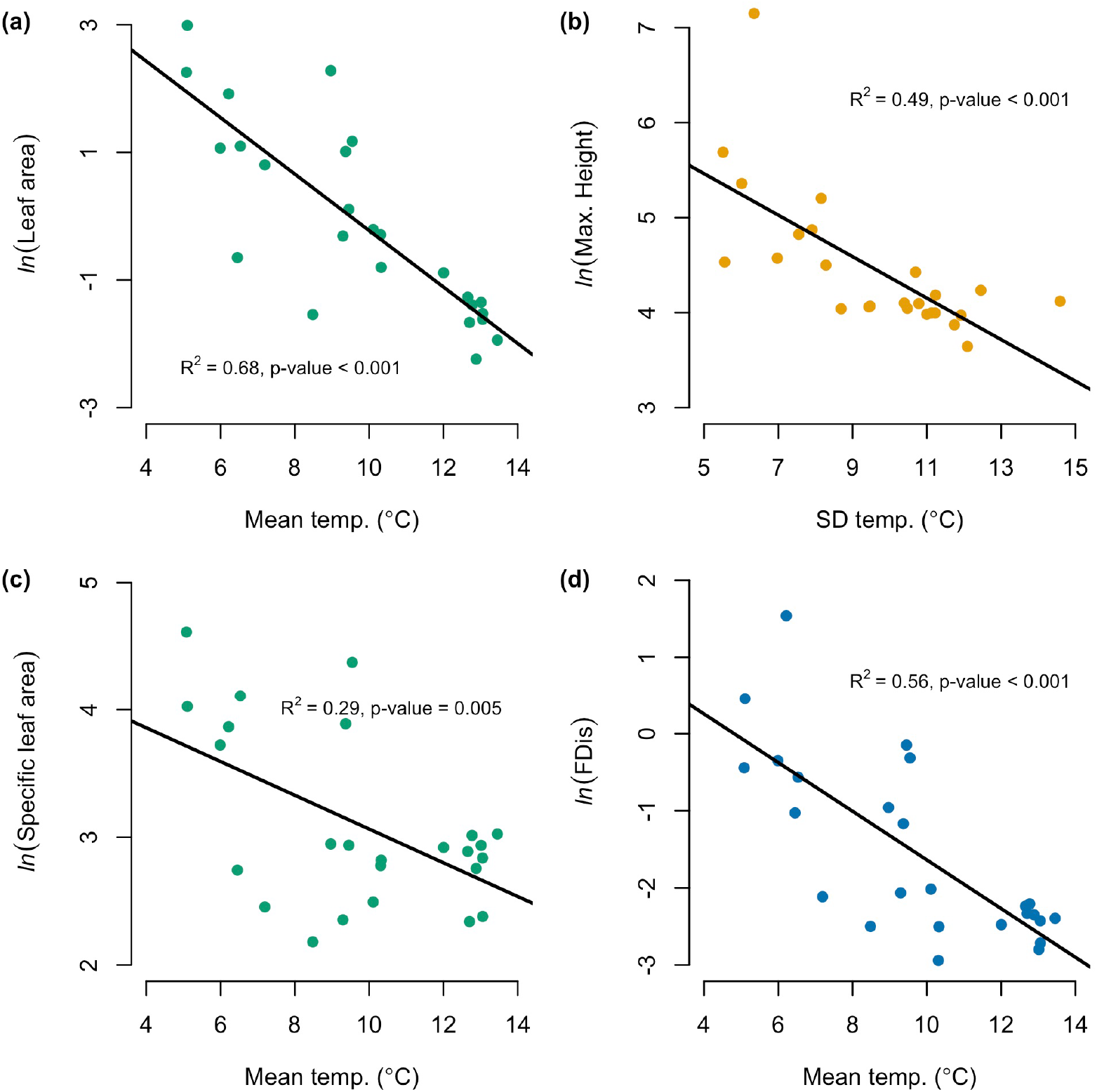
Increases in mean soil temperature predict a decrease in the CWM of leaf traits and functional dispersion, while an increase soil temperature variation predicts a decrease in the CWM of maximum height. (a) Plots with cooler mean soil temperatures support larger leaves [higher logged CWM of LA (mm^2^), slope = -0.442, *F*_1,23_ = 49.97], (c) leaves with more acquisitive leaf economic strategies [higher logged CWM of SLA (mm^2^ g^-1^), slope = -0.132, *F*_1,23_ = 9.426], and assemblages with more variance in functional strategies [*i*.*e*. more biological insurance; higher logged FDis, slope = -0.316, *F*_1,23_ = 29.54]. (b) Plots with less variation in soil temperature support taller assemblages [higher logged CWM of maximum height (cm), slope = -0.218, *F*_1,23_ = 21.85]. Color coding described in Figure 1.

### 3.3 Effect of life history strategies on functional diversity-environment relationships

Site-wide, functional diversity varied across the life history strategies — woody or herbaceous perennial, and annual/biennial. Herbaceous perennials made up the largest group of species (54/84) and had the most acquisitive leaves and largest leaf area (Supplementary information). Annuals and biennials made up the next largest group of species (17/84) and had the smallest functional dispersion (therefore, lowest biological insurance), least acquisitive and smallest leaves, and shortest height. Woody perennials had more than four times the functional dispersion of herbaceous perennials (the largest amount of biological insurance) and were the tallest group.

The life history strategy of species affected the relationship between functional diversity and microenvironment. Herbaceous and woody perennials drove the interaction between leaf area and mean temperature and functional dispersion and mean temperature (Figure 4). Both groups had larger leaves and higher dispersion when the mean temperature was lower.

**Figure 4:**
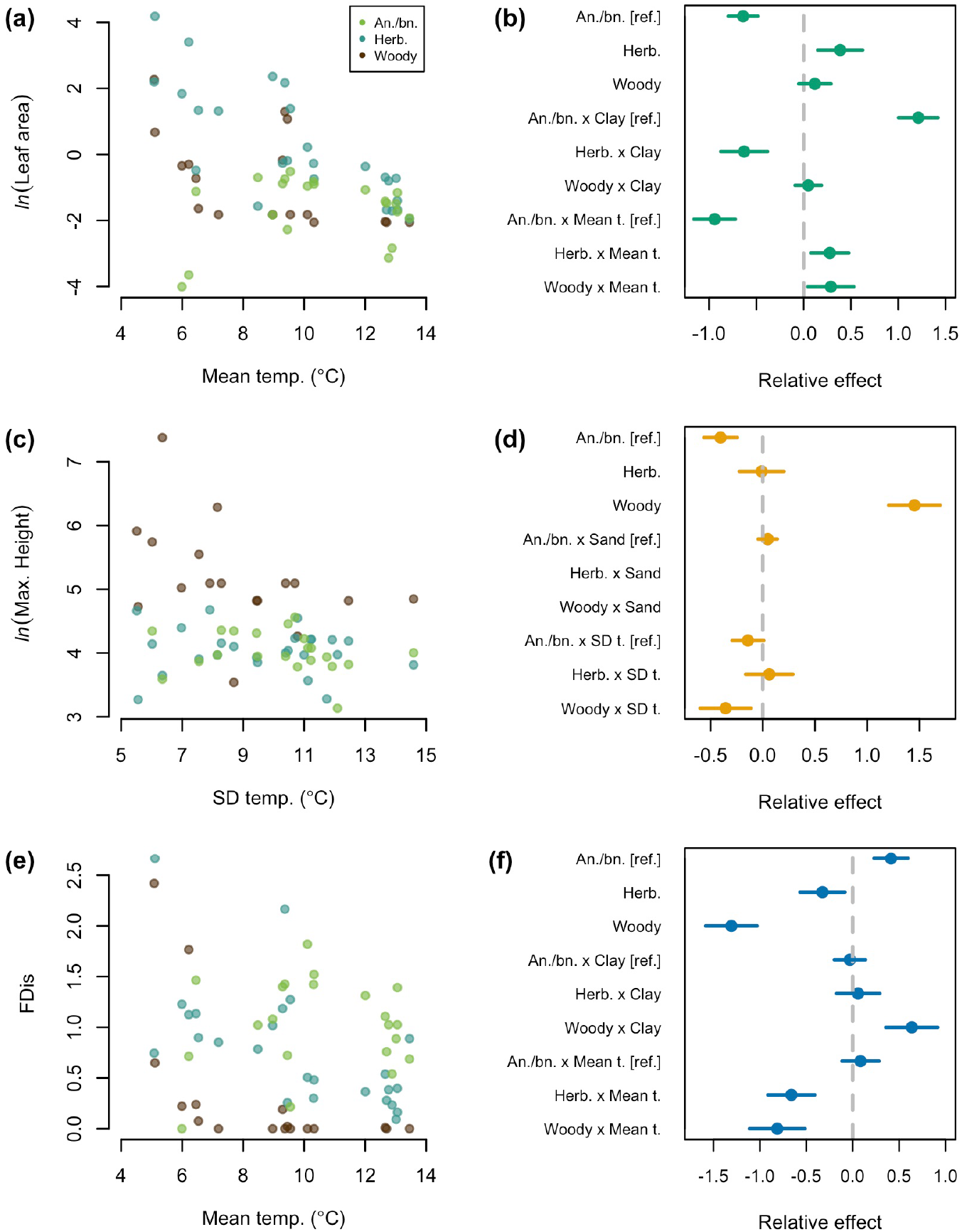
Life history strategies affect the relationship between functional diversity and microenvironment. Plots in the left column show how the logged CWM of (a) leaf area, (c) maximum height, and (e) [unlogged] functional dispersion vary across the temperature variable they were most correlated with when subset by life history strategy —annuals/biennials (green), herbaceous perennials (blue), and woody perennials (brown). Plots in the right column show the relative effect of each explanatory variable in models that look at how these life history strategies affect the relationship between each functional metric (in the left column) and both the soil temperature and texture variable most correlated with that functional metric. Coefficient values are reference contrasts from those labeled as such. Values further from zero indicate that a variable or interaction between variables has a greater effect on a functional metric. For example, when a life history strategy interacts with one of the microenvironment variables and has a large relative effect, as woody and herbaceous perennials interact with mean temperature in (b), plants with these life history strategies have a larger impact on the relationship between that functional metric and microenvironmental variable.

Woody perennials had a small effect on the relationship between functional dispersion and the amount of clay in the soil; functional dispersion was higher when the amount of clay in the soil was higher within this group. Herbaceous perennials had the biggest effect on the relationship between specific leaf area and mean soil temperature (Supplementary information); species in this life history group had more acquisitive leaves when mean temperature was lower. The CWM of mean and maximum height was mostly driven by the presence of woody perennials, *i*.*e*. assemblages containing woody perennials were taller overall (Figure 4, Supplementary information). Additionally, when variation in temperature was lower, woody perennials achieved taller maximum heights. Overall, we did not detect a change in the functional diversity of annuals and biennials across microenvironment.

### 3.4 Effect of phylogenetic differences on understanding ecological differences

Broadly, functional and phylogenetic differences between species contributed similar information about the ecological differences between species at Right Hand Fork. Across the whole site, *SES*_*MNTD*_ was highest when calculated only from functional differences (*a* = 0), second highest when only calculated from phylogenetic differences (*a* = 1), and lowest when calculated from about half functional and half phylogenetic difference (*a* = 0.5, Figure 5).

**Figure 5:**
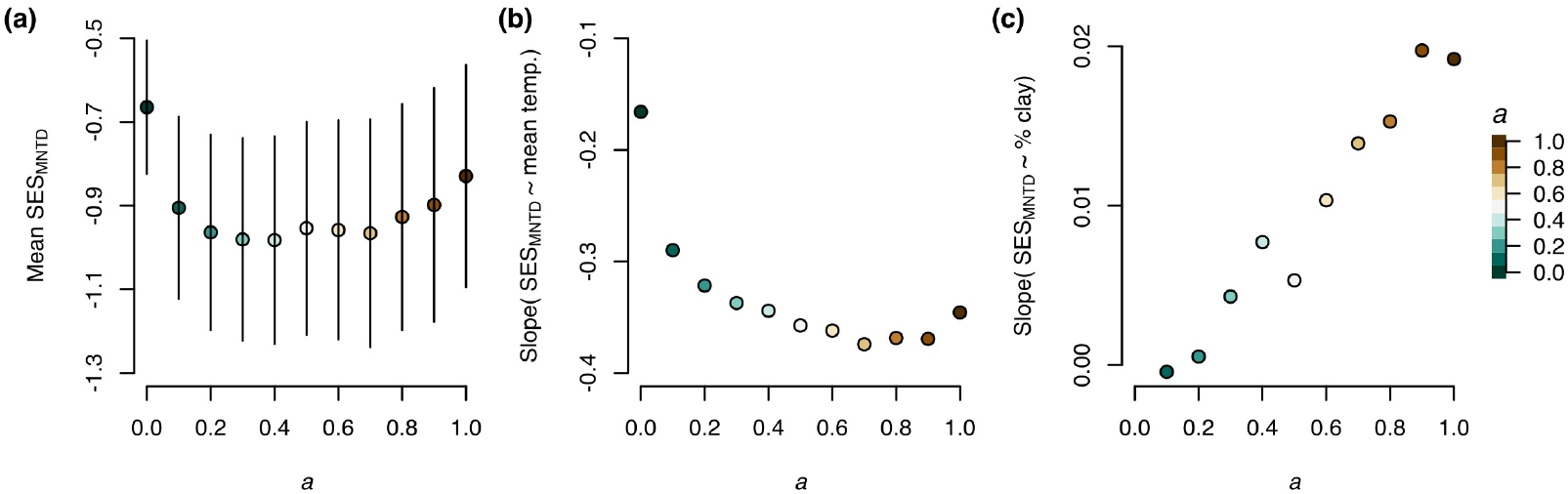
Adding phylogenetic information about ecological differences strengthens the relationship between *SES*_*MNTD*_ and microenvironment. The phylogenetic weighting parameter (*a*) changes the amount of information about ecological differences used to calculate *SES*_*MNTD*_ from all functional differences (teal, *a* = 0) to all phylogenetic differences (brown, *a* = 1). (a) Overall, including phylogenetic differences does not provide significantly different information about ecological differences than just using functional differences. (b) The relationship between *SES*_*MNTD*_ and mean temperature is stronger when *SES*_*MNTD*_ is calculated from about half functional and half phylogenetic differences or more phylogenetic than functional differences. (c) Similarly, the relationship between and the amount of clay in the soil is stronger when just calculated from phylogenetic differences.

However, none of the values of *SES*_*MNTD*_ we calculated across the phylogenetic weighting parameter (*a*) were significantly different. That said, the relationship between *SES*_*MNTD*_ and environment (both soil temperature and texture) significantly changed as the value of *a* changed. The relationship between *SES*_*MNTD*_ and mean temperature was least strong when *SES*_*MNTD*_ was calculated only from functional metrics (slope = -0.17) and most strong when calculated from about half functional and half phylogenetic difference or a greater amount of phylogenetic difference (*a >* 0.5, slope = -0.36). Similarly, the relationship between *SES*_*MNTD*_ and the amount of clay in the soil became more strong, albeit subtly, as phylogenetic differences were included (slope = -0.01 to slope = 0.02).

## 4 Discussion

The relationship between functional diversity and microenvironment quantifies where and how assemblages might maintain function in topographically complex, mountainous terrain in northeastern Utah. Broadly, we found a shift in overall functional strategy and biological insurance potential across soil temperature gradients across aspect. Herbaceous (and to some degree, woody) perennials had the strongest effect on these relationships. Integrating information about the functional and phylogenetic differences in a dispersion metric (*SES*_*MNTD*_) indicated that phylogeny represents ecological differences between species that we did not measure.

Here, we discuss the implications of changes in near-surface soil temperature for overall plant survival and growth across this landscape. Then, we describe how shifts in functional diversity — driven mainly by the CWM of leaf traits and dispersion of herbaceous perennials — point to an environmental filter across aspect. This trend highlights a contrasting potential for persistence, and management opportunities, on northvs. south-facing aspects. Finally, we discuss how the addition of phylogenetic differences enhances our understanding of ecological differences in these assemblages. Throughout, we highlight how assessing functional diversity and microenvironment, within the framework we describe, can inform management actions focused on maintaining plant assemblage functions and ecosystem services.

### 4.1 Topography shapes microenvironment

At the local spatial scales we sampled at (1900 m extent and smaller), aspect-driven topographic complexity moderates a site-wide climate regime. Across aspect we measured a 6.4 °C mean temperature change. Mean temperature correlated most strongly with aspect (Figure 2, R^2^ = 69%). Other mountainous environments throughout the northern hemisphere likely show similar trends, because south-facing aspects receive more solar radiation which increases soil temperature and evapotranspiration (Jackson 1967). However, we did not find differences in soil texture across aspect, indicating that this topographic gradient does not influence water regimes (Brown et al. 2004). A global study of temperate grass-/shrub-lands with similar mean annual temperatures (MAT = 7.23 °C, compared to a MAT = 8.4 °C at Right Hand Fork) also found that topography better predicted environmental heterogeneity in temperature than precipitation (Jiang et al. 2017). The strong relationship between mean temperature and aspect shows that small changes in near-surface soil temperature likely moderate resource availability, like soil moisture, that drive differences in functional strategies. If mean temperatures increase across this site, the variance and maximum temperatures could become more correlated with topography than the mean temperature (Lewis and King 2017). This would indicate a shift to a more variable climate regime associated with topographic complexity.

Near-surface soil texture varied across elevation, but not aspect. Soils at higher elevations were coarser with high sand (55%) and low clay content (15%) and overall, soils at Right Hand Fork are characterized by high sand content (*>*20%, Figure 2). Per the inverse-soil texture effect, coarse-textured soils decrease bare soil evaporation by allowing water to infiltrate to deeper soil layers (Noy-Meir 1973; Walter et al. 1973). This leads to greater (overall) water availability in deeper layers that supports higher plant productivity and more woody plant growth in arid climates (Renne et al. 2019; Pennington et al. 2017; Dodd and Lauenroth 1997; Sala et al. 1997). However, these soils hold less water and nutrients at the surface (Austin et al. 2004). Per this effect, the coarse composition of the soil at Right Hand Fork may support taller plant assemblages than would be present if the soil was finer. To verify this insight we need to measure soil depth, soil moisture content, and texture at multiple depths in the soil column because this insight only applies to deeper soils. We could also measure the root traits of herbaceous perennials; if they exhibit more conservative strategies and deeper rooting this would provide support for the inverse-soil texture effect.

### 4.2 Microenvironment predicts functionally-distinct assemblages

An increase in the mean and interannual variation of temperature across northto southfacing aspects supported functionally distinct assemblages (Figure 3). Plots with cooler and less variable temperatures contained assemblages that were taller, more functionally dispersed, and had more acquisitive leaf economic strategies. Conversely, plots with hotter and more variable temperatures supported assemblages that were shorter, less functionally dispersed, and had more conservative leaf economic strategies. This trend indicates that environmental filtering dominates on south-facing slopes, especially for herbaceous and woody perennials. Limited resources (*e*.*g*. less water) and environmental stress (*e*.*g*. higher temperatures) likely act as this filter, but biotic interactions may also contribute (Kraft et al. 2015; Cadotte and Tucker 2017; Mayfield and Levine 2010; Cornwell et al. 2006). Annuals and biennials appear to avoid this filter and use similar functional strategies across this temperature gradient. Soil texture did not predict functional differences in the aboveground traits we measured. This aligns with global studies where temperature predicts plant traits more strongly than precipitation (Moles et al. 2014). However, measuring traits more directly related to water acquisition (*i*.*e*. root traits) may provide better insight into how microenvironmental variation in water regimes influences assemblage function.

Abiotic and biotic factors operate across spatial scales to shape traits observed at small spatial scales; however, similar conditions can support assemblages with vastly different community-weighted trait values (Bruelheide et al. 2018). For example, in high-elevation Colorado mountains near the UT fieldsite we worked in, shorter plants with smaller leaves and more resource-conservative strategies were also found in locations with higher mean and variance in temperature (Stark et al. 2017). However, higher temperatures likely only constrain plant height and result in leaves with more conservative resource acquisition strategies in places where overall water availability is limited. In a study in the Arctic that compared locations with more and less moisture, warmer summer temperatures resulted in taller, more resource-acquisitive, aboveground plant traits, but only at the wet locations (Bjorkman et al. 2018). Notably, this pattern was mostly driven by species turnover, rather than intraspecific changes in functional traits.

Right Hand Fork spans active grazing allotments (USDA Forest Service 2022), where consistent vegetation production, especially by herbaceous perennials, supports people’s livelihoods. Additionally, herbaceous perennials drive shifts in the CWM of leaf traits and functional dispersion across mean temperature. As a result, it will be important to continue monitoring to understand how changes in the climate regime, especially soil temperature, might affect forage production. Environmental heterogeneity, especially aspect, has the potential to provide spatial insurance for these assemblages. Often, species are expected to move up in elevation and poleward in latitude in response to warming temperatures (Rubenstein et al. 2020). However, especially in arid, mountainous environments, aspect can buffer the overall effect of warming temperatures, supporting vegetation assemblages in ways that are similar to what we observed (Albrich et al. 2020; Yang et al. 2020). In response to increasing temperatures, species can also move across aspects and up in elevation, as seen with salamanders and lizards (Feldmeier et al. 2020).

Critically, even if cooler, north-facing slopes do not provide a buffer against increasing temperatures, vegetation often responds differently on northvs. south-facing slopes, which makes this an important topographic element to include in range shift studies and management plans (Ackerly et al. 2020; Elliott and Cowell 2015). Heterogeneity in soil resources also supports more diversity in functional traits, broadly speaking (Price et al. 2017). Future studies looking at how species adapt and move in response to changing environmental conditions should include responses to microenvironmental conditions, like aspect, not just broad climate regimes. This will facilitate a better understanding of subtle species shifts to, or maintenance within, more favorable environments to the preservation of the overall functionality of ecosystems (Fridley et al. 2011). Measuring dispersal traits, and monitoring how assemblage dispersion and forage production change over time, would further identify the potential for aspect to provide spatial insurance across this landscape.

### 4.3 Phylogenetic difference informs ecological difference

The overall value of *SES*_*MNTD*_, a measure of dispersion, was similar whether we calculated it from solely functional or phylogenetic information (Figure 5). However, the relationship between dispersion and environment was strongest when information from both phylogeny and functional traits was included. This example of how phylogenetic differences can support functional differences [see also de Bello et al. (2017)] adds to the ongoing debate about how many axes of functional variation are needed to understand diversity-environment relationships (Mouillot et al. 2021; Laughlin 2014). While the traits we measured provide information about how assemblages change across environment, the changes we detect are, if anything, conservative. Given the strength of these conservative trends, we suggest that the most effective use of management resources to understand the potential for this landscape to keep producing forage would be to track the productivity (and potentially variance in dispersal and water-acquisition traits) across aspect-driven temperature gradients.

### 4.4 Conclusion

Overall, we found evidence for differences in assemblages’ insurance potential and trait strategies driven by a 6.4°C shift in mean near-surface soil temperature across aspect, which likely affects water availability on these slopes. Assemblages on the warmer south-facing slopes showed less insurance potential (lower function dispersion) and had more resource-conservative leaf traits and shorter stature, consistent with adaptation to a harsher environment than north-facing slopes. The assemblage-level traits on north-facing slopes were consistent with cooler, less harsh environmental conditions. While we found taller species with more acquisitive leaf traits in these plots, they also had higher functional dispersion, indicating higher insurance potential on these slopes. Across the same temperature gradient, and differences in soil texture, adding information about phylogenetic difference to a metric of functional difference strengthened the relationship between ecological difference and environment. These results support the use of functional, and phylogenetic, diversity-environment relationships to understand assemblages’ current insurance potential and how it might respond to future change. We suggest that monitoring temporal trends in these relationships over time would provide more information about inter-annual variability in these relationships. For example, decreases in the FDis of assemblages over time might signal the increasing impact of an environmental filter, like higher temperatures, less precipitation, or land use intensification (Laliberté et al. 2010). We also advocate for more inclusion of aspect-differentiation in range-shift studies and vegetation management plans, because we found strong variation in assemblages’ insurance potential and related evolutionary history in response to this gradient.

## Supporting information

Supplement - data and analysis

## 5 Author Contributions

William D. Pearse and Elizabeth G. Simpson contributed equally to the study conceptualization, methodology, analysis, and manuscript preparation but EGS did the majority of data collection. Ian Fraser and Hillary Woolf provided considerable assistance with data collection.

## 6 Data Availability Statement

We intend to release all data and analysis code as part of the supplementary materials and intend to archive the data and code in a linked, public Github repository.

## 7 Acknowledgments

We thank the Department of Biology, Ecology Center, and Intermountain Herbarium of Utah State University for their support. We are grateful to M. Barkworth and M. Piep for helping us with plant identification, to E. James, S. Kinosian, A. Koontz, M. Stemkovsi, and B. Weedop for fieldwork assistance, to B. Waring for the soil texture protocols and lab equipment, and to the U.S Forest Service for permission to conduct this research in the Uintah-Wasatch-Cache National Forest. We appreciate feedback from and discussion with N. Huntly, K. Kettenring, R. Schaeffer, and B. Waring, which helped improve this manuscript.

