## Supplemental tables and figures for "Variation in near-surface soil temperature drives plant assemblage insurance potential"

### A Supplementary Information

Table 1: **Near-surface soil temperature does not significantly correlate with soil texture.** This table shows the correlation coefficients (Pearson's r) and p-values for each temperature variable with the three soil texture components.

| | Sand | Silt | Clay |
|---|---|---|---|
| Mean temp. | -0.15 | 0.03 | 0.27 |
| *p* value | 0.48 | 0.88 | 0.20 |
| SD temp. | -0.09 | -0.02 | 0.23 |
| *p* value | 0.66 | 0.92 | 0.28 |
| Max temp. | -0.13 | 0.03 | 0.23 |
| *p* value | 0.54 | 0.89 | 0.26 |
| Mean temp. | 0.24 | -0.10 | -0.37 |
| *p* value | 0.25 | 0.65 | 0.07 |

Table 2: **Near-surface soil temperature most strongly and significantly correlates with aspect, while soil texture components most strongly and significantly correlate with elevation.** This table shows the correlation coefficients (Pearson's $r$ and p-values for each temperature and texture variable in relation to three topographic variables—aspect, elevation, slope. The coefficient for the most strongly correlated topographic variable is bolded.
Here, * means that $p < 0.05$.

| | Aspect | Elevation | Slope |
|---|---|---|---|
| Mean temp. | **-0.827** | -0.025 | -0.250 |
| *p* value | <0.001* | 0.906 | 0.227 |
| SD temp. | **-0.677** | 0.134 | -0.322 |
| *p* value | <0.001* | 0.524 | 0.116 |
| Max. temp. | **-0.595** | 0.023 | -0.291 |
| *p* value | 0.002* | 0.914 | 0.159 |
| Min. temp. | 0.240 | 0.319 | 0.027 |
| *p* value | 0.248 | 0.120 | 0.896 |
| % Sand | 0.278 | **0.450** | -0.091 |
| *p* value | 0.178 | 0.024* | 0.666 |
| % Silt | -0.278 | -0.344 | -0.072 |
| *p* value | 0.179 | 0.092 | 0.733 |
| % Clay | -0.205 | **-0.477** | 0.291 |
| *p* value | 0.326 | 0.016* | 0.158 |

Table 3: **The CWM of leaf traits and functional dispersion significantly and most strongly correlates with mean soil temperature, while the CWM of height traits significantly and most strongly correlate with inter-annual variation in soil temperature**. Leaf traits and functional dispersion calculated from four traits correlated most strongly with the amount of clay in the soil while height traits and functional dispersion calculated from the CWM of SLA and maximum height correlated most strongly with the amount of sand in the soil. All functional metrics were logged to improve normality in resulting models. For each functional metric, this table shows the correlation coefficients (Pearson's $r$) and p-values between that metric and seven soil temperature and texture environmental variables. Since the temperature and texture variables exhibit co-linearity and we can only use two predictor variables in each multiplicative and additive linear model, we chose the temperature and texture variables that was were most correlated with each functional metric (bolded).
Here, * means that $p < 0.05$.

| | Mean temp. | SD temp. | Max. temp. | % Sand | % Clay |
|---|---|---|---|---|---|
| $ln$(CWM of LA) | **-0.825** | -0.695 | -0.536 | 0.059 | **-0.105** |
| $p$ value | <0.001* | <0.001* | 0.006* | 0.779 | 0.617 |
| $ln$(CWM of SLA) | **-0.539** | -0.485 | -0.381 | 0.026 | **-0.028** |
| $p$ value | 0.005* | 0.014* | 0.060 | 0.903 | 0.893 |
| $ln$(CWM of max. height) | -0.666 | **-0.698** | -0.683 | **-0.144** | 0.113 |
| $p$ value | <0.001* | <0.001* | <0.001* | 0.493 | 0.591 |
| $ln$(CWM of mean height) | -0.643 | **-0.679** | -0.666 | **-0.092** | 0.069 |
| $p$ value | 0.001* | <0.001* | <0.001* | 0.662 | 0.741 |
| $ln$(FDis - 4 tr.) | **-0.750** | -0.699 | -0.603 | <0.001 | **-0.033** |
| $p$ value | <0.001* | <0.001* | 0.001* | 0.998 | 0.877 |
| $ln$(FDis - 2 tr.) | **-0.748** | -0.709 | -0.614 | **-0.055** | -0.015 |
| $p$ value | <0.001* | <0.001* | 0.001* | 0.792 | 0.942 |

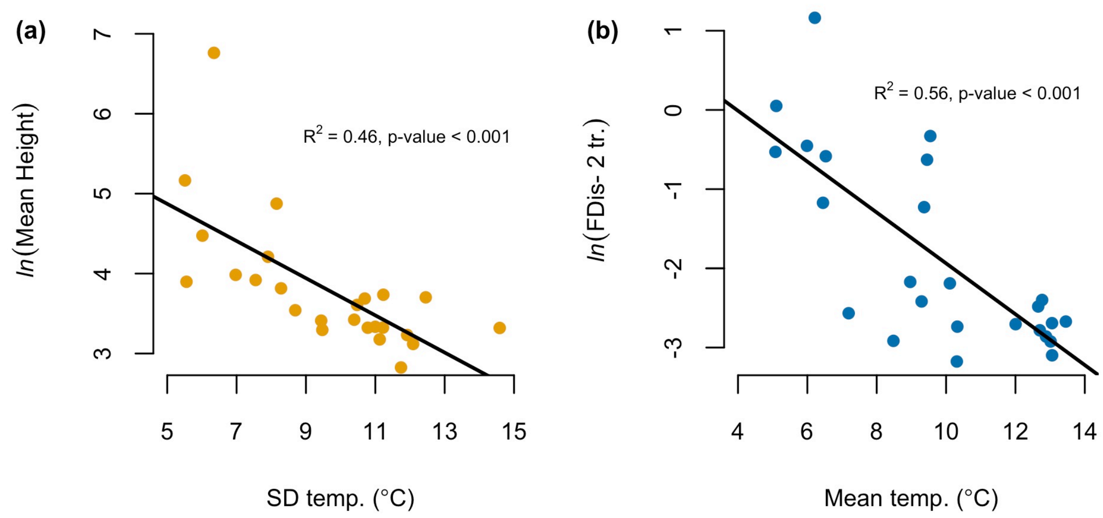


Figure 6: **Increases in soil temperature variation and mean soil temperature predict a decrease in the CWM of mean height and functional dispersion, calculated with two traits, respectively.** (a) Plots with less variation in soil temperatures support taller assemblages [higher logged CWM of mean height (cm), slope = -0.233, $F_{1,23}$ = 19.67]. (b) Even when FDis is calculated from only two traits—maximum height and SLA—plots with lower mean temperatures support assemblages with more functional variation [*i.e.* more biological insurance; higher FDis, slope = -0.321, $F_{1,23}$ = 29.19]. Color coding described in Figure 1 in the main text.

Table 4: **Functional dispersion and the CWM of traits vary for different life history strategies when aggregated across all twenty-five plots at Right Hand Fork.** Most species across the site were herbaceous perennials 54/84. Woody perennials had a functional dispersion over four times higher than herbaceous perennials or annuals/biennials. Herbaceous perennials had the most acquisitive leaves (highest SLA), followed by woody perennials and then annuals/biennials; however, they had much higher leaf area than either of the other groups. Woody perennials were much taller than other life history groups, whether measured as maximum or mean height.

| | Sp. richness | FDis | CWM SLA | CWM LA | CWM max. height | CWM mean height |
|---|---|---|---|---|---|---|
| Annual/biennials | 17 | 0.11 | 23.48 | 0.27 | 41.47 | 22.48 |
| Herbaceous perennials | 54 | 0.81 | 73.83 | 15.01 | 61.89 | 40.02 |
| Woody perennials | 13 | 4.20 | 34.96 | 3.33 | 1046.43 | 687.19 |

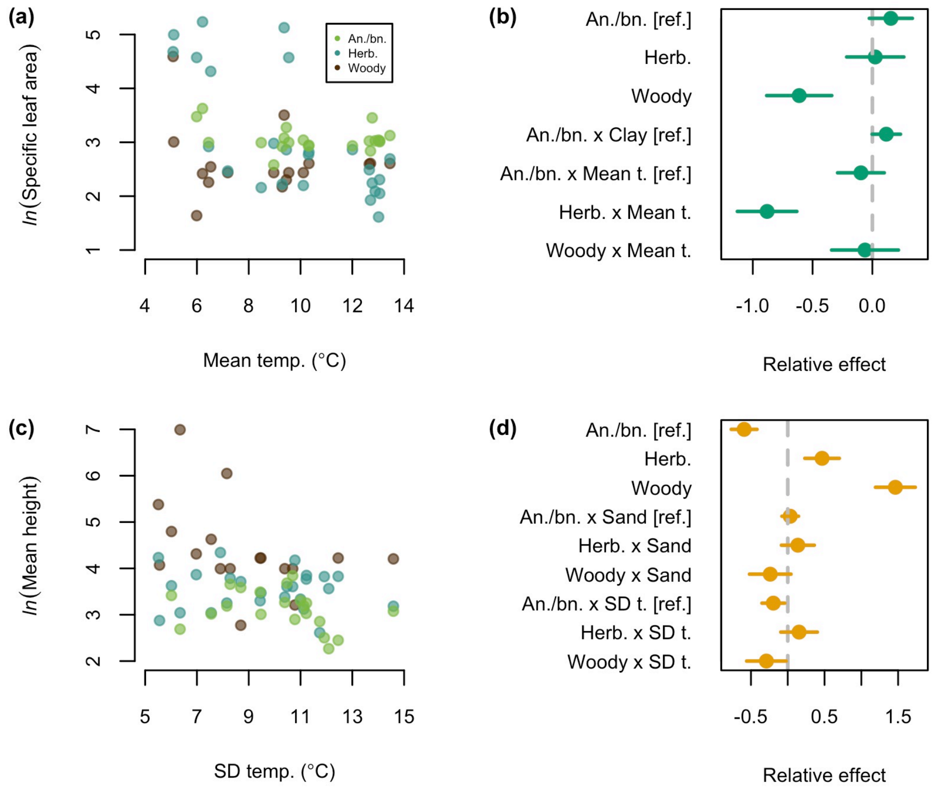


Figure 7: **Life history strategies affect the relationship between additional functional diversity metrics and microenvironment.** Plots in the left column show how the logged CWM of (a) specific leaf area and (c) and mean height, vary across the temperature variable they were most correlated with when subset by life history strategy — annuals/biennials (green), herbaceous perennials (blue), and woody perennials (brown). Plots in the right column show the relative effect of each explanatory variable in models that look at how these life history strategies affect the relationship between each functional metric (in the left column) and both the soil temperature and texture variable most correlated with that functional metric. Coefficient values are reference contrasts from those labeled as such. Values further from zero indicate that a variable or interaction between variables has a greater effect on a functional metric.
