## Supplement - data and analysis for "Variation in near-surface soil temperature drives plant assemblage insurance potential": env.R

```
# Microenvironment at core RHF plots: soil temp (2017-18 and 2018-19), texture, & terrain
# Elizabeth Simpson # 2019-07-29 | 2021-1-25
# Set working directory
# setwd("~/Documents/projects/functional_traits_rhf")

library(soiltexture)
library(lubridate)
library(readxl)
library(weathermetrics)
library(tidyverse)

#############
# FUNCTIONS #
#############

### To calculate soil texture ###

# Temperature correction function for hydrometer readings (+0.4 g/L for each degree above 20 C, - 0.4 g/L for each degree below 20 C)
temp.correct <- function(temp, hydro_rdg){
  return(hydro_rdg + ((temp-20)*0.4))
}

# Soil texture calculations
texture <- function(soil, plot_ID, soil_wt, temp_calib, hydro_calib, temp_40, hydro_40, temp_120, hydro_120){
  clean_soil <- matrix(nrow=nrow(soil), ncol=3)
  for(i in 1:nrow(soil)){
    B <- with(soil, temp.correct(temp_calib[i], hydro_calib[i]))
    A_40 <- with(soil, temp.correct(temp_40[i], hydro_40[i]))
    A_120 <- with(soil, temp.correct(temp_120[i], hydro_120[i]))
    silt_clay <- ((A_40-B)/soil_wt[i])*100
    clean_soil[i,1] <- 100-silt_clay #sand
    clean_soil[i,2] <- ((A_120-B)/soil_wt[i])*100 #clay
    clean_soil[i,3] <- 100 - (100-silt_clay) - (((A_120-B)/soil_wt[i])*100) #silt
  }
  colnames(clean_soil) <- c("SAND", "CLAY", "SILT")
  output <- as.data.frame(cbind(plot_ID, clean_soil))
  output$SAND <- as.numeric(output$SAND)
  output$CLAY <- as.numeric(output$CLAY)
  output$SILT <- as.numeric(output$SILT)
  return(output)
}

# To format times in the temperature dataset - based on code from Michael Stemkovski (MS)
format.times <- function(x){ 
  #x <- all_data[4,1]
  hour <- as.character(hour(x))
  minute <- as.character(minute(x))
  if(nchar(hour) == 1) hour <- paste(0,hour,sep="")
  if(nchar(minute) == 1) minute <- paste(0,minute,sep="")
  return(paste(hour,minute,sep=":"))
}

#############
# LOAD DATA #
#############

### Topography - clean data from fractal paper
topo <- read.csv("./clean_data/clean_rhf_terrain_17-18.csv", as.is=TRUE)

### Soil texture
s.text <- read.csv("./raw_data/rhf_2018_soil_texture.csv", as.is=TRUE)

### Temperature - modified from code from MS
# Note, converted from .hobo file to .csv in HoboWare
temp.csvs <- list.files("./raw_data/17-18_temp/")[which(grepl("csv",list.files("raw_data/17-18_temp")))]

##############
# CLEAN DATA #
##############

### Topography - remove extra first col and TPI from the dataframe
topo <- topo[,-c(1,10)]
topo$plot_id <- as.character(topo$plot_id)

### Soil texture
s.text <- na.omit(s.text)
text <- with(s.text, texture(s.text, plot_ID, soil_wt, temp_calib, hydro_calib, temp_40, hydro_40, temp_120, hydro_120))
text <- cbind(text, TT.points.in.classes(tri.data = text, class.sys = "USDA.TT", PiC.type="t"))
colnames(text) <- c("plot_id", "SAND", "CLAY", "SILT", "class")

### 2017-2018 temperature cleaning - based on code from MS
temp <- data.frame(datetime = rep(NA,10000*26),
                   temp = rep(NA,10000*26),
                   plot = rep(NA,10000*26))
top <- 1
for(i in temp.csvs){
  file <- read.csv(paste("raw_data/17-18_temp/",i,sep=""), header=F, as.is=TRUE)[,c(2,3)]
  colnames(file) <- c("datetime", "temp")  
  file <- file[-c(1,2),]
  file[,1] <- file[,1]
  file[,2] <- as.numeric(file[,2])
  file[,3] <- as.integer(substr(i,1,4)) 
  bottom <- top + nrow(file) - 1
  temp[top:bottom,] <- file
  top <- bottom + 1
}

temp <- temp[-which(is.na(temp$datetime)),]
temp[,1] <- mdy_hms(temp[,1], tz = "America/Denver") # if this throws an error, re install lubridate & restart R

# tnext lines takes a couple minutes to run
f_temp <- data.frame(plot = temp[,3],
                     datetime = temp[,1],
                     year = year(temp[,1]),
                     month = month(temp[,1]),
                     day = day(temp[,1]),
                     time = sapply(temp[,1], format.times),
                     temp = fahrenheit.to.celsius(temp[,2]))

f_temp <- na.omit(f_temp)

# Summarize date ranges to get time period completely covered by sensors - all ES code from hereon
date.range <- as.data.frame(f_temp %>%
                              group_by(plot) %>%
                              summarize(first.m=first(month),first.d=first(day), first.t=first(time), first.y=first(year),last.m=last(month),last.d=last(day), last.t=last(time), last.y=last(year)))

# Subset a full year of temperature data from 2017 -> 2018 matched to look at how microenvironment varies across topography
f_temp_year <- subset(f_temp,
                datetime >= as.POSIXct('2017-09-28 00:00:00',
                                        tz = "MST") &
                   datetime <= as.POSIXct('2018-09-28 00:00:00',
                                          tz = "MST"))

# Calculate statistics by plot, could also summarize at the monthly level first and then at plot
year_17.18 <- as.data.frame(group_by(f_temp_year, plot) %>%
                           summarize(mean(temp), sd(temp), max(temp), min(temp)))
colnames(year_17.18) <- c("plot_id", "mean", "sd", "max", "min")

#########################
# COMBINE & EXPORT DATA #
#########################

# topography + soil texture - 76 plots
env <- merge(topo, text, by.x="plot_id", by.y="plot_id")

# Terrain, texture, and temperature (2017-18) - 25 plots 
env.t <- merge(env, year_17.18, by.x="plot_id", by.y="plot_id")
write.csv(env.t, "./clean_data/temp-year17-18_text_terr.csv")
```
