## Supplement - data and analysis for "Variation in near-surface soil temperature drives plant assemblage insurance potential": fdiv-env-phy_core18.R

```
# Assessing variance in functional dispersion and CWM of height and leaf traits across microenvironments
# Elizabeth Simpson # 2021-03-25
# Uses code from Will Pearse to get comparative.comm set up

setwd("~/Documents/projects/functional_traits_rhf") # uncomment to set working directory

###############
### HEADERS ###
library(pez)
library(FD)
library(lme4)
library(soiltexture)
library(xtable)
library(MuMIn)
library(RColorBrewer)
library(magick)

#################
### LOAD DATA ###
env.micro <- read.csv("./clean_data/temp-year17-18_text_terr.csv", as.is=TRUE)
d.eight <- read.csv("./raw_data/rhf_2018_cover.csv", as.is = TRUE)
tree <- read.tree("./clean_data/Vascular_Plants_rooted.dated.tre")
env.cvr18 <- read.csv("./clean_data/temp-year17-18_text_terr.csv", as.is=TRUE)
traits <- read.csv("./clean_data/clean_traits_rhf.csv", as.is=TRUE)

#################
### FUNCTIONS ###

# Add in overhanging cover
calc.over <- function(cover){
  cover$Cover.over <- NA
  for (i in seq_len(nrow(cover))){
    if (with(cover, grepl("overhang-", Notes[i]))){
      cover$Cover.over[i] <- with(cover, Cover[i] + as.numeric(unlist(regmatches(Notes[i], gregexpr('\\(?[0-9,.]+', Notes[i])))))
    } else {
      cover$Cover.over[i] <- cover$Cover[i]
    }
  }
  return(cover)
}

# Clean the community data
calc.comm <- function(cover){
  comm <- with(cover, tapply(Cover.over, list(Plot_id,Species), function(x) mean(x,na.rm=TRUE)))
  comm[is.na(comm)] <- 0
  comm <- comm[, colSums(comm != 0) > 0]
  comm <- comm[,!grepl("^[a-z]+", colnames(comm))]
  comm <- comm[,!grepl("\\(|/", colnames(comm))]
  comm <- comm[,!grepl("sp\\.", colnames(comm))]
  colnames(comm) <- tolower(colnames(comm))
  return(comm)
} 

# Calculate diversity metrics and add environmental data all to the same dataframe
# Focus on functional dispersion and community weighted means but the 3 Villeger et al. 2008 indicies are also in here
# Keep default arguments that weight each trait by a species' abundance in the community and standardize each trait value
div.calc <- function(c.data){
  # Calculate functional, phylogenetic, and taxonomic diversity indices
  # Focus on functional dispersion and community weighted means but the 3 Villeger et al. 2008 indicies are also in here
  f.div <- with(c.data, dbFD(data[,c(1:2,4:5)], comm, w.abun=TRUE, stand.x=TRUE, print.pco=TRUE))
  f.distwo <- with(c.data, dbFD(data[,c(1,4)], comm, w.abun=TRUE, stand.x=TRUE, print.pco=TRUE))
  # Put together all of the pieces needed
  div <- with(f.div, cbind(FDis, CWM))
  div <- with(f.distwo, cbind(div, FDis))
  colnames(div) <- c("fdis","CWM.SLA", "CWM.LA", "CWM.maxht","CWM.mn.ht", "fdis.two")
  return(cbind(div, c.data$env))
}

# Calculate Pearson's r and p-values between microenvironment and topography
# calculated the corrleation of the first input to all the rest
all.corr <- function(...){
  data <- list(...)
  n <- length(data)
  output <- matrix(nrow=2, ncol=n-1)
  for (i in 1:(n-1)){
    output[,i] <- c(cor.test(data[[1]], data[[i+1]])$estimate, cor.test(data[[1]], data[[i+1]])$p.value)
  }
  output <- as.data.frame(output)
  rownames(output) <- c("pearsons.r", "p-value")
  colnames
  return(output)
}

#################################################################################################################
### Q1: How does microenvironment (soil temp. and texture) vary across topography (aspect, elevation, slope)? ###
# Temperature from 2017-09-28 at 00:00 to 2018-09-29 at 00:00 at 25 plots.
# Note: Temp. sensors serviced mid-Sept. 2018. A few time points are missing on the days that that happened.

env18.core <- env.micro[,c(7:12,14:17)]

### Does temperature correlate with soil texture at all?
mn.text <- with(env18.core, all.corr(mean, SAND, SILT, CLAY))
sd.text <- with(env18.core, all.corr(sd, SAND, SILT, CLAY))
max.text <- with(env18.core, all.corr(max, SAND, SILT, CLAY))
min.text <- with(env18.core, all.corr(min, SAND, SILT, CLAY))

temp.text <- rbind(mn.text, sd.text, max.text, min.text)
colnames(temp.text) <- c("Sand", "Silt", "Clay")
xtable(temp.text, digits=3)

# Because of low replication (25 plots), only have statistical power to do a univariate analysis.
# So, look at which topo. var. is most correlated with each microenv. var. using Pearson's r
mn.topo <- with(env18.core, all.corr(mean, aspect, elev, slope))
sd.topo <- with(env18.core, all.corr(sd, aspect, elev, slope))
max.topo <- with(env18.core, all.corr(max, aspect, elev, slope))
min.topo <- with(env18.core, all.corr(min, aspect, elev, slope))
sand.topo <- with(env18.core, all.corr(SAND, aspect, elev, slope))
silt.topo <- with(env18.core, all.corr(SILT, aspect, elev, slope))
clay.topo <- with(env18.core, all.corr(CLAY, aspect, elev, slope))

env.topo <- rbind(mn.topo, sd.topo, max.topo, min.topo, sand.topo, silt.topo, clay.topo)
colnames(env.topo) <- c("aspect", "elevation", "slope")
xtable(env.topo, digits=3)

# Univariate models of how temperature and texture vary across microenvironment
mn.a <- with(env18.core, lm(mean~aspect))
summary(mn.a)
sd.a <- with(env18.core, lm(sd~aspect))
summary(sd.a)
max.a <- with(env18.core, lm(max~aspect))
summary(max.a)
min.1 <- with(env18.core, lm(min~1))
summary(min.1)

sand.e <- with(env18.core, lm(SAND~elev))
summary(sand.e)
silt.1 <- with(env18.core, lm(SILT~1))
summary(silt.1)
clay.e <- with(env18.core, lm(CLAY~elev))
summary(clay.e)

# FIG 2 - PLOTTING changes in temp. and texture across microenvironment 
# plus the USDA soil texture triangle for all of the plots
jpeg("./analysis/figures/env-core18.jpeg", width=7, height=7, unit="in",res=300)
par(mfrow=c(2,2))
par(mar=c(5,5,2,0.5))
par(oma=c(1,1,1,1))

with(env18.core, plot(mean~aspect, pch=16, xlab="Aspect (S = -1, N = 1)", ylab=expression(paste('Temperature (',degree~'C)')), cex=1.2, axes=FALSE, ylim=c(-10,70), col="#E69F00"))
abline(mn.a, lwd=3, col="#E69F00")
with(env18.core, points(max~aspect, pch=16, cex=1.2, col="#D55E00"))
abline(max.a, lwd=3, col="#D55E00")
with(env18.core, points(min~aspect, pch=16, cex=1.2, col="#56B4E9"))
axis(1)
axis(2, at=c(-10,10,30,50,70))
mtext(expression(bold("(a)")),side=3,line=0.2, adj=-0.32)
legend(-0.5,62, expression(~ R^2 ~ "= 0.35, p-value = 0.002"), text.col="#D55E00", box.lty=0, bg="transparent", cex=0.8)
legend(-1.2,25, expression(~ R^2 ~ "= 0.68, p-value < 0.001"), text.col= "#E69F00", box.lty=0, bg="transparent", cex=0.8)

with(env18.core, plot(sd~aspect, pch=16, xlab="Aspect (S = -1, N = 1)", ylab=expression(paste('SD - Temperature (',degree~'C)')), cex=1.2, axes=FALSE, col="#E69F00", ylim=c(5,15)))
abline(sd.a, lwd=3, col="#E69F00")
axis(1)
axis(2, at=c(5,10,15))
mtext(expression(bold("(b)")),side=3,line=0.2, adj=-0.32)
legend(0.5,14.5, legend=c('Min', 'Mean', 'Max'), pch=16, pt.cex=1.2, cex=0.6, col = c('#56B4E9', '#E69F00', '#D55E00'))
legend(-1.2,6.5, expression(~ R^2 ~ "= 0.46, p-value < 0.001"), text.col="#E69F00", box.lty=0, bg="transparent", cex=0.8)

with(env18.core, plot(SILT~elev, pch=16, xlab="Elevation (m.s.l.)", ylab= "Soil component (%)", cex=1.2, axes=FALSE, col="gray", xlim=c(1700,2100), ylim=c(0,80)))
with(env18.core, points(SAND~elev, pch=16, cex=1.2, col="goldenrod3"))
abline(sand.e, lwd=3, col="goldenrod3")
with(env18.core, points(CLAY~elev, pch=16, cex=1.2, col="sienna"))
abline(clay.e, lwd=3, col="sienna")
axis(1)
axis(2)
legend(2010,80, legend=c("Sand", "Silt","Clay"), pch=16, pt.cex=1.2, cex=0.6, col = c("goldenrod3","gray", "sienna"))
legend(1670,70, expression(~ R^2 ~ "= 0.20, p-value = 0.024"), text.col="goldenrod3", box.lty=0, bg="transparent", cex=0.8)
legend(1835,29, expression(~ R^2 ~ "= 0.23, p-value = 0.016"), text.col="sienna", box.lty=0, bg="transparent", cex=0.8)
mtext(expression(bold("(c)")),side=3,line=0.2, adj=-0.32)

TT.plot(class.sys = "USDA.TT", tri.data = env18.core, pch=19, cex.axis=0.8, cex.lab=0.8, lwd=0.8, lwd.axis=0.8, lwd.lab=0.8, main="", cex=0.95, new.mar=c(3.7,3.7,0,0))
mtext(expression(bold("(d)")),side=3,line=-2, adj=-0.19)

dev.off()

####################################################################################################################################################################
### Q2: Do spatial differences in microenvironment predict variation in the mean and/or variance of functional diversity? ##########################################
### Sub-Q2: Do species that use a particular life history strategy --- woody perennial, herbaceous perennial, or annual/biennial --- drive these relationships ? ###

# VEG. COVER - overhang included - int.18.over includes all plots surveyed in 2018 (78)
# Subset to the core 25 plots where both temp. and texture were collected and make community matrix
int.18.over <- calc.over(d.eight)
core.18.over <- int.18.over[int.18.over$Plot_id %in% env.cvr18$plot_id,]
comm18.core <- calc.comm(core.18.over)

### PHYLOGENY (Zanne et. al. 2014) - reformat sp. names, subset to just the species in the core plots
tree$tip.label <- tolower(gsub("_", " ", tree$tip.label))
tree18.core <- congeneric.merge(tree, colnames(comm18.core), split = " ")

### TRAITS - leaves and plant height
rownames(traits) <- tolower(traits$species)
traits <- traits[,c(3:5,9:10)]

### ENVIRONMENT - Uses temperature data collected from 2017-09-28 at 00:00 to 2018-09-28 at 00:00 at 25 plots.
rownames(env18.core) <- as.character(env.cvr18$plot_id)

### Make COMPARATIVE COMMUNITY OBJECT & calculate diversity metrics
core18cdata <- comparative.comm(tree18.core, comm18.core, traits, env18.core)
core18div <- div.calc(core18cdata)

### MODELS
# Because of low replication (25 plots), only have statistical power to look at one temp. and texture variable, additively
# So, look at which temp. var. and which text. var is most correlated with each diversity metric using Pearson's r
# Addtl. note: There is also covariation within temp. and within texture variables
# --> another good reason to look at one microenvironment variable within each of these categories

## Correlations btn. fdiv and microenv (but just mean, sd, max, sand, and clay, because these were the ones that correlated with topography)
# (now the analysis builds on itself!)
la.env <- with(core18div, all.corr(log(CWM.LA), mean, sd, max, SAND, CLAY))
sla.env <- with(core18div, all.corr(log(CWM.SLA), mean, sd, max, SAND, CLAY))
mxH.env <- with(core18div, all.corr(log(CWM.maxht), mean, sd, max, SAND, CLAY))
mnH.env <- with(core18div, all.corr(log(CWM.mn.ht), mean, sd, max, SAND, CLAY))
fdis.env <- with(core18div, all.corr(log(fdis), mean, sd, max, SAND, CLAY))
fdis.two.env <- with(core18div, all.corr(log(fdis.two), mean, sd, max, SAND, CLAY))

fdiv.env <- rbind(la.env, sla.env, mxH.env, mnH.env, fdis.env, fdis.two.env)
colnames(fdiv.env) <- c("mean", "sd", "max", "sand", "clay")
xtable(fdiv.env, digits=3)


### CWM of LA
la <- with(core18div, lm(log(CWM.LA)~mean+CLAY))
summary(la)
la.mn <- with(core18div, lm(log(CWM.LA)~mean))
summary(la.mn)
anova(la, la.mn) # not significant, plot la.mn

### CWM of SLA
sla <- with(core18div, lm(log(CWM.SLA)~mean+CLAY))
summary(sla)
sla.mn <- with(core18div, lm(log(CWM.SLA)~mean))
summary(sla.mn)
anova(sla, sla.mn) # not significant, plot sla.mn

### CWM of max HT
mxH <- with(core18div, lm(log(CWM.maxht)~sd+SAND)) 
summary(mxH)
mxH.sd <- with(core18div, lm(log(CWM.maxht)~sd))
summary(mxH.sd)
anova(mxH, mxH.sd) # not significant, plot mxH.sd

### CWM of mean HT
mnH <- with(core18div, lm(log(CWM.mn.ht)~sd+SAND)) 
summary(mnH)
mnH.sd <- with(core18div, lm(log(CWM.mn.ht)~sd))
summary(mnH.sd)
anova(mnH, mnH.sd) # not significant, plot mnH.mx

### FDis - with four traits
fdis.4 <- with(core18div, lm(log(fdis)~mean+CLAY)) 
summary(fdis.4)
fdis.4.mn <- with(core18div, lm(log(fdis)~mean))
summary(fdis.4.mn)
anova(fdis.4, fdis.4.mn) # not significant, plot fdis.4.mx

### FDIS - with just two traits, max.height and LA
fdis.2 <- with(core18div, lm(log(fdis.two)~mean+SAND))
summary(fdis.2)
fdis.2.mn <- with(core18div, lm(log(fdis.two)~mean))
summary(fdis.2.mn)
anova(fdis.2, fdis.2.mn) # not significant, plot fdis.2.mx

### FIG. 3 - PLOTTING: how logged mean and variance in function shifts across microenvironment
jpeg("./analysis/figures/fdiv-env-core18.jpeg", width=7, height=7, unit="in",res=300)
par(mfrow=c(2,2))
par(mar=c(4,5,0.1,0.5))
par(oma=c(1.5,1.5,1.5,1.5))

# LA ~ mean
with(core18div, plot(log(CWM.LA)~mean, pch=19, xlab=expression(paste('Mean temp. (',degree,'C)')), ylab=expression(italic("ln")("Leaf area")), cex=0.9, axes=FALSE, xlim=c(4,14),ylim=c(-3,3.4), col="#009E73" )) # col was
abline(la.mn, lwd=2)
axis(1, cex.lab=1.5)
axis(2, at=c(-3,-1,1,3), cex.lab=1.5)
legend(4,-2, expression(R^2 ~ "= 0.68, p-value < 0.001"), box.lty=0, bg="transparent", cex = 0.8)
mtext(expression(bold("(a)")),side=3, line=-2, adj=-0.35, cex = 0.85)

# mxH ~ sd
with(core18div, plot(log(CWM.maxht)~sd, pch=19, xlab=expression(paste('SD temp. (',degree,'C)')), ylab=expression(italic("ln")("Max. Height")), cex=0.9, axes=FALSE, xlim=c(5,15),ylim=c(3,7.3),col="#E69F00")) 
abline(mxH.sd, lwd=2)
axis(1, at=c(5,7,9,11,13,15),cex.lab=1.5)
axis(2, at=c(3,4,5,6,7), cex.lab=1.5)
legend(8,6.5, expression(R^2 ~ "= 0.49, p-value < 0.001"), box.lty=0, bg="transparent", cex = 0.8)
mtext(expression(bold("(b)")),side=3, line=-2, adj=-0.35, cex = 0.85)

# SLA ~ mean
with(core18div, plot(log(CWM.SLA)~mean, pch=19, xlab=expression(paste('Mean temp. (',degree,'C)')), ylab=expression(italic("ln")("Specific leaf area")), cex=0.9, axes=FALSE, xlim=c(4,14), ylim=c(2,5.2), col="#009E73" ))
abline(sla.mn, lwd=2)
axis(1, cex.lab=1.5)
axis(2, at=c(2,3,4,5), cex.lab=1.5)
legend(6.7,4.2, expression(R^2 ~ "= 0.29, p-value = 0.005"), box.lty=0, bg="transparent", cex = 0.8)
mtext(expression(bold("(c)")),side=3, line=-2, adj=-0.35, cex = 0.85)

# fdis.4 ~ mean
with(core18div, plot(log(fdis)~mean, pch=19, xlab=expression(paste('Mean temp. (',degree,'C)')), ylab=expression(italic("ln")("FDis")), cex=0.9, axes=FALSE, xlim=c(4,14), ylim=c(-3,2.3), col="#0072B2")) 
abline(fdis.4.mn, lwd=2)
axis(1, cex.lab=1.5)
axis(2, at=c(-3, -2, -2, -1, 0, 1, 2), cex.lab=1.5)
legend(7,1, expression(R^2 ~ "= 0.56, p-value < 0.001"), box.lty=0, bg="transparent", cex = 0.8)
mtext(expression(bold("(d)")),side=3, line=-2, adj=-0.35, cex = 0.85)

dev.off()

# PLOT the similar ones in  the supplement
jpeg("./analysis/figures/SUPP-fdiv-env-core18.jpeg", width=7, height=3.5, unit="in",res=300)
par(mfrow=c(1,2))
par(mar=c(4,5,0.1,0.5))
par(oma=c(1.5,1.5,1.5,1.5))

# mnH ~ mean --> bump to the supplement
with(core18div, plot(log(CWM.mn.ht)~sd, pch=19, xlab=expression(paste('SD temp. (',degree,'C)')), ylab=expression(italic("ln")("Mean Height")), cex=0.8, cex.lab=0.8,axes=FALSE, xlim=c(5,15), ylim=c(2.9,7.2), col="#E69F00")) 
abline(mnH.sd, lwd=2)
axis(1, at=c(5,7,9,11,13,15), cex.axis=0.8)
axis(2, at=c(3,4,5,6,7), cex.axis=0.8)
legend(8,6, expression(R^2 ~ "= 0.46, p-value < 0.001"), box.lty=0, bg="transparent", cex = 0.6)
mtext(expression(bold("(a)")),side=3, line=-1.3, adj=-0.4, cex = 0.85)

with(core18div, plot(log(fdis.two)~mean, pch=19, xlab=expression(paste('Mean temp. (',degree,'C)')), ylab=expression(italic("ln")("FDis- 2 tr.")), cex=0.8,cex.lab=0.8, axes=FALSE, xlim=c(4,14), col="#0072B2")) 
abline(fdis.2.mn, lwd=2)
axis(1, cex.axis=0.8)
axis(2, at=c(-3, -2, -2, -1, 0, 1), cex.axis=0.8)
legend(7,0.5, expression(R^2 ~ "= 0.56, p-value < 0.001"), box.lty=0, bg="transparent", cex = 0.6)
mtext(expression(bold("(b)")),side=3, line=-1.3, adj=-0.4, cex = 0.85)

dev.off()

#######################################################################################################################################################################
### Q3: How does including information about phylogenetic differences (in addition to functional differences) between species #########################################
### change our understanding of the ecological differences between these assemblages and how those assemblages change across spatially different microenvironments? ###

# Calculate SESmntd for different values of the phylogenetic weighting parameter (a)
# where a=0 means all difference is functionally based and a=1 means all differences is phylogenetically based 
# (this takes a few minutes)
# subset to species richness, SESmntd and the value of a
fun.phy.calc <- pez.dispersion(core18cdata, null.model="taxa.labels", abundance=TRUE, traitgram=seq(0,1,0.1))
fun.phy.s <- with(fun.phy.calc, cbind(ses.mntd.mntd.obs.z, traitgram))
fun.phy.s <- as.data.frame(fun.phy.s)
colnames(fun.phy.s) <- c("mntd", "a.f.p")

# replicate environmental variables for add-in
env18.core.rep <- sapply(env18.core, rep.int, times=11)
# combine the diversity and env info together
fun.phy <- cbind(fun.phy.s, env18.core.rep)

### Q3a: How does mntd vary overall, depending on how much functional (a=0) vs. phylogenetic (a=1) influence on difference?
mean.fp <- matrix(NA,nrow=11,ncol=3)
mean.fp[,1] <- seq(0,1,0.1)
mean.fp[,2] <- with(fun.phy, tapply(mntd, a.f.p, na.rm=TRUE,FUN=mean))
mean.fp[,3] <- with(fun.phy, tapply(mntd, a.f.p, na.rm=TRUE, FUN=sd)/sqrt(25))
mean.fp <- as.data.frame(mean.fp)
colnames(mean.fp) <- c("a", "mean.mntd", "se.mntd")

fp.mntd.anova <- with(fun.phy, aov(mntd ~ a.f.p))
summary(fp.mntd.anova) # no

### Q3b: Is there a difference in these relationships depending on the degree to which phylogeny is included?

# How the relationship between MNTD and mean temperature changes as the value of a changes
fp.mn.slope <- matrix(nrow=11, ncol=4)
for(i in seq_along(unique(fun.phy$a.f.p))){
  model.mn <- with(fun.phy[fun.phy$a.f.p == unique(fun.phy$a.f.p)[i],], lm(mntd~mean))
  fp.mn.slope[i,1] <- summary(model.mn)$coefficients[2,1]
  fp.mn.slope[i,2] <- summary(model.mn)$coefficients[2,4]
  fp.mn.slope[i,3] <- summary(model.mn)$r.squared
  fp.mn.slope[i,4] <- summary(model.mn)$fstatistic[1]
}

fp.mn.slope <- as.data.frame(fp.mn.slope)
colnames(fp.mn.slope) <- c("slope", "p-value", "r2", "f-statistic_{1,23}" )
fp.mn.slope$a.f.p <- seq(0,1,0.1)
xtable(fp.mn.slope, digits=3)

# How the relationship between MNTD and clay (%) changes as the value of a changes
fp.c.slope <- matrix(nrow=11, ncol=4)
for(i in seq_along(unique(fun.phy$a.f.p))){
  model.c <- with(fun.phy[fun.phy$a.f.p == unique(fun.phy$a.f.p)[i],], lm(mntd~CLAY))
  fp.c.slope[i,1] <- summary(model.c)$coefficients[2,1]
  fp.c.slope[i,2] <- summary(model.c)$coefficients[2,4]
  fp.c.slope[i,3] <- summary(model.c)$r.squared
  fp.c.slope[i,4] <- summary(model.c)$fstatistic[1]
}
fp.c.slope <- as.data.frame(fp.c.slope)
colnames(fp.c.slope) <- c("slope", "p-value", "r2", "f-statistic_{1,23}" )
fp.c.slope$a.f.p <- seq(0,1,0.1)
xtable(fp.c.slope, digits=3)

# FIG 4 - PLOTTING
ub <- with(mean.fp, as.vector(mean.mntd+se.mntd))
lb <- with(mean.fp, as.vector(mean.mntd-se.mntd))

svg("./analysis/figures/funct-phy-core18-raw.svg", width=10, height=2.7)
par(mfrow=c(1,4))
par(mar=c(4,5,0.1,0.5))
par(oma=c(1.5,1.5,1.5,1.5))

# how the value of mntd changes across a as different amounts of function and phylogeny are included
with(mean.fp, plot(mean.mntd~a, ylim=c(-1.3, -0.5), pch=19, col=brewer.pal(11,"BrBG")[11:1], axes=FALSE, 
                   ylab=expression("Mean SES"[MNTD]), xlab=expression(italic("a"))))
with(mean.fp, points(mean.mntd~a, cex=1.2))
axis(1)
axis(2, at=c(-1.3, -1.1, -0.9, -0.7, -0.5))
segments(mean.fp$a, ub, mean.fp$a, lb)
mtext(expression(bold("(a)")),side=3, line=-0.5, adj=-0.42, cex = 0.8)

# across mean
with(fp.mn.slope, plot(slope~a.f.p, pch=19, col=brewer.pal(11,"BrBG")[11:1], axes=FALSE, ylim=c(-0.40, -0.10),
                         ylab=expression(paste('Slope('~SES[MNTD]~'~ mean temp.)')), xlab=expression(italic(a))))
with(fp.mn.slope, points(slope~a.f.p, cex=1.2))
axis(1)
axis(2, at=c(-0.4, -0.3, -0.2, -0.1))
mtext(expression(bold("(b)")),side=3, line=-0.5, adj=-0.42, cex = 0.8)

# across clay
with(fp.c.slope, plot(slope~a.f.p, pch=19, col=brewer.pal(11,"BrBG")[11:1], axes=FALSE, ylim=c(-0.0005, 0.0205),
                       ylab=expression(paste('Slope('~SES[MNTD]~'~ % clay)')), xlab=expression(italic(a))))
with(fp.c.slope, points(slope~a.f.p, cex=1.2))
axis(1)
axis(2, at=c(0, 0.01, 0.02))
mtext(expression(bold("(c)")),side=3, line=-0.5, adj=-0.42, cex = 0.8)

par(pin=c(0.07,0.7))
image(1, unique(fun.phy$a.f.p), t(seq_along(unique(fun.phy$a.f.p))), col=brewer.pal(11, "BrBG")[11:1], axes=FALSE, xlab="", ylab="")
axis(4, las=1, cex=0.3)
title(expression(italic(a)), line=0.7, cex=0.8)

dev.off()
```
