## Supplement - data and analysis for "Variation in near-surface soil temperature drives plant assemblage insurance potential": fdiv-env_life-hist_core18.R

```
# Assessing how categorization of species by life histories affects FT~env analysis
# Elizabeth Simpson
# 2022-08-08
source("~/Documents/projects/functional_traits_rhf/analysis/fdiv-env-phy_core18.R")

########################################################################################################################################
### Sub-Q2: Do species that use a particular life history strategy --- woody perennial, herbaceous perennial, or annual/biennial --- ###
### drive the relationships betwee the mean and variance of functional diversity and spatial variable microenvironment? ################

#######################################
### Load & format life history data ###
cat.tr <- read.csv("./raw_data/cat_traits_rhf.csv", as.is=TRUE)
lh.tr <- as.data.frame(cat.tr[,2])
rownames(lh.tr) <- cat.tr$species
colnames(lh.tr) <- "life_hist"

### Make a species lists for each life history strategy
# Woody perennials
p.wood <- cat.tr$species[cat.tr$life_history=="perennial-woody"]
# Herbaceous perennials
p.herb <- cat.tr$species[cat.tr$life_history=="perennial-herb"]
# Annual/Biennials
anubi <- cat.tr$species[cat.tr$life_history=="annual"]

# Add life history type to cover data
cover.lh <- core.18.over
for(i in seq_len(nrow(cover.lh))){
  if(cover.lh$Species[i] %in% p.wood){
    cover.lh$lh[i] <- "P_wood"
  } else {
    if(cover.lh$Species[i] %in% p.herb){
      cover.lh$lh[i] <- "P_herb"
    } else{
      if(cover.lh$Species[i] %in% anubi){
        cover.lh$lh[i] <- "Anubi"
      } else {
        cover.lh$lh[i] <- "na"
      }
    }
  }
}

##############################################################################################################################################
### Sub-Q2a: What is the overall difference in the meand and variance of functional traits in each of these three life history categories? ###

# Clean and re-sort all cover data by hand (w/o function, because it doesn't match) to rename as the three different types of things
comm.lh <- with(cover.lh, tapply(Cover.over, list(lh, Species), function(x) mean(x,na.rm=TRUE)))
comm.lh[is.na(comm.lh)] <- 0
comm.lh <- comm.lh[, colSums(comm.lh != 0) > 0]
comm.lh <- comm.lh[,!grepl("^[a-z]+", colnames(comm.lh))]
comm.lh <- comm.lh[!grepl("^[a-z]+", rownames(comm.lh)),]
comm.lh <- comm.lh[,!grepl("\\(|/", colnames(comm.lh))]
comm.lh <- comm.lh[,!grepl("sp\\.", colnames(comm.lh))]
colnames(comm.lh) <- tolower(colnames(comm.lh))

# Just look at functional dispersion characterisitics
cdata.lh <- comparative.comm(phy=tree18.core, comm=comm.lh, traits=traits)

# Calculate functional, phylogenetic, and taxonomic diversity indices
# Focus on functional dispersion and community weighted means but the 3 Villeger et al. 2008 indicies are also in here
# Weighted by abundance, traits standardized
fdiv.lh <- with(cdata.lh, dbFD(data[,c(1:2,4:5)], comm))

# Put together all of the pieces needed
div.lh <- with(fdiv.lh, cbind(nbsp, FDis, CWM))
colnames(div.lh) <- c("nsp","fdis","CWM.SLA", "CWM.LA", "CWM.maxht","CWM.mn.ht")
xtable(div.lh, digits=2) # SUPPLEMENT

######################################################################
### Sub-Q2b: Actually answering Sub-Q2 from the top of this script ###

# Sort out the 2018 cover data from the 25 core plots with temperature data into these life hist. cats.
wood <- core.18.over[core.18.over$Species %in% p.wood,]
herb <- core.18.over[core.18.over$Species %in% p.herb,]
anbi <- core.18.over[core.18.over$Species %in% anubi,]

# Make community matrices for each of these and remove communities that have zero abundance across all species types
# Remove communities that have zero abundance across all species
wood.comm <- calc.comm(wood)
wood.comm <- wood.comm[,colnames(wood.comm) %in% rownames(traits)]
wood.comm <- wood.comm[rowSums(wood.comm)>0,]

herb.comm <- calc.comm(herb)
herb.comm <- herb.comm[,colnames(herb.comm) %in% rownames(traits)]
herb.comm <- herb.comm[rowSums(herb.comm)>0,]

anbi.comm <- calc.comm(anbi)
anbi.comm <- anbi.comm[,colnames(anbi.comm) %in% rownames(traits)]
anbi.comm <- anbi.comm[rowSums(anbi.comm)>0,]

###################################
# Make comparative community object & calculate diversity metrics from that
wood.cdata <- comparative.comm(tree18.core, wood.comm, traits, env18.core) # 17 plots
wood.div <- div.calc(wood.cdata)
wood.div$lh <- "wood"
herb.cdata <- comparative.comm(tree18.core, herb.comm, traits, env18.core) # 25 plots
herb.div <- div.calc(herb.cdata)
herb.div$lh <- "herb"
anbi.cdata <- comparative.comm(tree18.core, anbi.comm, traits, env18.core) # 21 plots
anbi.div <- div.calc(anbi.cdata)
anbi.div$lh <- "anbi"

 # COMBINED them into one data frame
unscaled.lh.div <- rbind(wood.div, herb.div, anbi.div) #unscaled
log.lh.div <- unscaled.lh.div
log.lh.div[,2:5] <- log(log.lh.div[,2:5]) # log all leaf and height trait values, but NOT fdis, because some values are zero
lh.div <- log.lh.div
# scale them so the coefficients are comparable
lh.div[,1:16] <- as.numeric(scale(lh.div[,1:16]))

### CWM of LA
la.lh <- with(lh.div, lm(CWM.LA~(mean*lh)+(CLAY*lh), na.action=na.pass))
la.dredge <- dredge(la.lh, beta="none", rank="AIC")
summary(model.avg(la.dredge, subset=delta<4, fit=TRUE))
#get.models(la.dredge, subset=TRUE) # uncomment to look at the models - models are all variations on a theme - it doesn't look like anything conflicting is happening to me

# Plotting setup for LA 
la.coef <- coefTable(model.avg(la.dredge, subset=delta<4, fit=TRUE))
colnames(la.coef)[colnames(la.coef)=="Std. Error"] <- "se"
la.coef <- as.data.frame(la.coef[,1:2])
la.coef <- la.coef[c(1,3,4,2,6,7,5,8,9),]

la.coef$var <- c("An./bn. [ref.]", "Herb.", "Woody", "An./bn. x Clay [ref.]", "Herb. x Clay", "Woody x Clay", "An./bn. x Mean t. [ref.]", "Herb. x Mean t.", "Woody x Mean t.")
la.coef$num <- seq(9,1,-1)
la.coef$ub <- with(la.coef, as.vector(Estimate+se))
la.coef$lb <- with(la.coef, as.vector(Estimate-se))

### CWM of SLA
sla.lh <- with(lh.div, lm(CWM.SLA~(mean*lh)+(CLAY*lh), na.action=na.pass))
sla.dredge <- dredge(sla.lh, beta="none", rank="AIC")
summary(model.avg(sla.dredge, subset=delta<4, fit=TRUE))
#get.models(la.dredge, subset=TRUE) # uncomment to look at the models - models are all variations on a theme - it doesn't look like anything conflicting is happening to me

# Plotting setup for SLA 
sla.coef <- coefTable(model.avg(sla.dredge, subset=delta<4, fit=TRUE))
colnames(sla.coef)[colnames(sla.coef)=="Std. Error"] <- "se"
sla.coef <- as.data.frame(sla.coef[,1:2])
sla.coef <- sla.coef[c(1,2,3,7,4,5,6),]

sla.coef$var <- c("An./bn. [ref.]", "Herb.", "Woody", "An./bn. x Clay [ref.]", "An./bn. x Mean t. [ref.]", "Herb. x Mean t.", "Woody x Mean t.")
sla.coef$num <- seq(7,1,-1)
sla.coef$ub <- with(sla.coef, as.vector(Estimate+se))
sla.coef$lb <- with(sla.coef, as.vector(Estimate-se))

### max height
mxH.lh <- with(lh.div, lm(CWM.maxht~(sd*lh)+(SAND*lh), na.action=na.pass))
mxH.dredge <- dredge(mxH.lh, beta="none", rank="AIC")
summary(model.avg(mxH.dredge, subset=delta<4, fit=TRUE))
#get.models(mxH.dredge, subset=TRUE)

# Plotting setup for mxH
mxH.coef <- coefTable(model.avg(mxH.dredge, subset=delta<4, fit=TRUE))
colnames(mxH.coef)[colnames(mxH.coef)=="Std. Error"] <- "se"
mxH.coef <- as.data.frame(mxH.coef[,1:2])
mxH.coef <- mxH.coef[c(1,2,3,7,8,9,4,5,6),]

mxH.coef$var <- c("An./bn. [ref.]", "Herb.", "Woody", "An./bn. x Sand [ref.]", "Herb. x Sand", "Woody x Sand", "An./bn. x SD t. [ref.]", "Herb. x SD t.", "Woody x SD t.")
mxH.coef$num <- seq(9,1,-1)
mxH.coef$ub <- with(mxH.coef, as.vector(Estimate+se))
mxH.coef$lb <- with(mxH.coef, as.vector(Estimate-se))

### mean height - pop this into the supplement
mnH.lh <- with(lh.div, lm(CWM.mn.ht~(sd*lh)+(SAND*lh), na.action=na.pass))
mnH.dredge <- dredge(mnH.lh, beta="none", rank="AIC")
summary(model.avg(mnH.dredge, subset=delta<4, fit=TRUE))
#get.models(mnH.dredge, subset=TRUE)

# Plotting setup for mnH
mnH.coef <- coefTable(model.avg(mnH.dredge, subset=delta<4, fit=TRUE))
colnames(mnH.coef)[colnames(mnH.coef)=="Std. Error"] <- "se"
mnH.coef <- as.data.frame(mnH.coef[,1:2])
mnH.coef <- mnH.coef[c(1,2,3,7,8,9,4,5,6),]

mnH.coef$var <- c("An./bn. [ref.]", "Herb.", "Woody", "An./bn. x Sand [ref.]", "Herb. x Sand", "Woody x Sand", "An./bn. x SD t. [ref.]", "Herb. x SD t.", "Woody x SD t.")
mnH.coef$num <- seq(9,1,-1)
mnH.coef$ub <- with(mnH.coef, as.vector(Estimate+se))
mnH.coef$lb <- with(mnH.coef, as.vector(Estimate-se))

### functional dispersion
fdis.lh <- with(lh.div, lm(fdis~(mean*lh)+(CLAY*lh), na.action=na.pass)) 
fdis.dredge <- dredge(fdis.lh, beta="none", rank="AIC")
summary(model.avg(fdis.dredge, subset=delta<4, fit=TRUE)) 
#get.models(fdis.dredge, subset = TRUE)

# Plotting setup for Fdis
fdis.coef <- coefTable(model.avg(fdis.dredge, subset=delta<4, fit=TRUE))
colnames(fdis.coef)[colnames(fdis.coef)=="Std. Error"] <- "se"
fdis.coef <- as.data.frame(fdis.coef[,1:2])
fdis.coef <- fdis.coef[c(1,3,4,2,6,7,5,8,9),]

fdis.coef$var <- c("An./bn. [ref.]", "Herb.", "Woody", "An./bn. x Clay [ref.]", "Herb. x Clay", "Woody x Clay", "An./bn. x Mean t. [ref.]", "Herb. x Mean t.", "Woody x Mean t.")
fdis.coef$num <- seq(9,1,-1)
fdis.coef$ub <- with(fdis.coef, as.vector(Estimate+se))
fdis.coef$lb <- with(fdis.coef, as.vector(Estimate-se))

########################
# transparent colors for this plot
t.brown <- rgb(84, 48, 5, max = 255, alpha = 170, names = "t.brown")
t.blue <- rgb(53, 151, 143, max = 255, alpha = 170, names = "t.blue")
t.green <- rgb(127, 188, 65, max = 255, alpha = 170, names = "t.green")

#### PLOTTING LIFE HISTORY AFFECTS
jpeg("./analysis/figures/fdiv-env-core18-wLH.jpeg", width=6, height=7.5, unit="in",res=300)
par(mfrow=c(3,2))
par(oma=c(1.5,1.2,1.5,1.2))

###### LA plots #####

# LA - plotted by life history
par(mar=c(4,5,1.5,2.5))
with(unscaled.lh.div, plot(log(CWM.LA)~mean, pch=19, cex=0.9, col=ifelse(lh=="wood","#543005AA", ifelse(lh=="herb", "#35978FAA","#7FBC41AA")), 
                     xlab=expression(paste('Mean temp. (',degree,'C)')), ylab=expression(italic("ln")("Leaf area")), 
                     axes=FALSE, xlim=c(4,14)))
axis(1, cex.lab=1.5)
axis(2, cex.lab=1.5, at=c(-4,-2,0,2,4))
legend(11.5,4.5, legend=c('An./bn.','Herb.','Woody'), pch=16, cex=0.7, col = c("#7FBC41","#35978F","#543005"))
mtext(expression(bold("(a)")), side=3, line=-1.3, adj=-0.35, cex = 0.85)

# Relative effect plot for LA
par(mar=c(4,8,1.5,2))
with(la.coef, plot(num~Estimate, pch=19, xlim=c(-1.15, 1.5), xlab="Relative effect", ylab="", axes=FALSE, cex=1.2, col="#009E73"))
abline(v=0, lwd=2, lty=2, col="gray")
with(la.coef, segments(Estimate, num, ub, num, lwd=2, col="#009E73"))
with(la.coef, segments(Estimate, num, lb, num, lwd=2, col="#009E73"))
axis(1)
with(la.coef, axis(2, at=num, lwd.ticks=0,labels=var, las=1, cex.axis=0.8))
box()
mtext(expression(bold("(b)")),side=3, line=-1.3, adj=-0.8, cex = 0.85)

###### Max. height plots #####
# Max. height - plotted by life history
par(mar=c(4,5,1.5,2.5))
with(unscaled.lh.div, plot(log(CWM.maxht)~sd, pch=19, cex=0.9, col=ifelse(lh=="wood","#543005AA", ifelse(lh=="herb", "#35978FAA","#7FBC41AA")), 
                           xlab=expression(paste('SD temp. (',degree,'C)')), ylab=expression(italic("ln")("Max. Height")), 
                           axes=FALSE, xlim=c(5,15)))
axis(1, cex.lab=1.5, at=c(5,7,9,11,13,15))
axis(2, cex.lab=1.5)
mtext(expression(bold("(c)")),side=3, line=-1.3, adj=-0.35, cex = 0.85)

# Relative effect plot for mxH
par(mar=c(4,8,1.5,2))
with(mxH.coef, plot(num~Estimate, pch=19, xlab="Relative effect", ylab="", xlim=c(-0.65, 1.75), axes=FALSE, cex=1.2, col="#E69F00"))
abline(v=0, lwd=2, lty=2, col="gray")
with(mxH.coef, segments(Estimate, num, ub, num, lwd=2, col="#E69F00"))
with(mxH.coef, segments(Estimate, num, lb, num, lwd=2, col="#E69F00"))
axis(1)
with(mxH.coef, axis(2, at=num, lwd.ticks=0,labels=var, las=1, cex.axis=0.8))
box()
mtext(expression(bold("(d)")),side=3, line=-1.3, adj=-0.8, cex = 0.85)

###### Fdis plots #####
# Fdis - plotted by life history
par(mar=c(4,5,1.5,2.5))
with(unscaled.lh.div, plot(fdis~mean, pch=19, cex=0.9, col=ifelse(lh=="wood","#543005AA", ifelse(lh=="herb", "#35978FAA","#7FBC41AA")), 
                           xlab=expression(paste('Mean temp. (',degree,'C)')), ylab="FDis", 
                           axes=FALSE, xlim=c(4,14)))
axis(1, cex.lab=1.5)
axis(2, cex.lab=1.5)
mtext(expression(bold("(e)")),side=3, line=-1.3, adj=-0.35, cex = 0.85)

# Relative effect plot for Fdis
par(mar=c(4,8,1.5,2))
with(fdis.coef, plot(num~Estimate, pch=19, xlab="Relative effect", ylab="", xlim=c(-1.7, 1),axes=FALSE, cex=1.2, col="#0072B2"))
abline(v=0, lwd=2, lty=2, col="gray")
with(fdis.coef, segments(Estimate, num, ub, num, lwd=2, col="#0072B2"))
with(fdis.coef, segments(Estimate, num, lb, num, lwd=2, col="#0072B2"))
axis(1, cex.axis=0.9)
with(fdis.coef, axis(2, at=num, lwd.ticks=0,labels=var, las=1, cex.axis=0.8))
box()
mtext(expression(bold("(f)")),side=3, line=-1.3, adj=-0.8, cex = 0.85)

dev.off()

##### SUPPLEMENT #####
### Mean height plots
jpeg("./analysis/figures/SUPP-fdiv-env-core18-wLH.jpeg", width=6, height=5, unit="in",res=300)
par(mfrow=c(2,2))
par(oma=c(1,1,1,1))

###### SLA plots #####
# SLA - plotted by life history
par(mar=c(4,5,1.5,2.5))
with(unscaled.lh.div, plot(log(CWM.SLA)~mean, pch=19, cex=0.9, col=ifelse(lh=="wood","#543005AA", ifelse(lh=="herb", "#35978FAA","#7FBC41AA")), 
                           xlab=expression(paste('Mean temp. (',degree,'C)')), ylab=expression(italic("ln")("Specific leaf area")), 
                           axes=FALSE, xlim=c(4,14), ylim=c(1,5.3), cex.lab=0.8))
axis(1, cex.axis=0.8)
axis(2, cex.axis=0.8)
legend(11,5.2, legend=c('An./bn.','Herb.','Woody'), pch=16, cex=0.5, col = c("#7FBC41","#35978F","#543005"))
mtext(expression(bold("(a)")),side=3, line=-0.85, adj=-0.5, cex = 0.75)

# Relative effect plot for SLA
par(mar=c(4,8,1.5,2))
with(sla.coef, plot(num~Estimate, pch=19, xlab="Relative effect", ylab="", xlim=c(-1.2, 0.4), axes=FALSE, cex=1.2, col="#009E73", , cex.lab=0.8))
abline(v=0, lwd=2, lty=2, col="gray")
with(sla.coef, segments(Estimate, num, ub, num, lwd=2, col="#009E73"))
with(sla.coef, segments(Estimate, num, lb, num, lwd=2, col="#009E73"))
axis(1, cex.axis=0.8)
with(sla.coef, axis(2, at=num, lwd.ticks=0,labels=var, las=1, cex.axis=0.8))
box()
mtext(expression(bold("(b)")),side=3, line=-0.85, adj=-1.3, cex = 0.75)

###### Mean. height plots #####
# Mean. height - plotted by life history
par(mar=c(4,5,1.5,2.5))
with(unscaled.lh.div, plot(log(CWM.mn.ht)~sd, pch=19, cex=0.9, col=ifelse(lh=="wood","#543005AA", ifelse(lh=="herb", "#35978FAA","#7FBC41AA")), 
                           xlab=expression(paste('SD temp. (',degree,'C)')), ylab=expression(italic("ln")("Mean height")), 
                           axes=FALSE, xlim=c(5,15), ylim=c(2,7), cex.lab=0.8))
axis(1, cex.axis=0.8, at=c(5,7,9,11,13,15))
axis(2, cex.axis=0.8)
mtext(expression(bold("(c)")),side=3, line=-0.85, adj=-0.5, cex = 0.75)

# Relative effect plot for mnH
par(mar=c(4,8,1.5,2))
with(mnH.coef, plot(num~Estimate, pch=19, xlab="Relative effect", ylab="", xlim=c(-0.8, 1.8), axes=FALSE, cex=1.2, col="#E69F00", cex.lab=0.8))
abline(v=0, lwd=2, lty=2, col="gray")
with(mnH.coef, segments(Estimate, num, ub, num, lwd=2, col="#E69F00"))
with(mnH.coef, segments(Estimate, num, lb, num, lwd=2, col="#E69F00"))
axis(1, cex.axis=0.8)
with(mnH.coef, axis(2, at=num, lwd.ticks=0,labels=var, las=1, cex.axis=0.8))
box()
mtext(expression(bold("(d)")), side=3, line=-0.85, adj=-1.3, cex = 0.75)

dev.off()
```
