## Supplement - data and analysis for "Variation in near-surface soil temperature drives plant assemblage insurance potential": traits_clean.R

```
### Clean & combine all trait data into one data set ###
### Elizabeth Simpson 2021-03-24 ######################

# Uncomment to set working directory
setwd("~/Documents/projects/functional_traits_rhf")

library("plyr")

#############
# FUNCTIONS #
#############
# Convert pixels to mm: pixels/(DPI:inch/pixels)*25.3mm/inch
pix2mm <- function(pixels, DPI){
  mm.out <- NA
  for(i in seq_along(pixels)){
    mm.out[i] <- (pixels[i]/DPI)*25.4
  }
  return(mm.out)
}

#############
# LOAD DATA #
#############

# Leaf area - old scanner - 300 dpi
LA.os <- rbind(read.delim("./analysis/stalkless-EGS/rhf_old_scan_just-mean/statistics.txt"),
               read.delim("./analysis/stalkless-EGS/rhf_old_scan_plus-sd/statistics.txt"),
               read.delim("./analysis/stalkless-EGS/rhf_old_scan_plus-sd-div-2/statistics.txt"),
               read.delim("./analysis/stalkless-EGS/rhf_old_scan_plus-sd-mult-1_3/statistics.txt"))

# Leaf area - new scanner - 600 dpi
LA.ns <- rbind(read.delim("./analysis/stalkless-EGS/rhf_new_scan_just-mean/statistics.txt"),
               read.delim("./analysis/stalkless-EGS/rhf_new_scan_plus-sd-div-2/statistics.txt"),
               read.delim("./analysis/stalkless-EGS/rhf_new_scan_plus-sd-mult-1_3/statistics.txt"))

# Leaf mass
mass18 <- read.csv("./raw_data/rhf_2018_traits_sla_leaf_mass.csv", as.is=TRUE)
mass19 <- read.csv("./raw_data/rhf_2019_traits_sla_leaf_mass.csv", as.is=TRUE)

# Height
ht18 <- read.csv("./raw_data/rhf_2018_traits_height.csv", as.is=TRUE)
ht19 <- read.csv("./raw_data/rhf_2019_traits_height.csv", as.is=TRUE)

# Collection notes
col.notes <- read.csv("./raw_data/trait-collection-notes.csv", as.is=TRUE)
# Filter collection notes to only those that processed the leaves and petioles properly
col.notes.Y <- col.notes[col.notes$process.leaves.pets=="Y",]

##############
# CLEAN DATA #
##############

### LEAF AREA
# Convert surface.area pixels to mm - convert any other columns of interest to mm too
LA.os$sa.mm <- pix2mm(LA.os$surface.area, 300)
LA.ns$sa.mm <- pix2mm(LA.ns$surface.area, 600)

# combine them
area.stat <- rbind(LA.os, LA.ns)

# Give each individual a unique identifier across rows of data (multiple scans), called summary name (sum.name)
# Note to self: Your naming strategy made this unnecessarily complicated.
area.stat$sum.name <- NA
for (i in seq_len(nrow(area.stat))){
  if (grepl("^[A-D]", area.stat$original.file[i])){
    area.stat$sum.name[i] <- substr(area.stat$original.file[i], start=1,stop=8)
  } else if (grepl("\\d{2}-\\d{4}-\\d{2}", area.stat$original.file[i])){
    area.stat$sum.name[i] <- substr(area.stat$original.file[i], start=1,stop=11)
  } else if (grepl("\\d{2}-\\d{4}-\\d", area.stat$original.file[i])){
    area.stat$sum.name[i] <- substr(area.stat$original.file[i], start=1,stop=10)
  } else if (grepl("\\d{4}-\\d{2}", area.stat$original.file[i])){
    area.stat$sum.name[i] <- substr(area.stat$original.file[i], start=1,stop=8)
  } else {
    area.stat$sum.name[i] <- substr(area.stat$original.file[i], start=1,stop=7)
  }
}

# Calculate the total leaf area for each individual based on sum.name
# Includes all leaves + scans for each individual
area.indv <- as.data.frame(with(area.stat, tapply(sa.mm, sum.name, sum)))
area.indv$indv <- row.names(area.indv)
colnames(area.indv) <- c("t_sa", "indv")

### LEAF MASS
# Combine 2018 + 2019 leaf mass data, columns: N_leaves, Species, Bag_label [=indv], Leaf_mass_g, Petiole_mass_g
odw <- rbind(mass18[,c(4:6,9:10)], mass19[,c(4:5,7,11:12)]) # oven dry weight

# Give each individual a unique code
odw$indv <- NA
for (i in seq_len(nrow(odw))){
  if (grepl("^[A-D]", odw$Bag_label[i])){
    odw$indv[i] <- substr(odw$Bag_label[i], start=1,stop=8)
  } else if (grepl("\\d{2}-\\d{4}-\\d{2}", odw$Bag_label[i])){
    odw$indv[i] <- substr(odw$Bag_label[i], start=1,stop=11)
  } else if (grepl("\\d{2}-\\d{4}-\\d", odw$Bag_label[i])){
    odw$indv[i] <- substr(odw$Bag_label[i], start=1,stop=10)
  } else if (grepl("\\d{4}-\\d{2}", odw$Bag_label[i])){
    odw$indv[i] <- substr(odw$Bag_label[i], start=1,stop=8)
  } else {
    odw$indv[i] <- substr(odw$Bag_label[i], start=1,stop=7)
  }
}

# Combine petiole and leaf mass into total mass
odw$t_mass <- NA
for(i in seq_len(nrow(odw))){
  odw$t_mass[i]<- with(odw, Leaf_mass_g[i]+Petiole_mass_g[i])
}

# Calculate number of leaves, total mass, and the odw for each individual (sometimes multiple rows in the mass data sheet (L5/RF for example))
N_leaves <- as.data.frame(with(odw, tapply(N_leaves, indv, sum)))
t_mass <- as.data.frame(with(odw, tapply(t_mass, indv, sum)))
species <- as.data.frame(with(odw, tapply(Species, indv, unique)))
odw.indv <- cbind(species, N_leaves, t_mass)
odw.indv$indv <- rownames(odw.indv)
colnames(odw.indv) <- c("species", "N_leaves", "t_mass", "indv")

### Combine all leaf data and CALCULATE SLA
sla.indv <- merge(odw.indv, area.indv, by.x="indv", by.y="indv")

sla.indv$i_mass <- NA
sla.indv$i_sa <- NA
sla.indv$i_sla <- NA

for(i in seq_len(nrow(sla.indv))){
  sla.indv$i_mass[i]<- with(sla.indv, t_mass[i]/N_leaves[i])
  sla.indv$i_sa[i] <- with(sla.indv, t_sa[i]/N_leaves[i])
  sla.indv$i_sla[i] <-with(sla.indv, i_sa[i]/i_mass[i])
}

# Filter to only be the best collection for each species
# need to remove letter off the end of each indv id to aggregate to species
sla.indv$col.id <- NA
for (i in seq_len(nrow(sla.indv))){
    sla.indv$col.id[i] <- substr(sla.indv$indv[i], start=1,stop=nchar(sla.indv$indv[i])-1)
}

# subset species by chosen best collection and good leaf scans from the collection notes
sla.indv.bc <- sla.indv[sla.indv$col.id %in% col.notes.Y$leaf_col_analyze,]

# Summarize leaf traits (mean and variation) for each species based on number of observations for each species
sla.mean <- as.data.frame(with(sla.indv.bc, tapply(i_sla, species, mean)))
la.mean <- as.data.frame(with(sla.indv.bc, tapply(i_sa, species, mean)))
odw.mean <- as.data.frame(with(sla.indv.bc, tapply(i_mass, species, mean)))
sla.sd <- as.data.frame(with(sla.indv.bc, tapply(i_sla, species, sd)))
la.sd <- as.data.frame(with(sla.indv.bc, tapply(i_sa, species, sd)))
odw.sd <- as.data.frame(with(sla.indv.bc, tapply(i_mass, species, sd)))
total.N <- as.data.frame(with(sla.indv.bc, tapply(N_leaves, species, sum)))
mean.N <- as.data.frame(with(sla.indv.bc, tapply(N_leaves, species, mean)))

leaf.traits <- cbind(sla.mean, la.mean, odw.mean, sla.sd, la.sd, odw.sd, total.N, mean.N)
colnames(leaf.traits) <- c("sla.m", "la.m", "odw.m", "sla.sd", "la.sd", "odw.sd", "N.total", "N.m")

# Calculate variation in traits based on total number of leaves measured
# (Instead of mean number of leaves collected from each individual)
leaf.traits$sla.se <- NA
leaf.traits$la.se <- NA
leaf.traits$odw.se <- NA

for(i in seq_len(nrow(leaf.traits))){
  leaf.traits$sla.se[i]<- with(leaf.traits, sla.sd[i]/sqrt(N.total[i]))
  leaf.traits$la.se[i] <- with(leaf.traits, la.sd[i]/sqrt(N.total[i]))
  leaf.traits$odw.se[i] <-with(leaf.traits, odw.sd[i]/sqrt(N.total[i]))
}

leaf.traits <- leaf.traits[,c(1:3,9:11)] # Cut out SD and N info
leaf.traits <- na.omit(leaf.traits) # 123 sp. total

### HEIGHT
# Making RHF height data long form - from Will Pearse, 2018-10-29 #
# Combining 2018-2019 height data - Elizabeth Simpson,  2019-07-31

# 2018 data
subset <- t(as.matrix(ht18[,-1:-7]))
heights <- as.numeric(subset)

# Expand the original data to match (dropping unneeded columns)
expanded.ht18 <- ht18[rep(1:nrow(ht18), each=25),]
expanded.ht18 <- expanded.ht18[,1:7]

# Match up the two
expanded.ht18$height <- heights
expanded.ht18 <- na.omit(expanded.ht18)
expanded.ht18 <- expanded.ht18[,-7]
rownames(expanded.ht18) <- seq_along(1:nrow(expanded.ht18))

# 2019 data (recorded in long form), match columns to 2018 data
ht19 <- ht19[,1:7]
colnames(ht19)[7] <- "height"

# put all height together 
all.height <- rbind(expanded.ht18, ht19)

#some height metrics
ht.max <- as.data.frame(with(all.height, tapply(height, Species, max)))
ht.mean <- as.data.frame(with(all.height, tapply(height, Species, mean)))
ht.sd <- as.data.frame(with(all.height, tapply(height, Species, sd)))
ht.N <- as.data.frame(table(all.height$Species))$Freq

ht.traits <- cbind(ht.max, ht.mean, ht.sd, ht.N)
colnames(ht.traits) <-c("ht.max", "ht.m", "ht.sd", "ht.N")

ht.traits$ht.se <- NA

for(i in seq_len(nrow(ht.traits))){
  ht.traits$ht.se[i]<- with(ht.traits, ht.sd[i]/sqrt(ht.N[i]))
}

ht.traits <- ht.traits[,c(1:2,5)] # remove sd and N info
ht.traits <- na.omit(ht.traits) # removes 7 species for 131 species total

##############################
### COMBINE & EXPORT DATA ###
#############################
leaf.traits$species <- rownames(leaf.traits)
ht.traits$species <- rownames(ht.traits)

all.traits <- merge(leaf.traits, ht.traits, by.x="species", by.y="species")
write.csv(all.traits, "./clean_data/clean_traits_rhf.csv")
```
