## Supplement - data and analysis for "Variation in near-surface soil temperature drives plant assemblage insurance potential": metadata_cat_traits_rhf.txt

species: Full Latin binomial of species observed. In the format "Genus species". If a place-holder unidentifid morphotype name, etc., this is written as-is but ALWAYS in lowercase. 

life_history: The life history strategy of this species. 
