## Supplement - data and analysis for "Variation in near-surface soil temperature drives plant assemblage insurance potential": metadata_rhf_2018_cover.txt

Date: International format (YYYY-MM-DD)

Recorder_1, Recorder_2, etc.: Name of person or multiple people, noted in addtional columns, who are observing and recording data. Everyone in the field party is an observer.

Plot_id: Code number of plot (originally called 'Site' in 2017), following triad format. Starting with outer-most triad and working in from there.

Quadrant: Which quadrant the cover recording was made in (using the 50cm x 50cm quadrant)---1: top-left quadrant; 2: top-right quadrant; 3: bottom-left quadrant; 4: bottom-right quadrant---when the recorder is at the bottom of the plot facing uphill

Species: Full Latin binomial of species observed. No common names or abbreviations. In the format "Genus species". If soil, rock, poo, or place-holder unidentifid morphotype name, etc., this is written as-is but ALWAYS in lowercase. Potential hybrids are identified to the species they match closest to. 

Cover: 0-100; the cover recorded for the "Species". No cover is recorded as 0. 0 is also recorded for species not rooted in the plot but overhanging the plot.

Notes: "overhang-" followed by a number denotes cover recorded for the "Species," (0-100) for species not rooted in, but overhanging the plot. If no overhanging cover, this column is left blank.
