## Supplement - data and analysis for "Variation in near-surface soil temperature drives plant assemblage insurance potential": metadata_rhf_2018_soil_texture.txt

date_processed: International format (YYYY-MM-DD). Date soil prepped for hydrometer analysis.

Plot_id: Code number of plot (originally called 'Site' in 2017), following triad format. Starting with outer-most triad and working in from there. If note added, was re-done elsewhere.

tin_weight: Tin mass in grams.

tin_air_dry_soil: Tin and air dried soil in grams.

tin_oven_dry_soil: Tin and oven dried soil in grams.

processor: Name of person who prepped the air dried soil for hydrometer readings. Elizabeth Simpson performed all hydrometer readings.

date_analyzed: International format (YYYY-MM-DD). Date that hydrometer analysis performed. 

hydro_calib: Hydrometer calibration; the hydrometer reading in a 50 g/L sodium hexametaphospate. solution.

temp_calib: Temperature reading associated with hydrometer calibration.

soil_wt: Mass of soil used to assess soil texture, in grams.

hydro_40: Soil texture hydrometer reading at 40 seconds.

temp_40: Temperature reading associated with soil texture hydrometer reading at 40 seconds.

hydro_120: Soil texture hydrometer reading at 120 minutes.

temp_120: Temperature reading associated with soil texture hydrometer reading at 120 minutes.
