## Supplement - data and analysis for "Variation in near-surface soil temperature drives plant assemblage insurance potential": metadata_rhf_2018_traits_height.txt

Date: International format (YYYY-MM-DD)

Recorder_1, Recorder_2, etc.: Name of person or multiple columns of
    recorders. Everyone in the field party is an observer.

Plot_id: Code number of plot following triad format. Starting with outer-most triad and working in from there.

Sp_num: Unique number associated with a given species in a given plot. Present for double checking species from cover and trait data. 

Species: Full Latin binomial of species observed. No common names or
    abbreviations - EVER. In the format "Genus species". Abiotic covers, species that trait data was not recorded for and debris identifiable to species not recorded here. If unidentifiable but height data was recorded, morphotype is lowercase

Notes: Notes and observations about species in plot. Includes information about phenological state, especially senescense, and number of individuals in the plot if >25. Can also include notes about height of different features of the plant (eg. basal vs. flowering stem leaves) and heights of nearby plants that were outside of the plot. 

i1-i25: Height of up to 25 individuals for each species in centimeters. If less than 25 individuals were present then less than 25 of these columns are filled. If height not recorded for a species it was too senesced (dried, curled leaves or leaves not present) to reasonably measure height. If height recorded but notes also read senesced, leaves were brown but present in complete enough form that a measure of height was taken.
