## Supplement - data and analysis for "Variation in near-surface soil temperature drives plant assemblage insurance potential": metadata_rhf_2018_traits_sla_leaf_mass.txt

Date_collected: International format (YYYY-MM-DD), date leaves were collected in the field and scanned in the lab, followed by 72 hours of drying.

Collector_1, Collector_2, etc.: Name of person or people (multiple columns) who collected leaves. Everyone in the field party is a collector.

N_leaves: number of leaves collected from each individual plant. 

Species: Full Latin binomial of the species that leaves were collected from. In the format "Genus species". Each row represents one individual of a species who's leaves were collected.

Bag_label: "Plot_id-sp_num-individual_lower_case_letter" Each individual leaves were collected from is labeled by the:
	Plot_id: Code number of plot following triad format. Starting with the outer-most triad and working in from there.
	sp_num: Unique number associated with a given species in a given plot. Used to associate leaf collections with species and associated scans.
	individual_lower_case_letter: Each individual of a given species was also given a lower case letter identifier a-e.

Date_weighed: International format (YYYY-MM-DD), date leaves were weighed in the lab.

Weigher: Name of person who weighed the leaves.

Leaf_mass_g: Leaf mass in grams.

Petiole_mass_g: Petiole mass if separated from leaf and weighed separately.

Scans: The filename of the scan(s) associated with each collection of leaves from an individual. Some scans included more than one individual; this scan name will be present for more than one individual. Some individuals required more than one scan; these are noted with capital letters in the scan name after the individual lower case letter. Scans with petiole include the abbreviation 'pet'.
