## Supplement - data and analysis for "Variation in near-surface soil temperature drives plant assemblage insurance potential": metadata_rhf_2019_traits_height.txt

Date: International format (YYYY-MM-DD)

Recorder_1, Recorder_2, etc.: Name of person or multiple columns of
    recorders. Everyone in the field party is an observer.

Plot_id: Code number of plot following triad format. Starting with outer-most triad and working in from there.

***lat: latitude if seeds likely collected >20m from plot
***long: longitude if seeds likely collected >20m frm plot

sp_num_indiv: Unique number associated with a given species in a given plot. Present for double checking species from cover and trait data.  ***when combined with plot id - this is a unique code for each collection***

Species: Full Latin binomial of species observed. No common names orabbreviations - EVER. In the format "Genus species". Abiotic covers, species that trait data was not recorded for and debris identifiable to species not recorded here. If unidentifiable but height data was recorded, morphotype is lowercase

seeds_individual: number of seeds collected per individual

notes: Notes and observations about seed collection if differs from 'all species easily collected', maturity and whether it seems like most seeds were on the plant when collected.
