## Supplement - data and analysis for "Variation in near-surface soil temperature drives plant assemblage insurance potential": metadata_trait_collection_notes.txt

species: Full Latin binomial of species observed. In the format "Genus species". If a place-holder unidentifid morphotype name, etc., this is written as-is but ALWAYS in lowercase. 

leaf_col_analyze: The identifier of the individual determined to be the best collection of leaves from each species. 

process-leaves-pets: Whether or not the scan from the best collection of leaves processessed well. "Y" = yes. "NO" = no.
