## Supplementary figures and images for "Variation in near-surface soil temperature drives plant assemblage insurance potential"

### 1111-2018-09-12.PNG

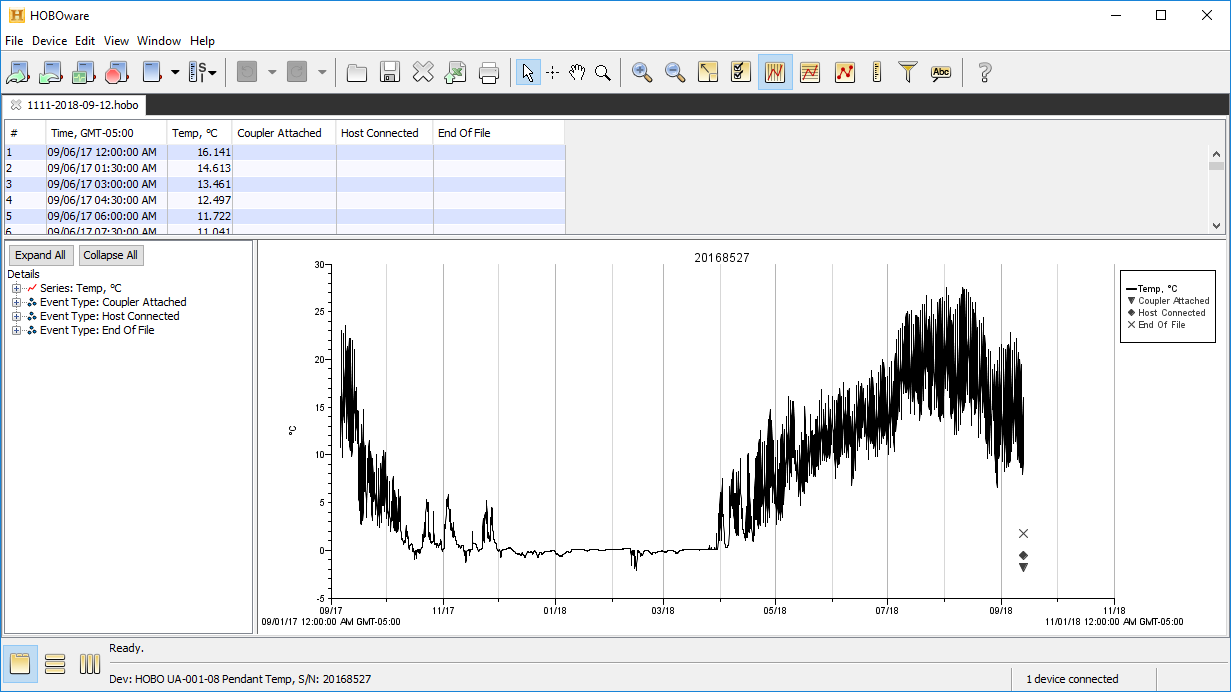

### 1111_2019_09_23_plot.png

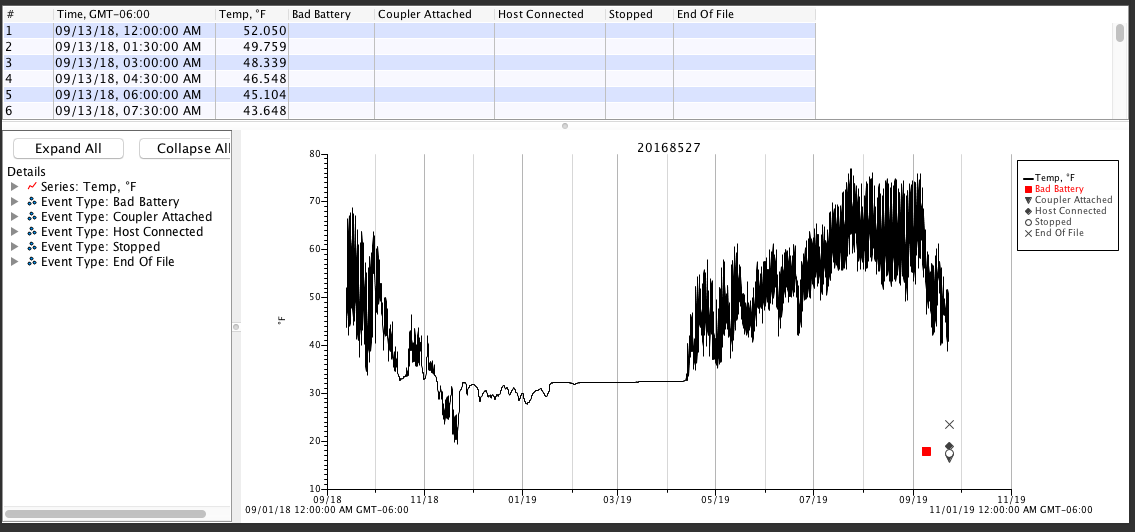

### 1111_2019_09_23_status.png

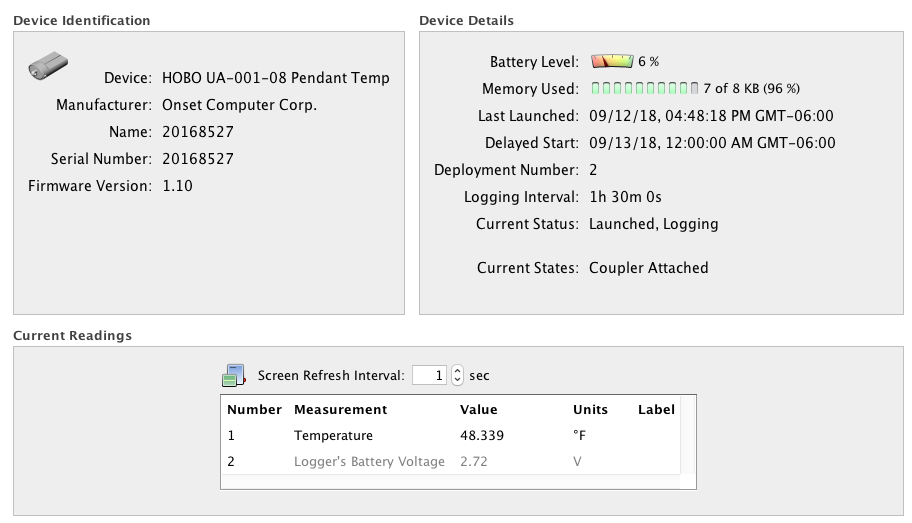

### 1122-2018-09-14.PNG

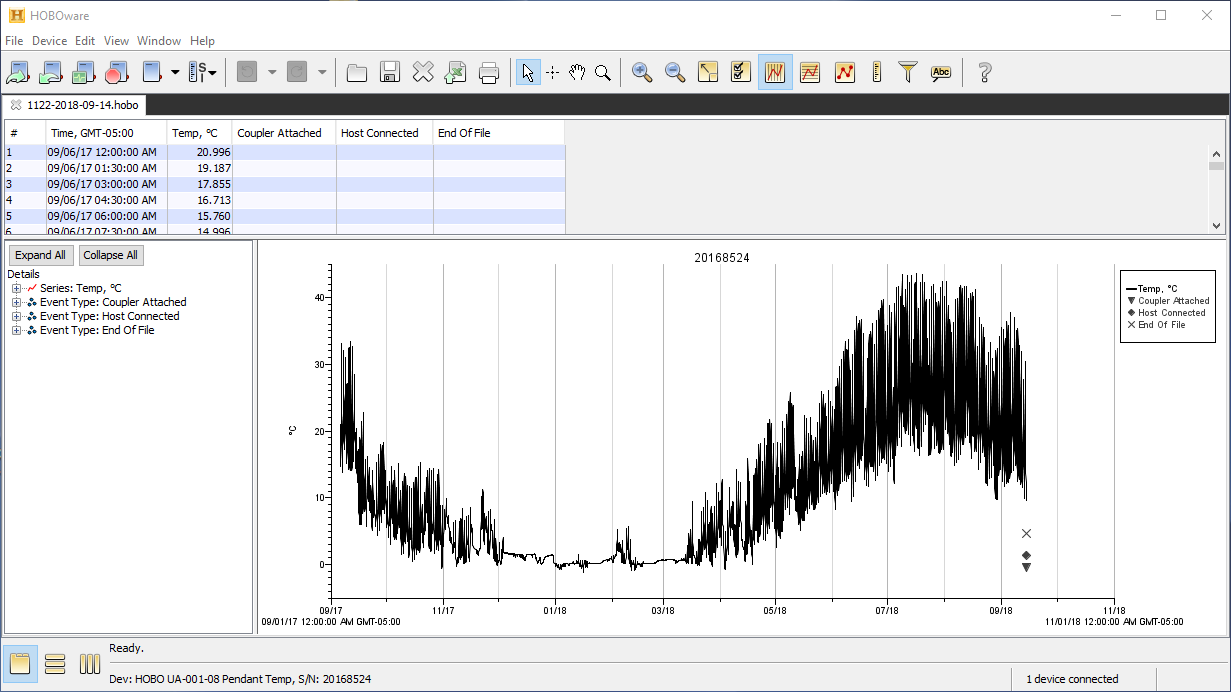

### 1122_2019_09_23_plot.png

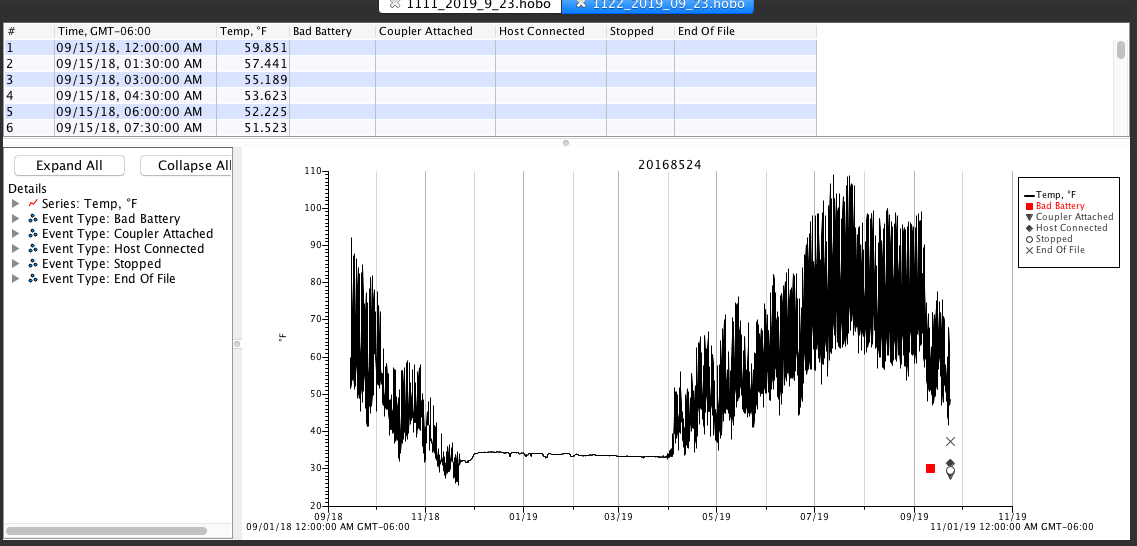

### 1122_2019_09_23_status.png

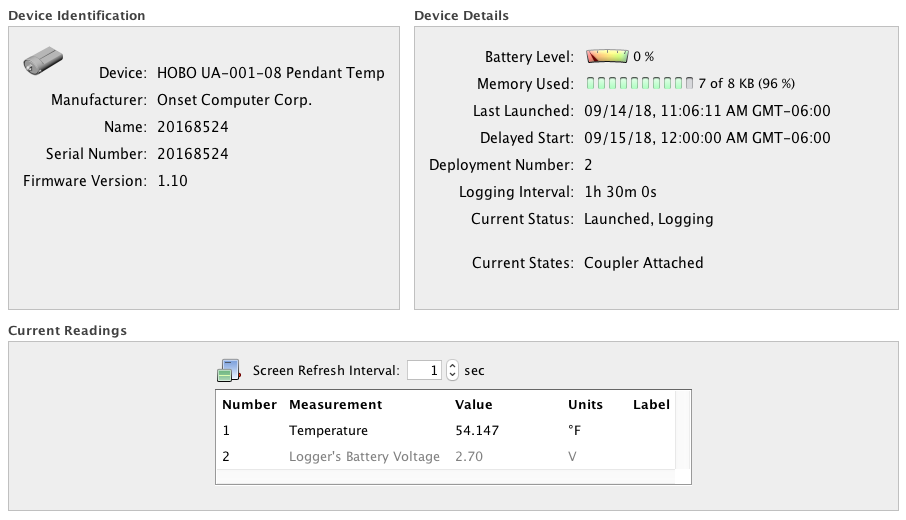

### 1133-2018-09-14.PNG

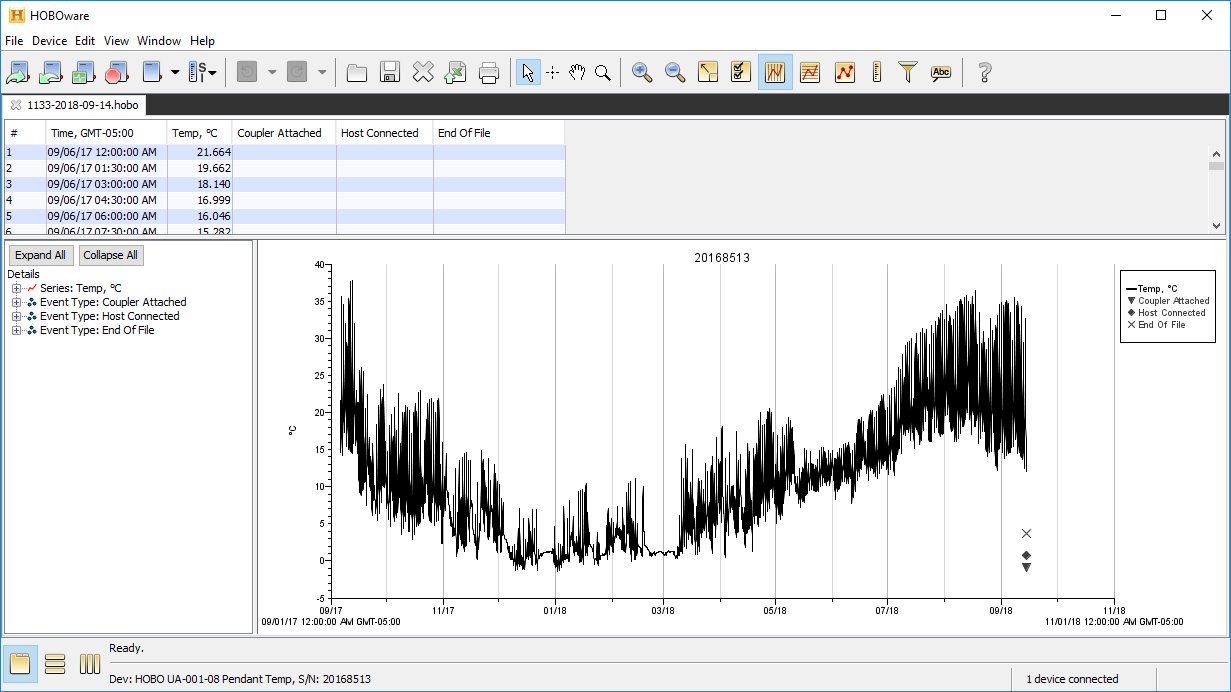

### 1133_2019_09_23_plot.png

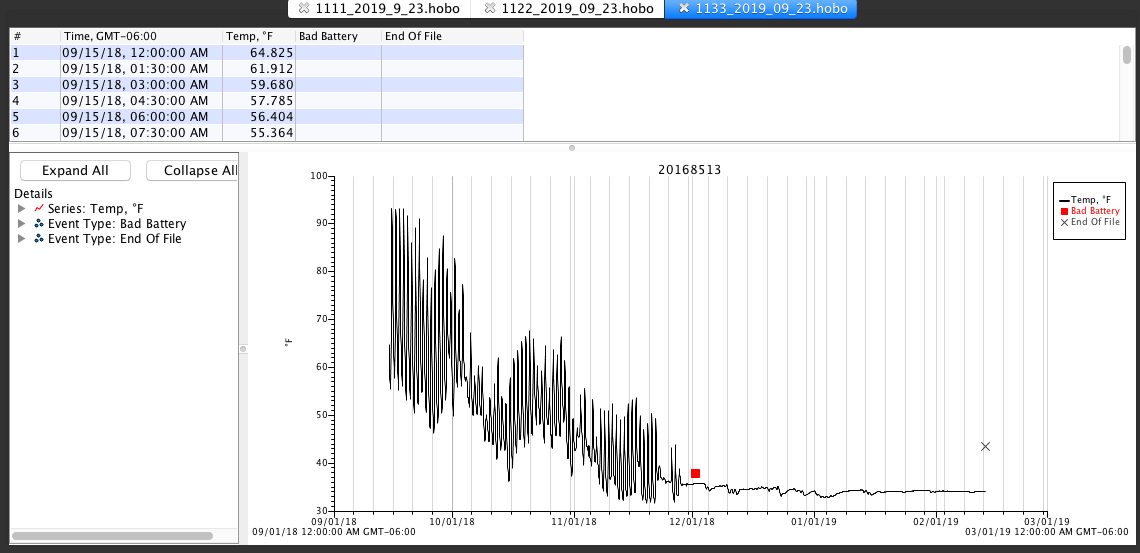

### 1133_2019_09_23_power_reset.png

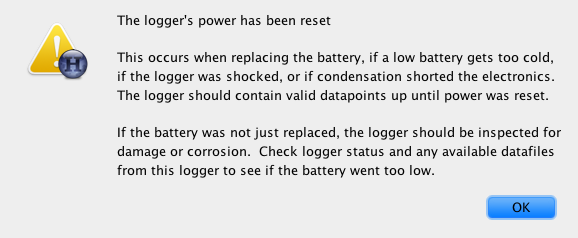

### 1133_2019_09_23_status.png

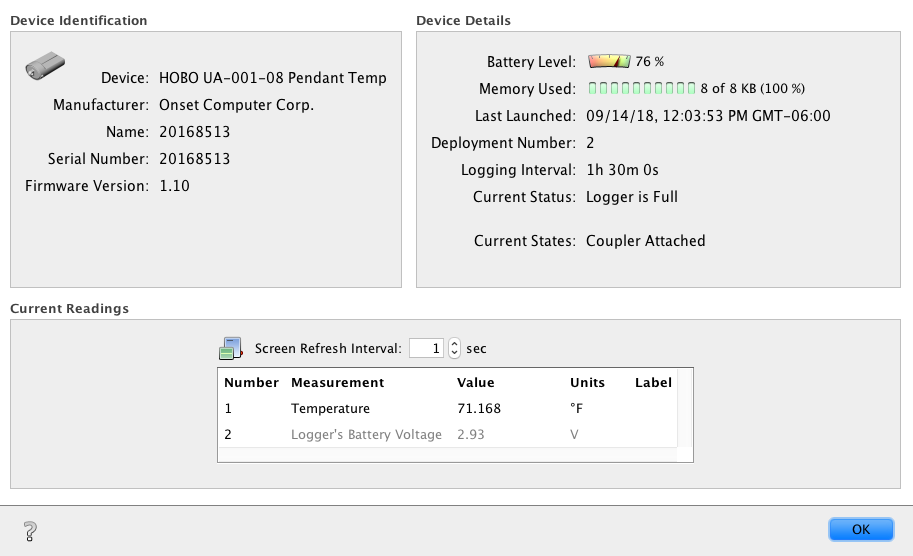

### 1211-2018-09-19.PNG

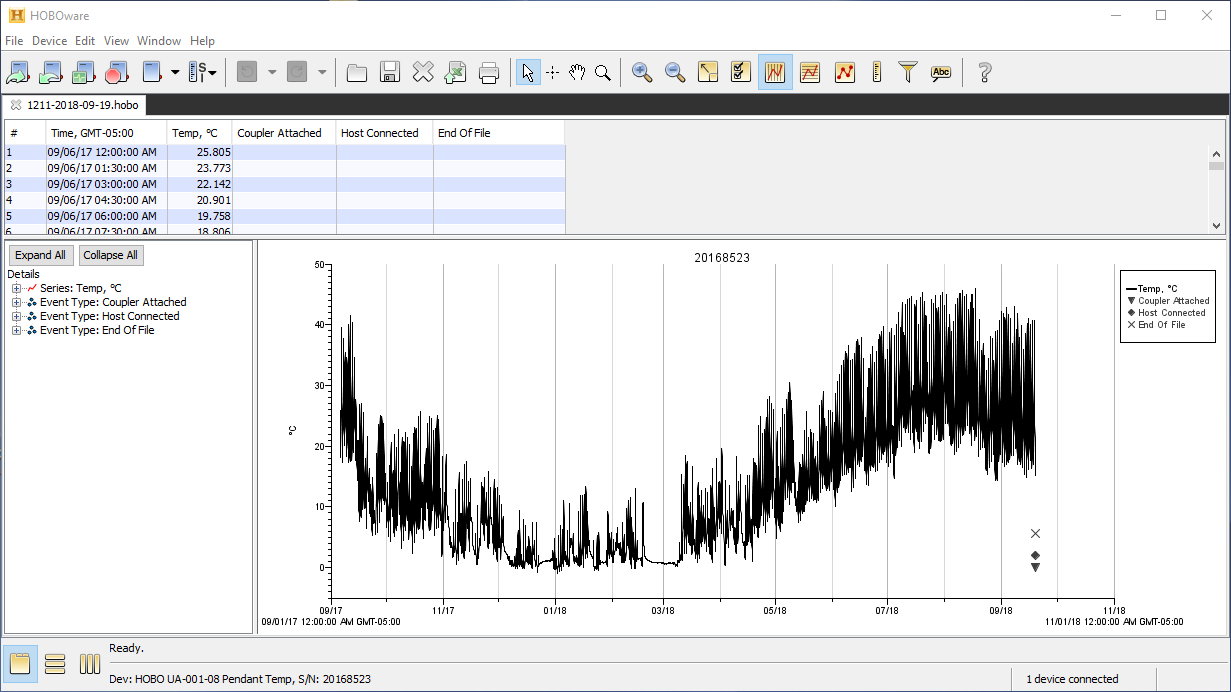

### 1222-2018-09-19.PNG

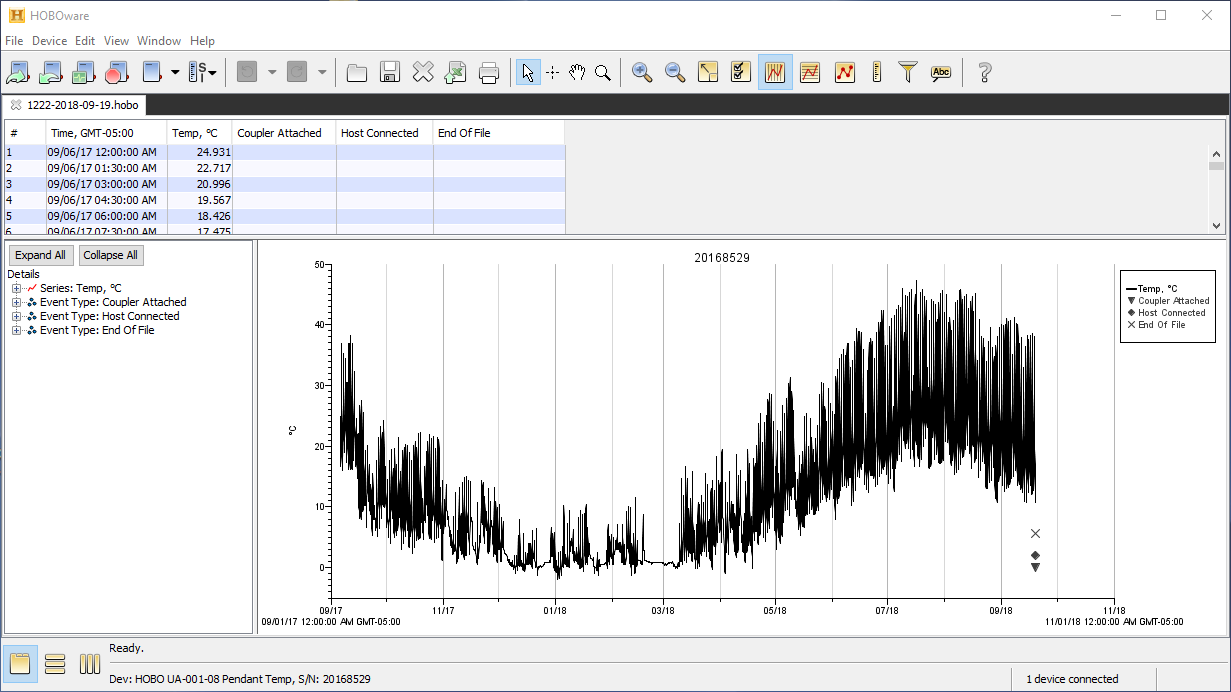

### 1222_2019_09_23_power_reset.png

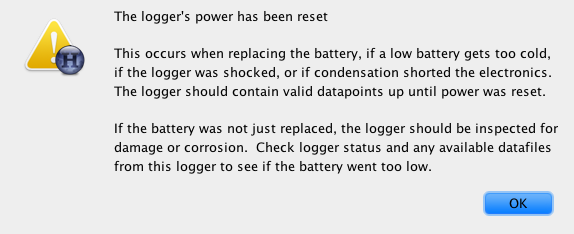

### 1233-2018-09-19.PNG

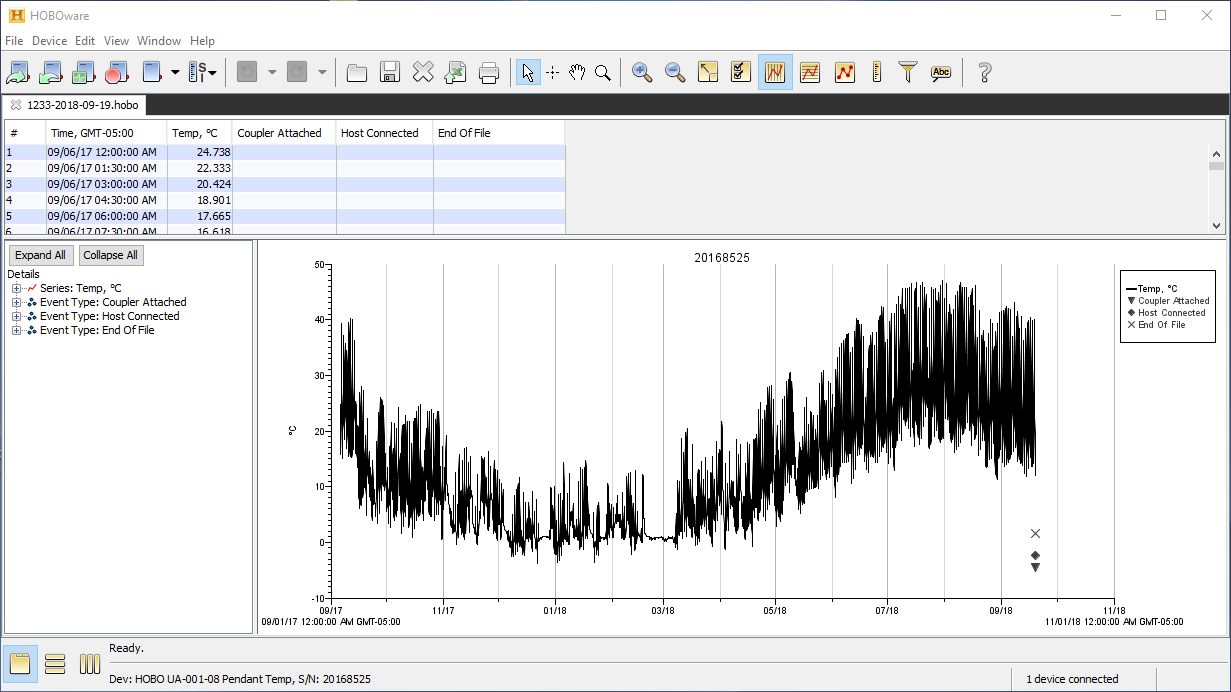

### 1233_2019_09_23_plot.png

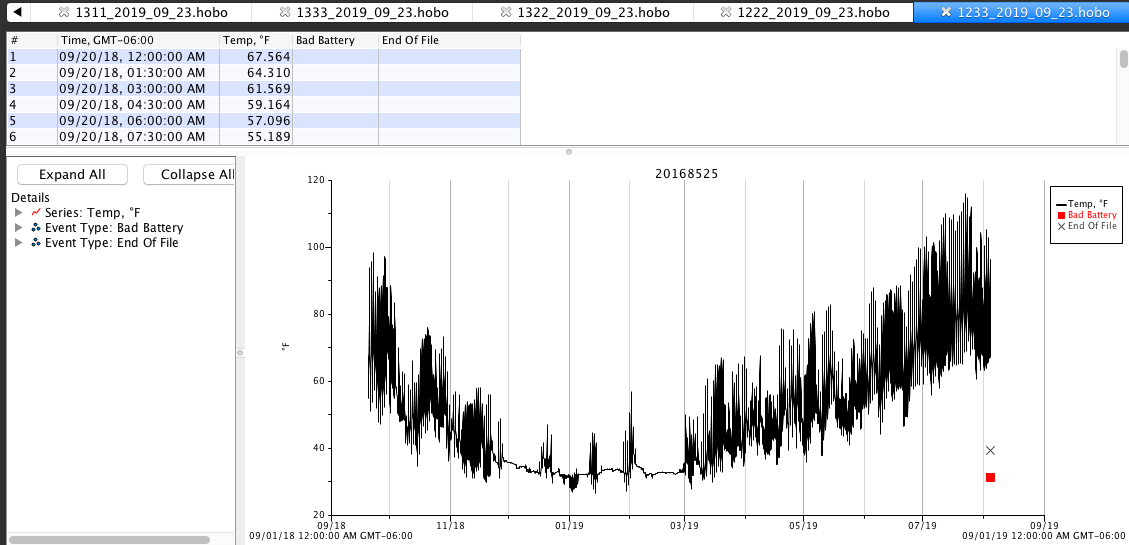

### 1233_2019_09_23_power_reset.png

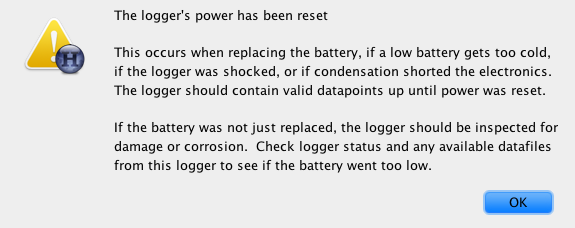

### 1233_2019_09_23_status.png

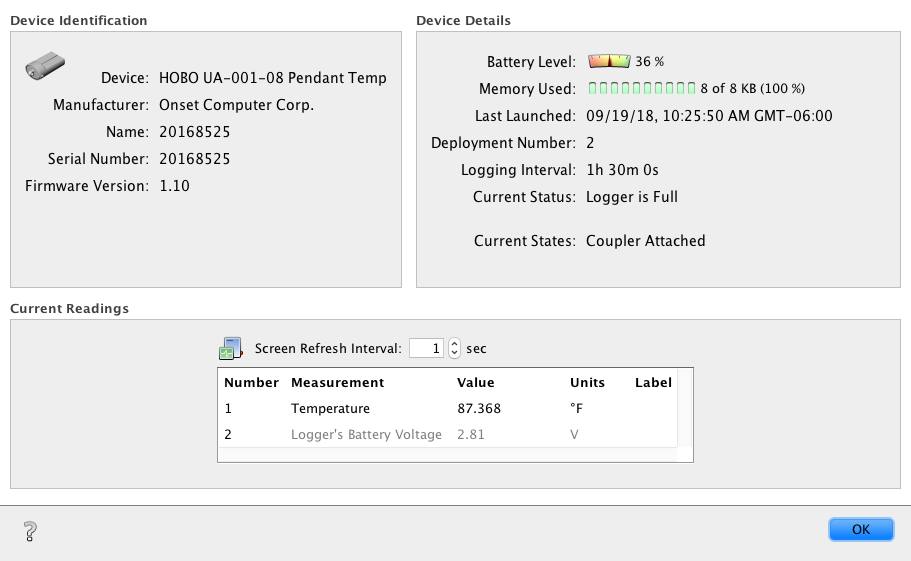

### 1311-2018-09-14.PNG

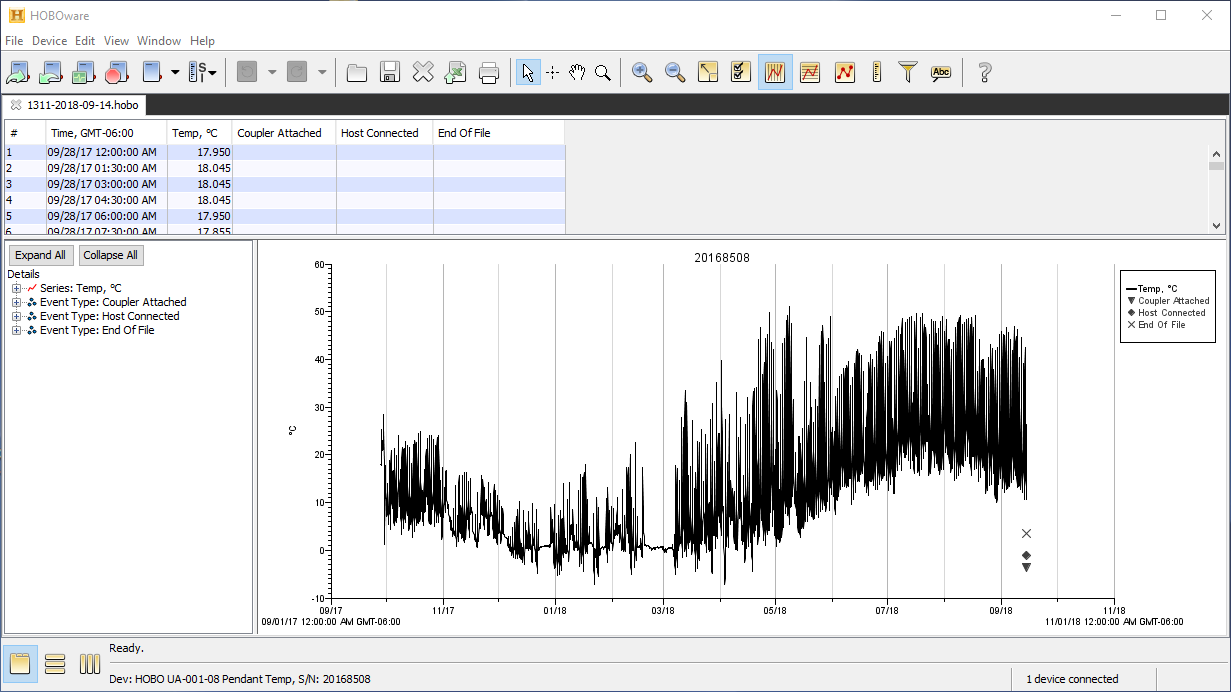

### 1311_2019_09_23_plot.png

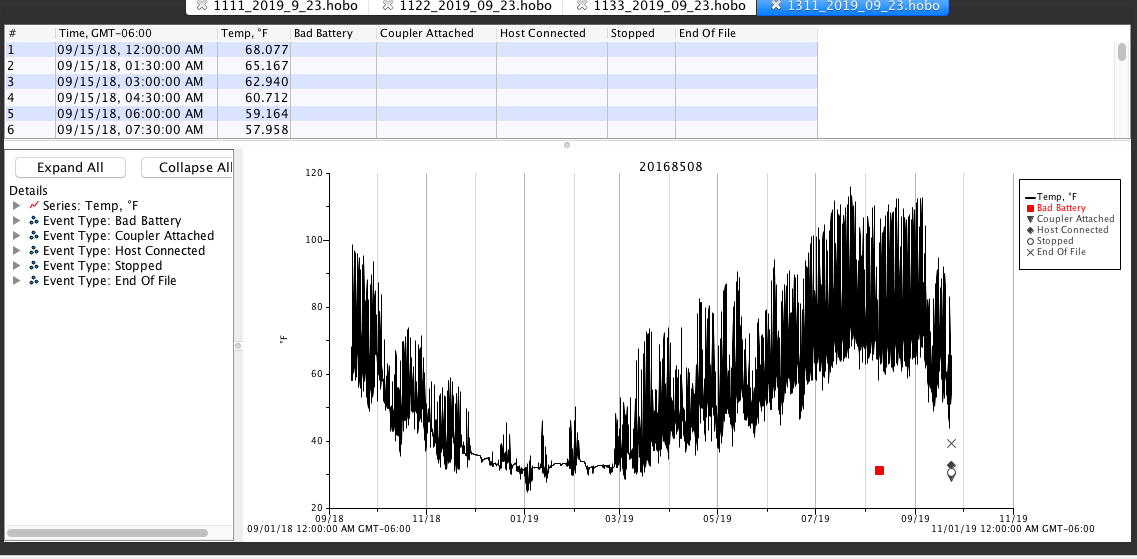

### 1311_2019_09_23_status.png

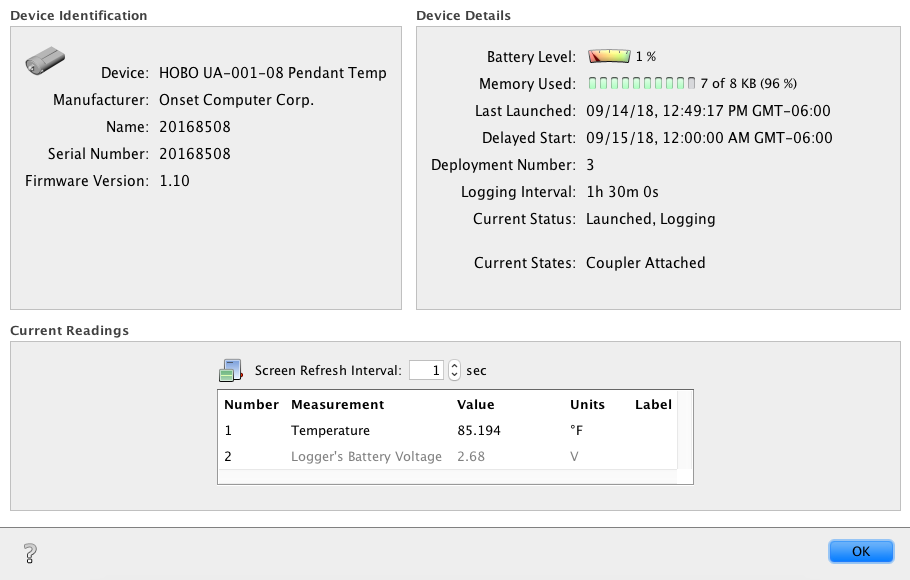

### 1322_2019_09_23_plot.png

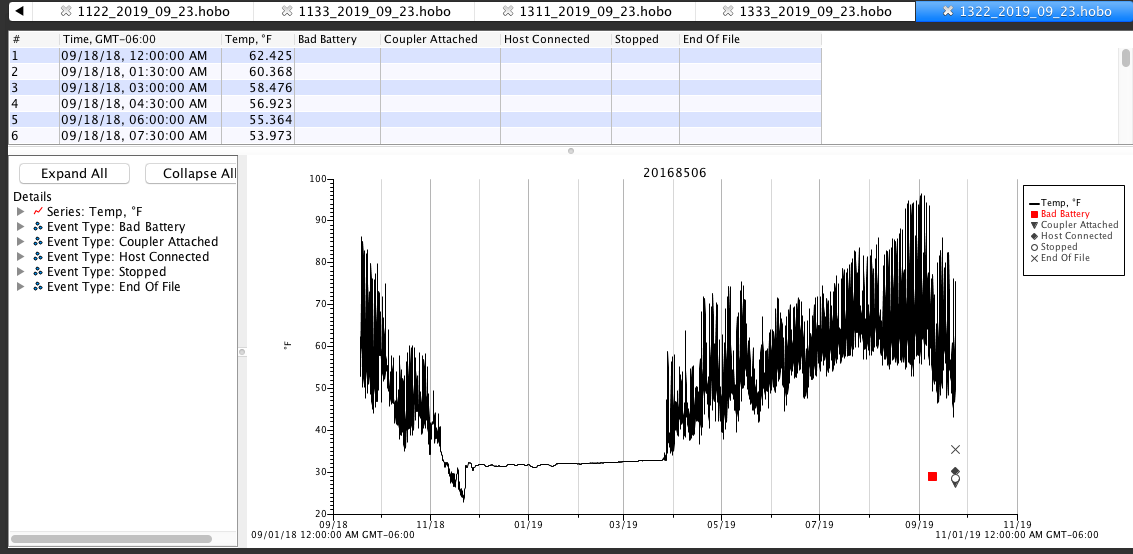

### 1322_2019_09_23_status.png

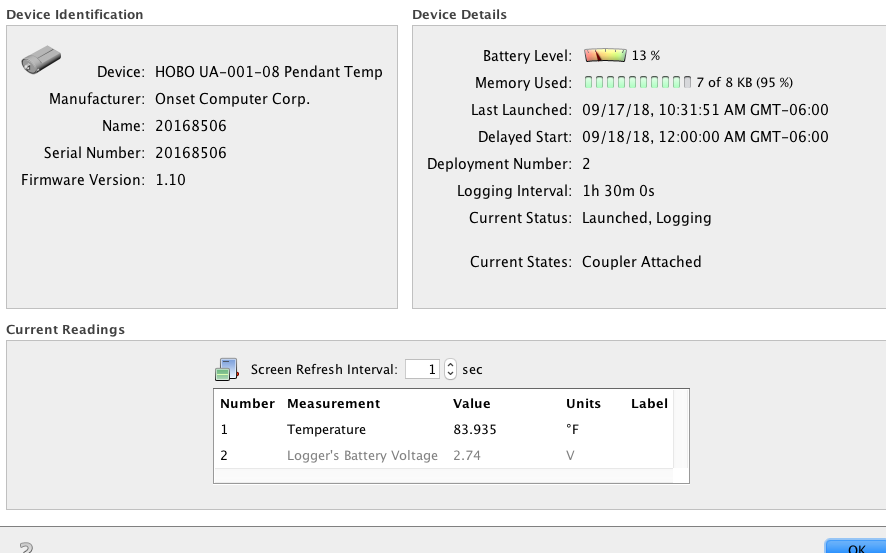

### 1333-2018-09-17.png

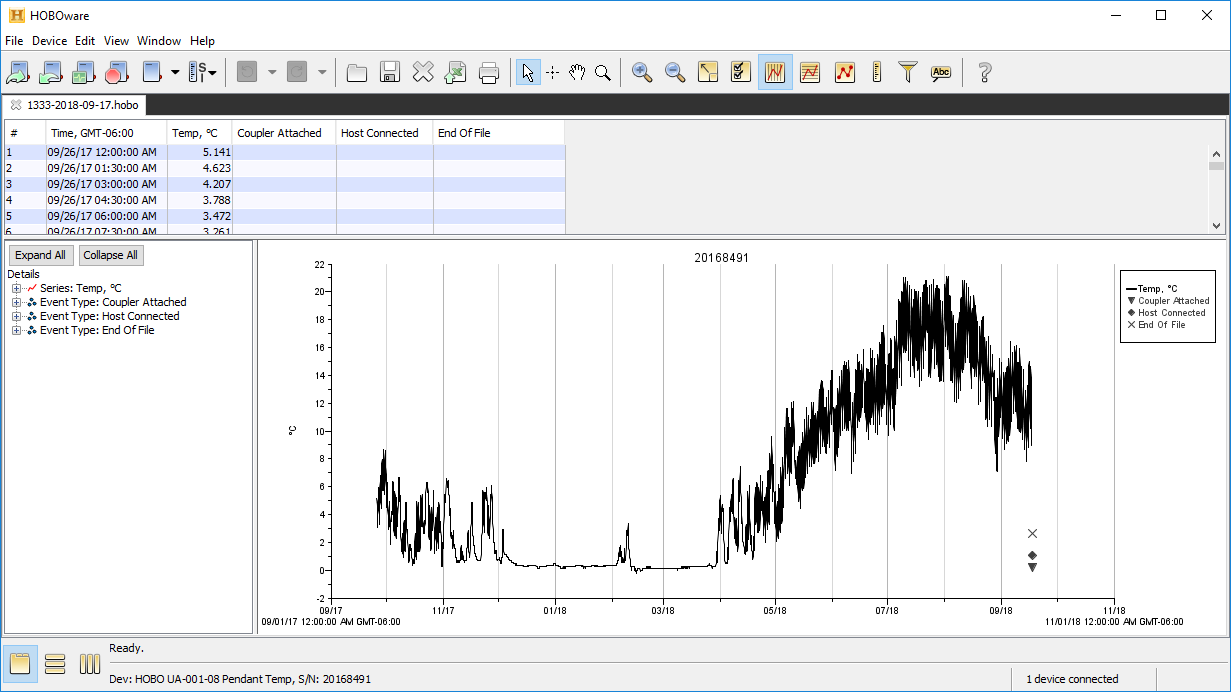

### 1333_2019_09_23_plot.png

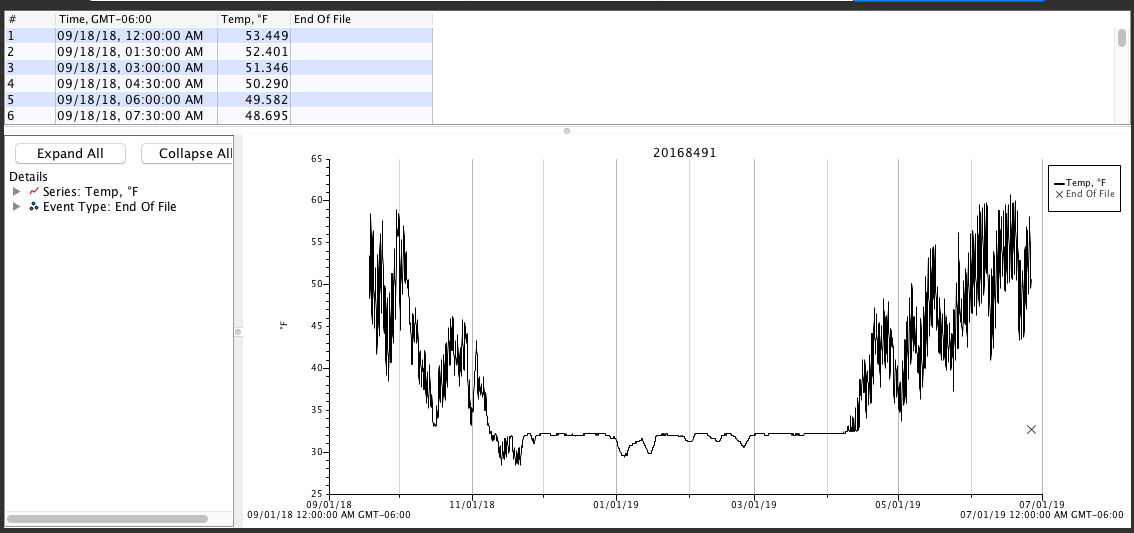

### 1333_2019_09_23_power_reset.png

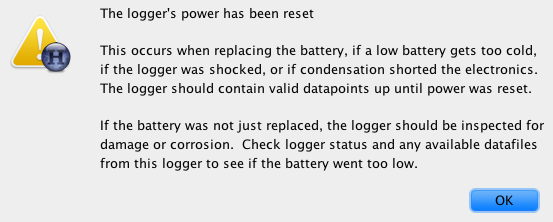

### 1333_2019_09_23_status.png

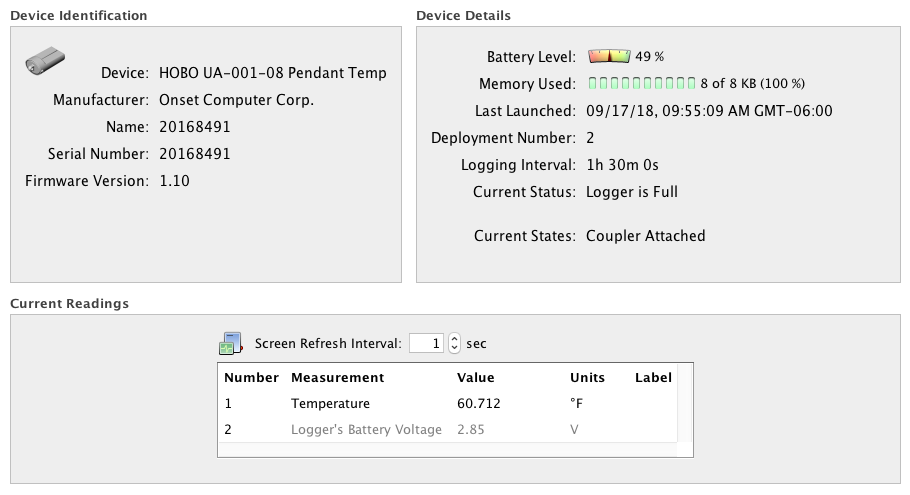

### 2111-2018-09-21.png

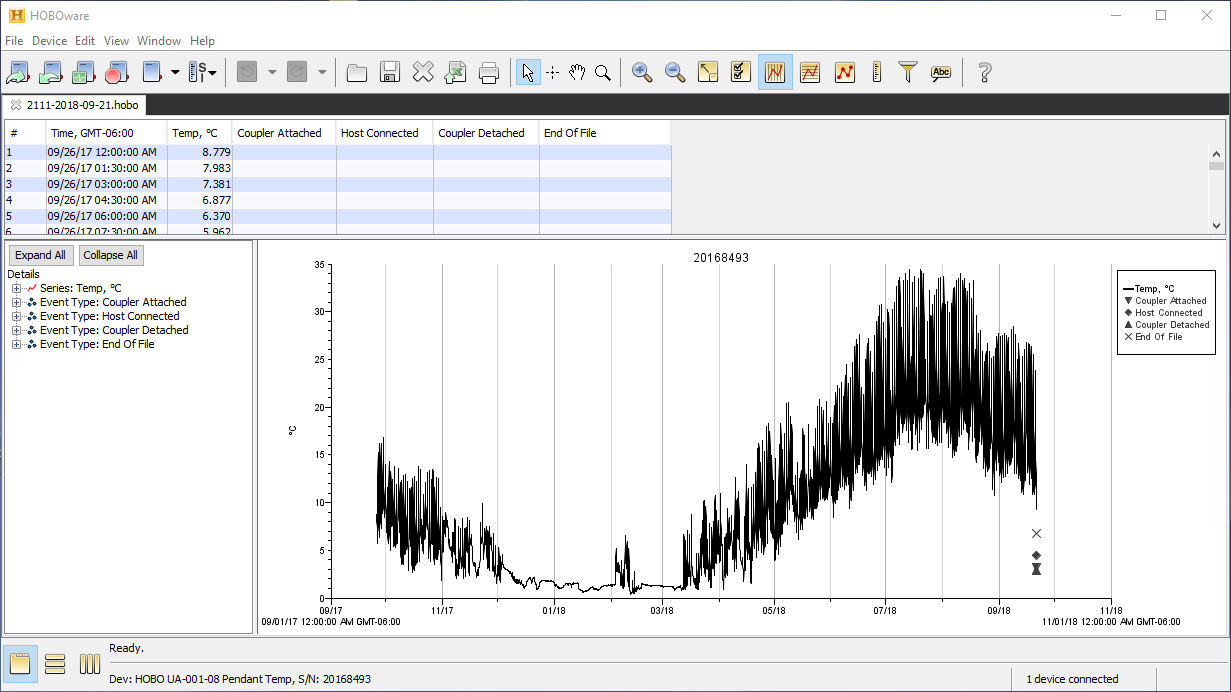

### 2111_2019_10_16.PNG

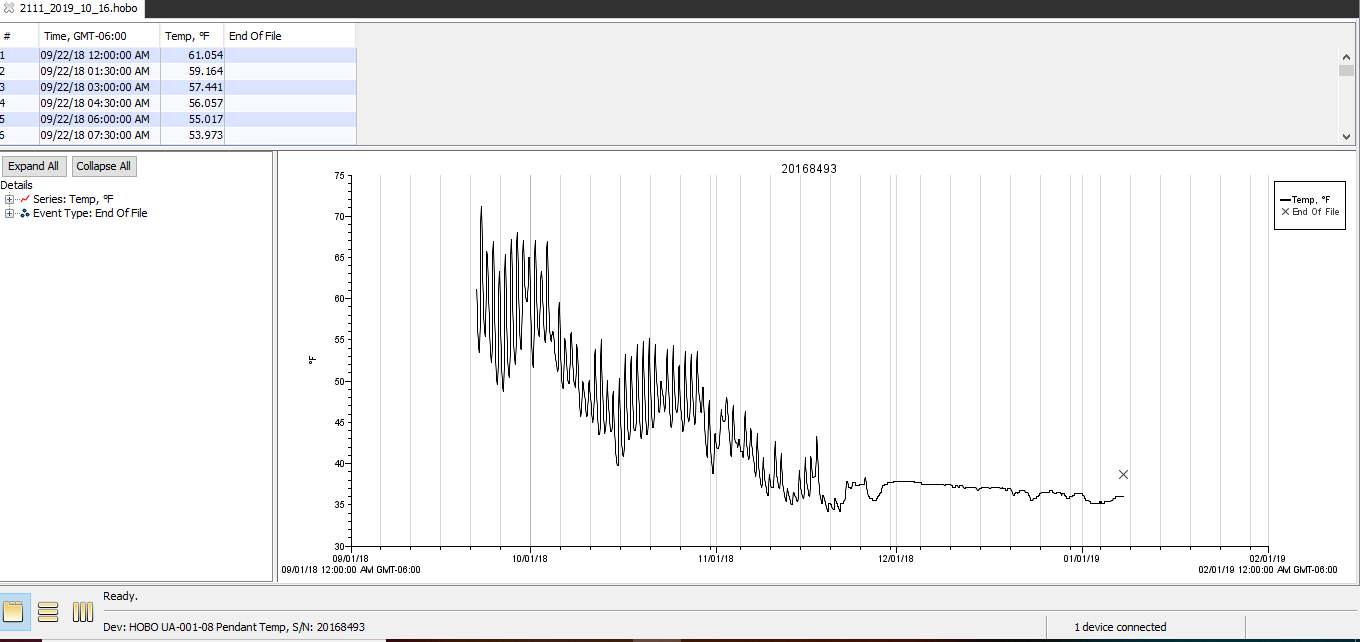

### 2122_2019-10-16_plot.PNG

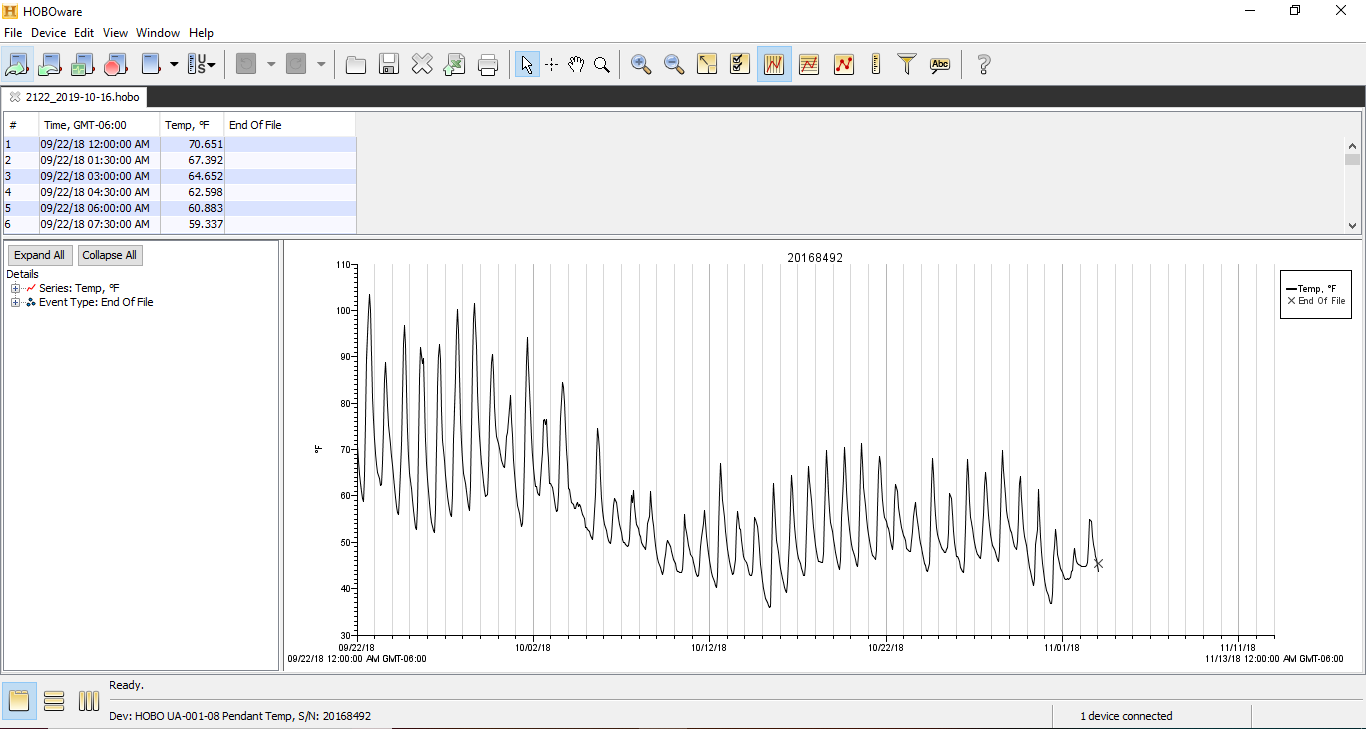

### 2122_2019-10-16_power-reset.PNG

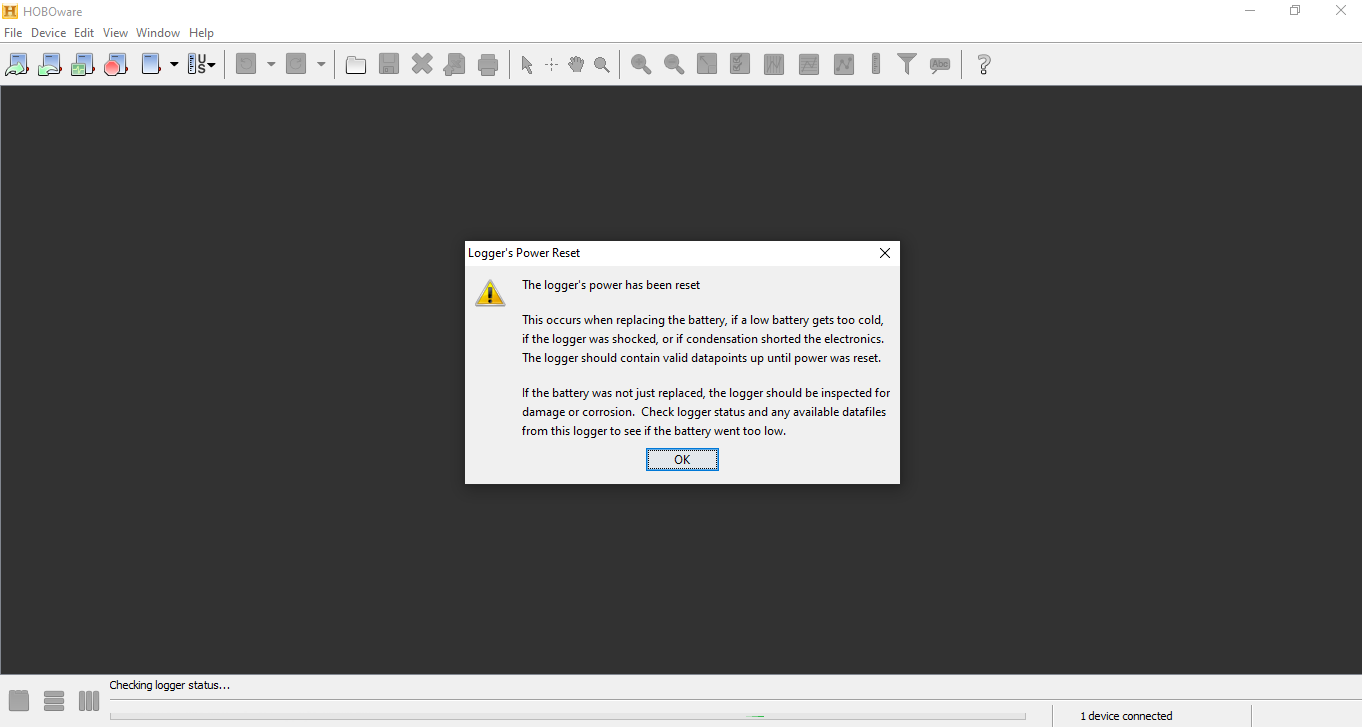

### 2122_2019-10-16_status.PNG

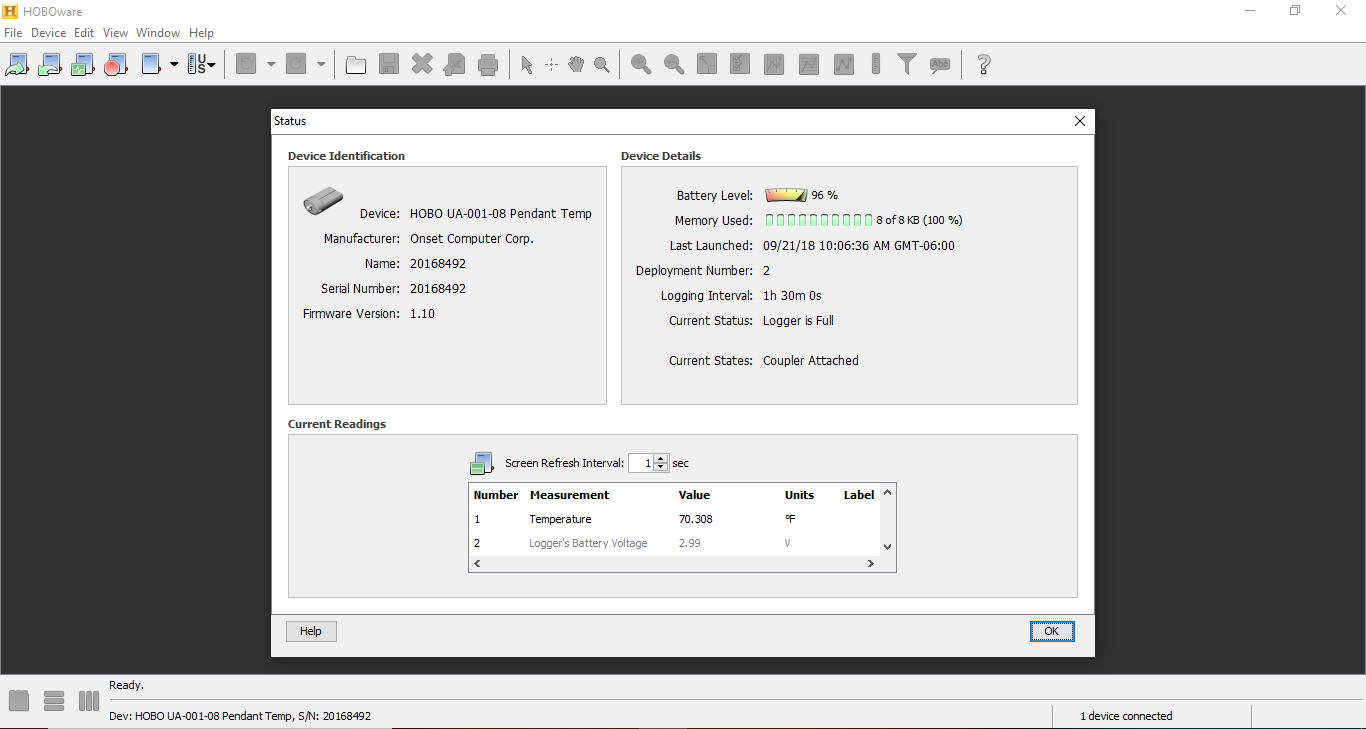

### 2133-2018-09-21.png

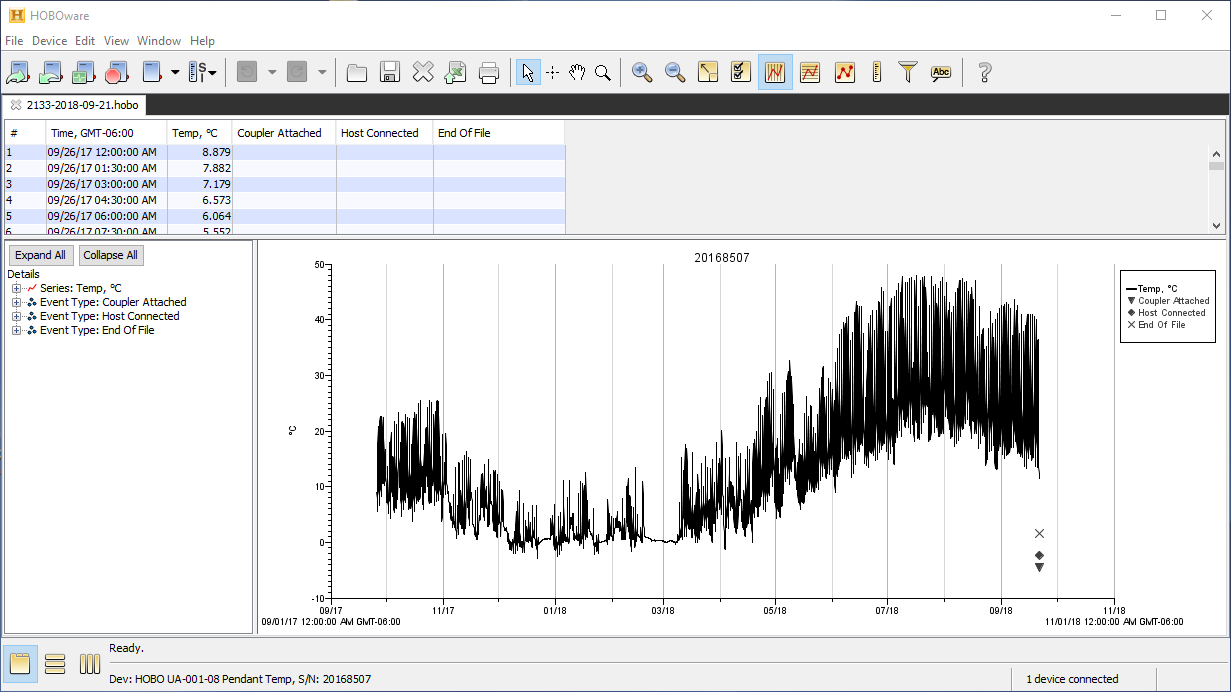

### 2133_2019-10-16_plot.PNG

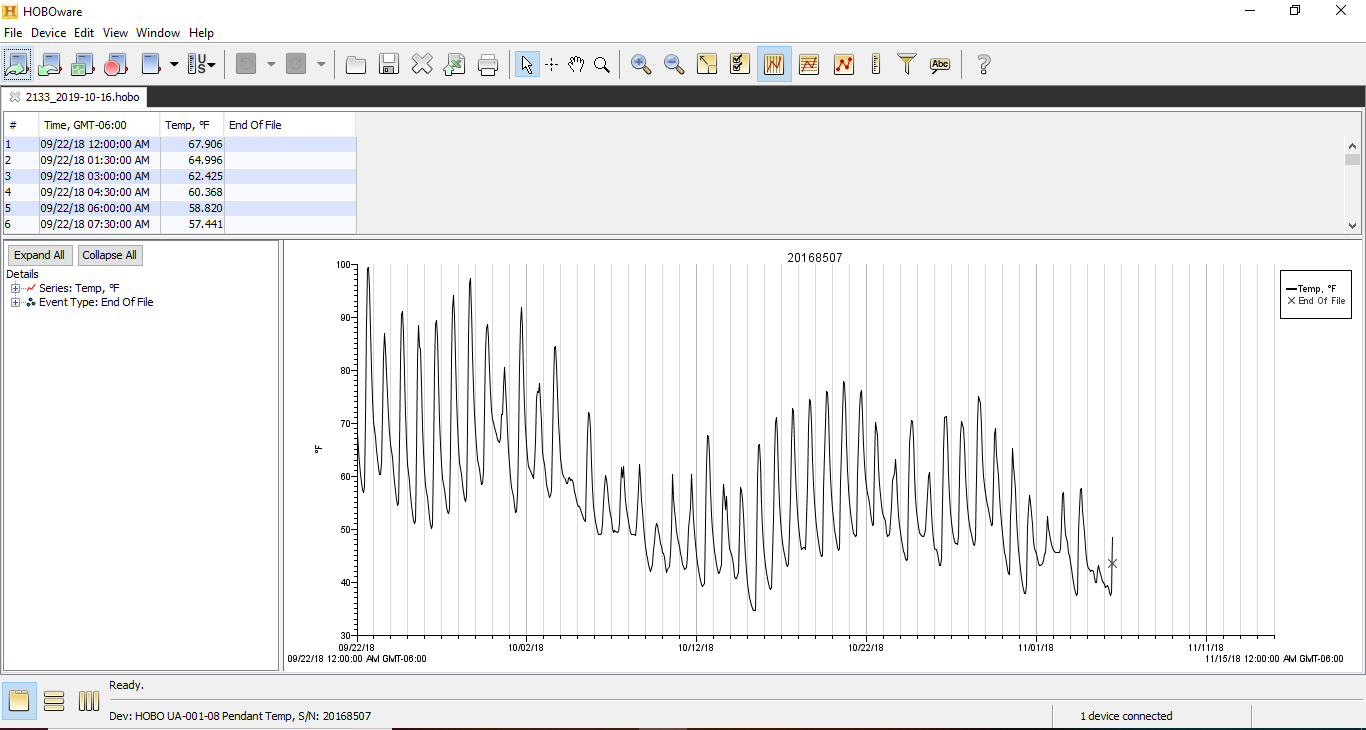

### 2133_2019-10-16_power-reset.PNG

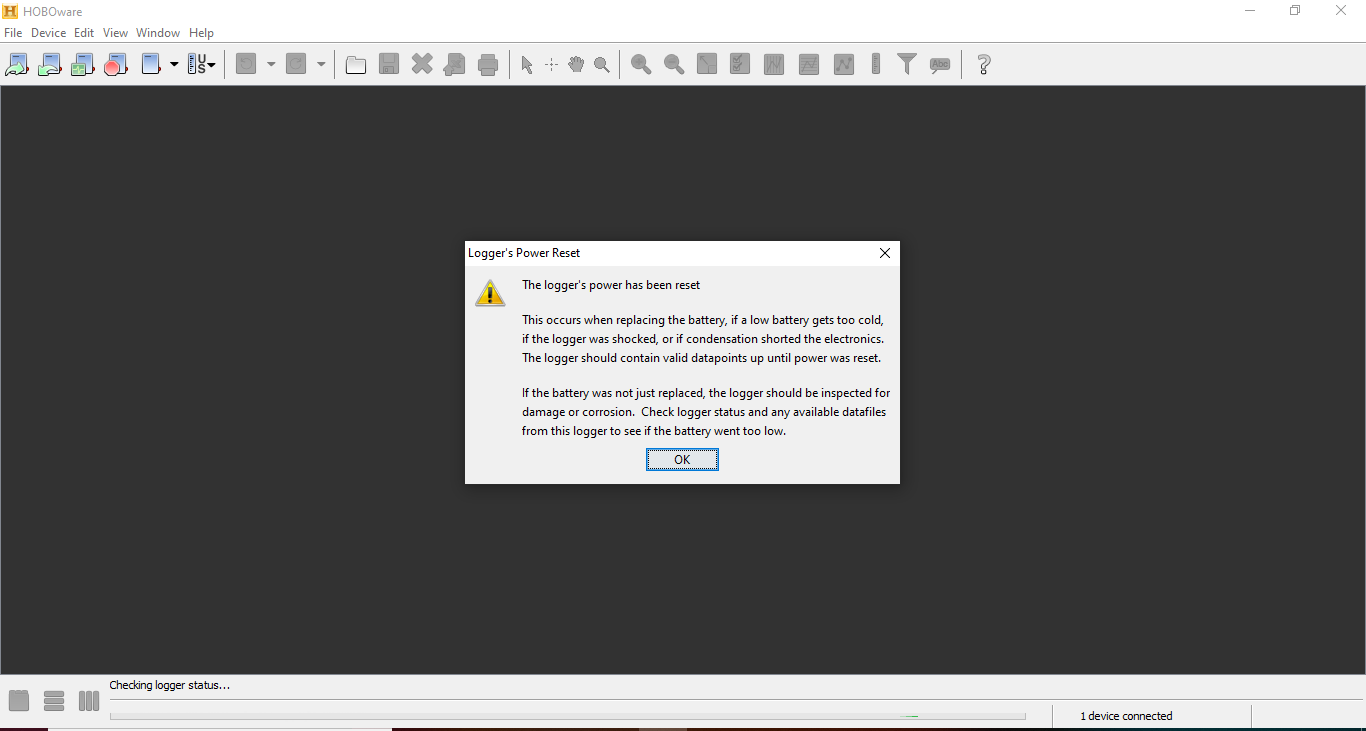

### 2133_2019-10-16_status.PNG

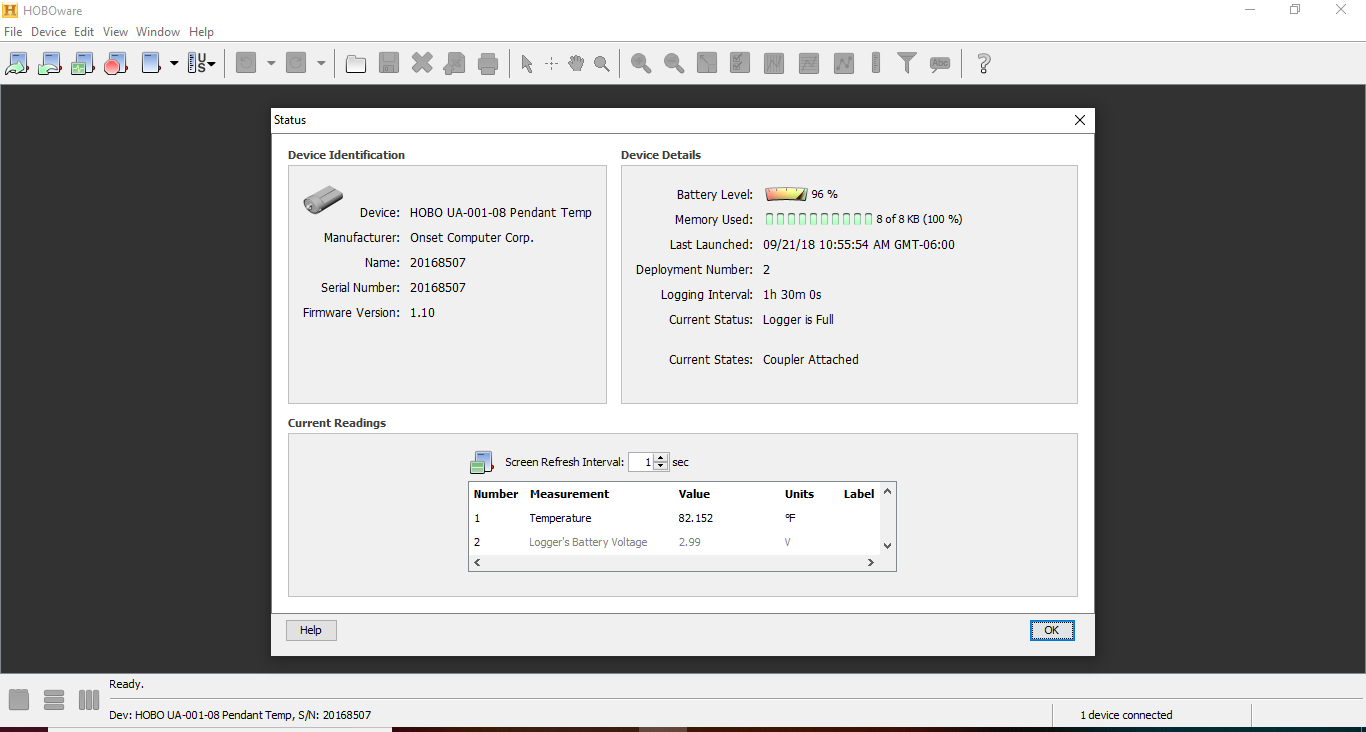

### 2211_2019-10-16_plot.PNG

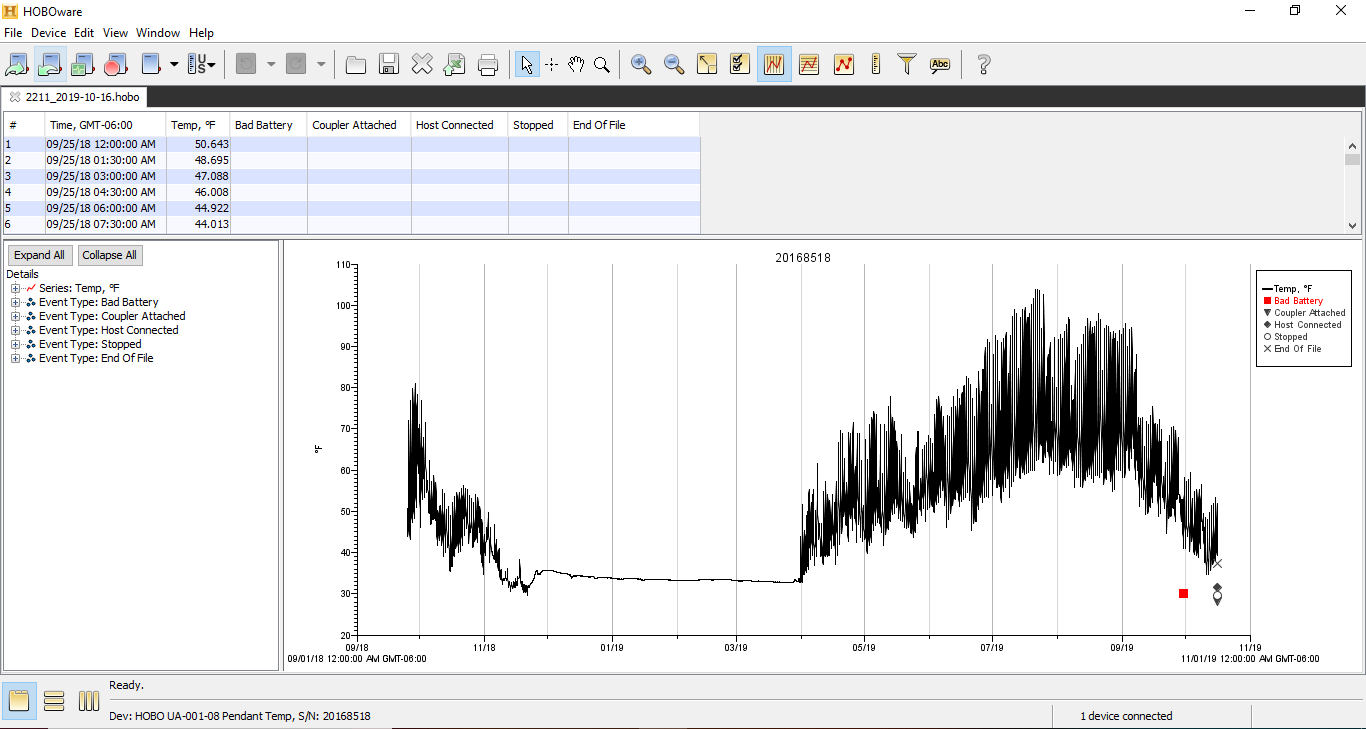

### 2211_2019-10-16_status.PNG

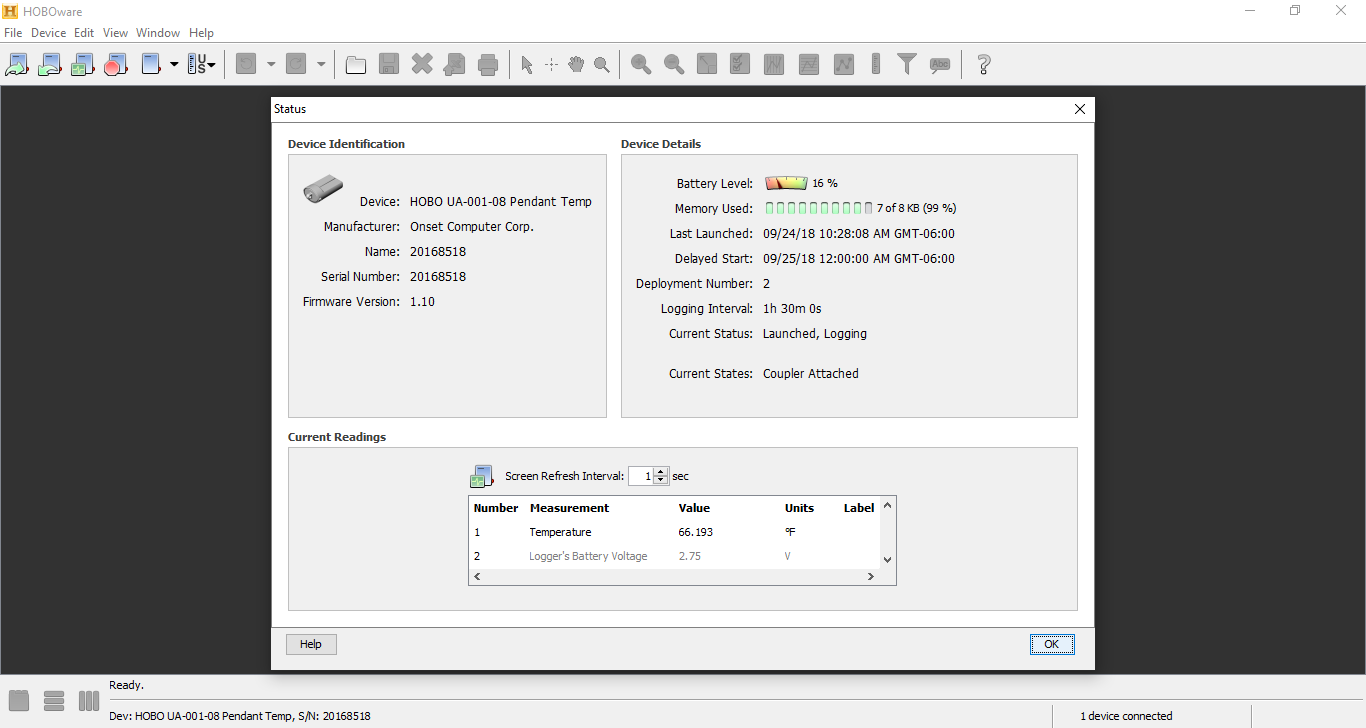

### 2222-2018-09-24.PNG

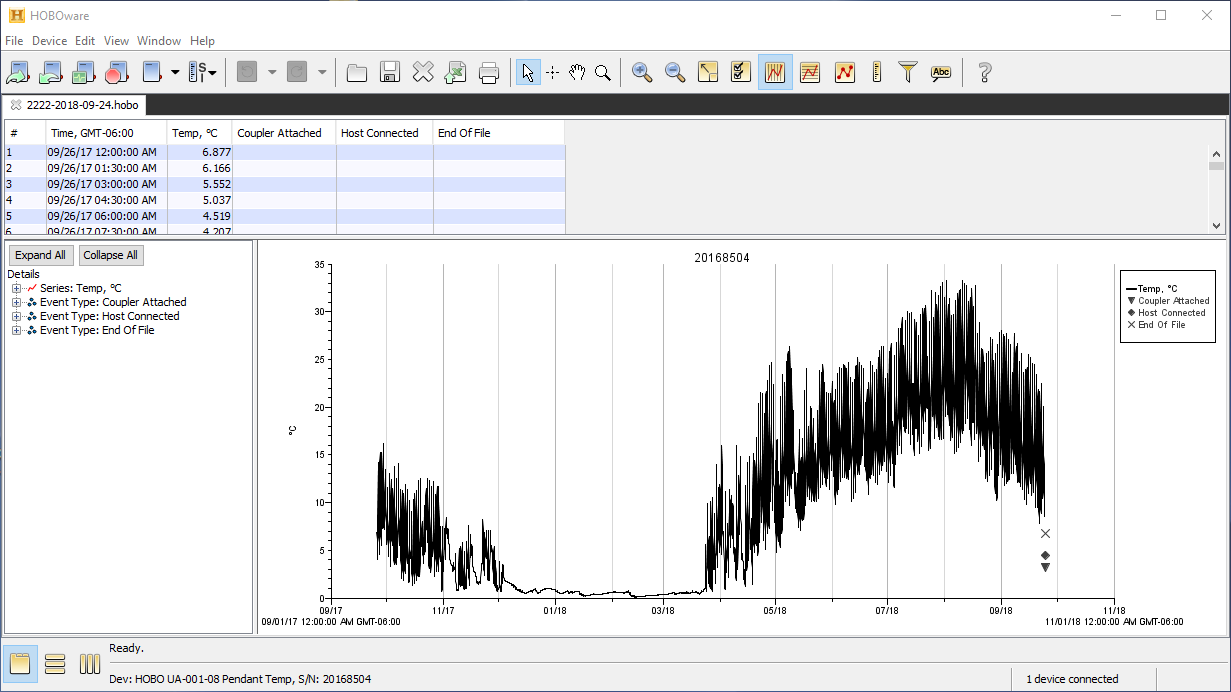

### 2222_2019-10-16_plot.PNG

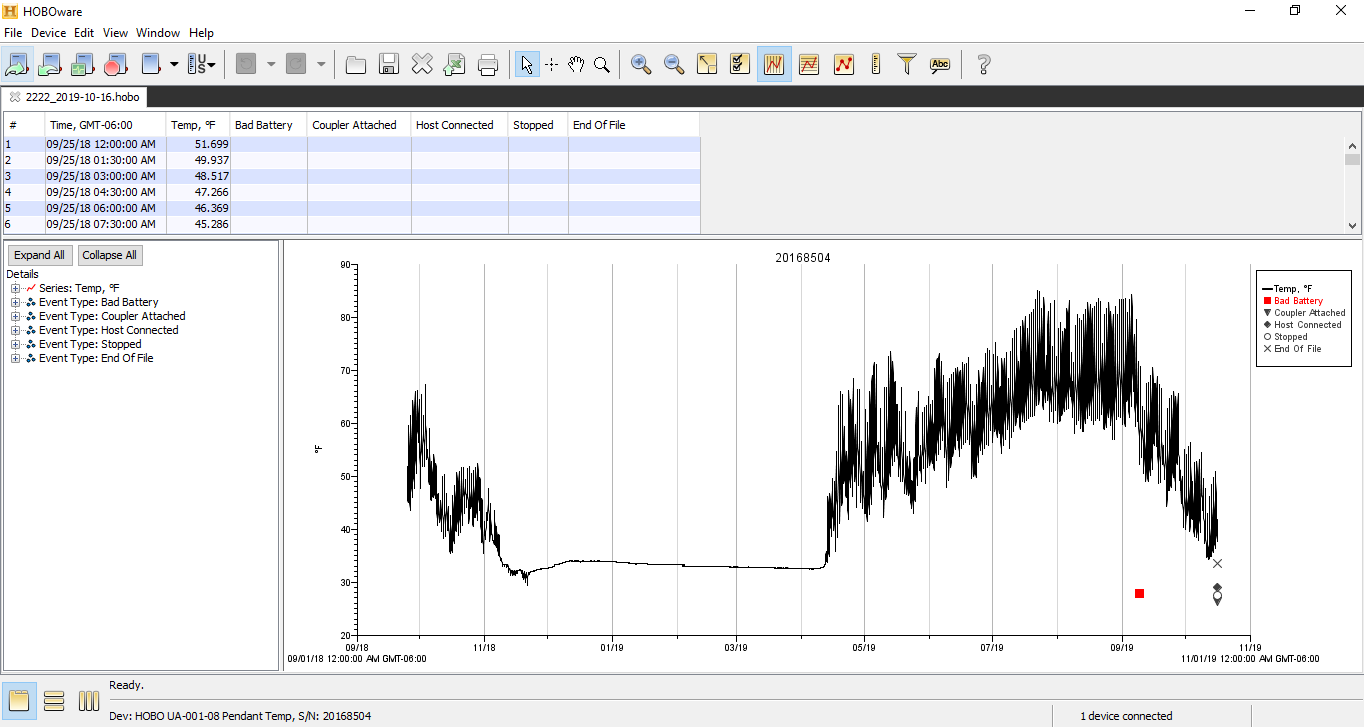

### 2222_2019-10-16_status.PNG

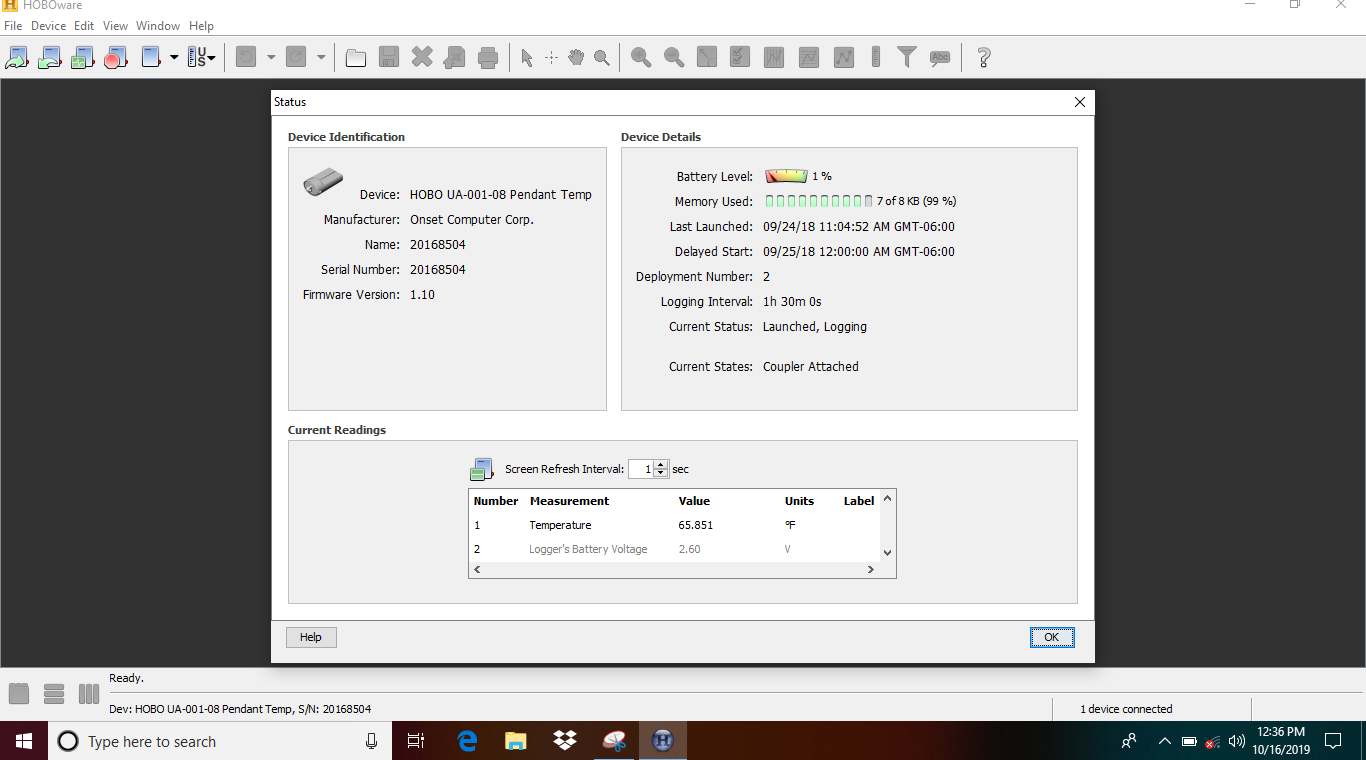

### 2233-2018-09-24.PNG

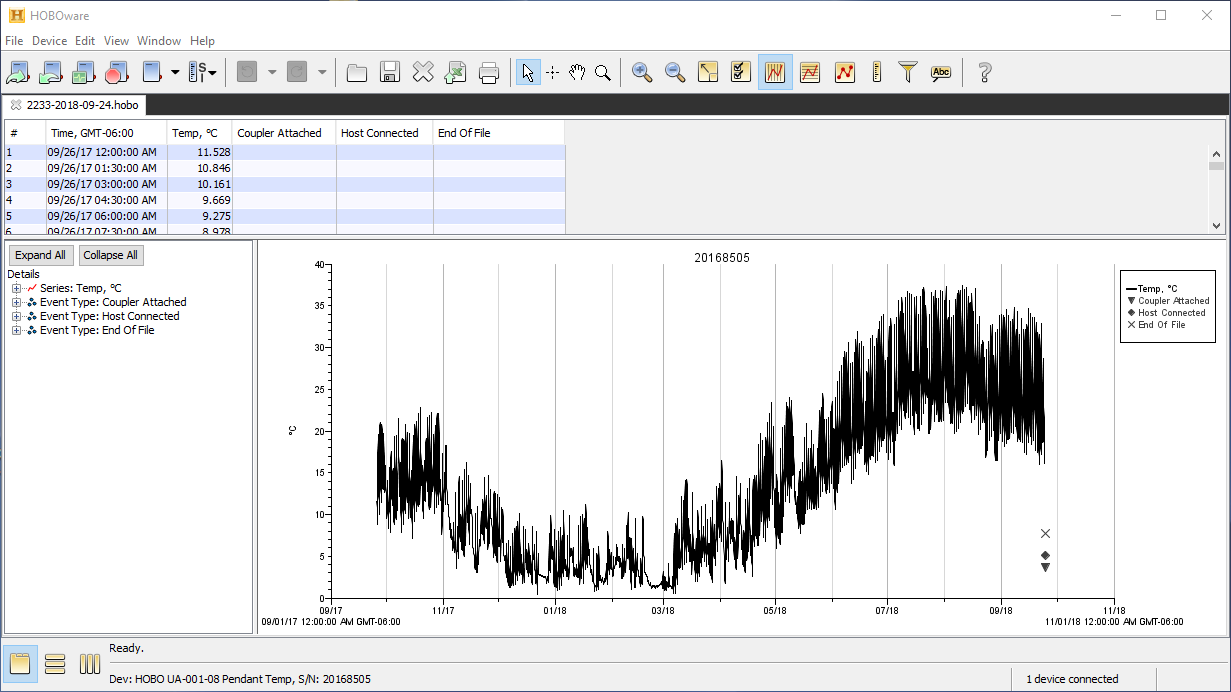

### 2233-2019-10-16_plot.PNG

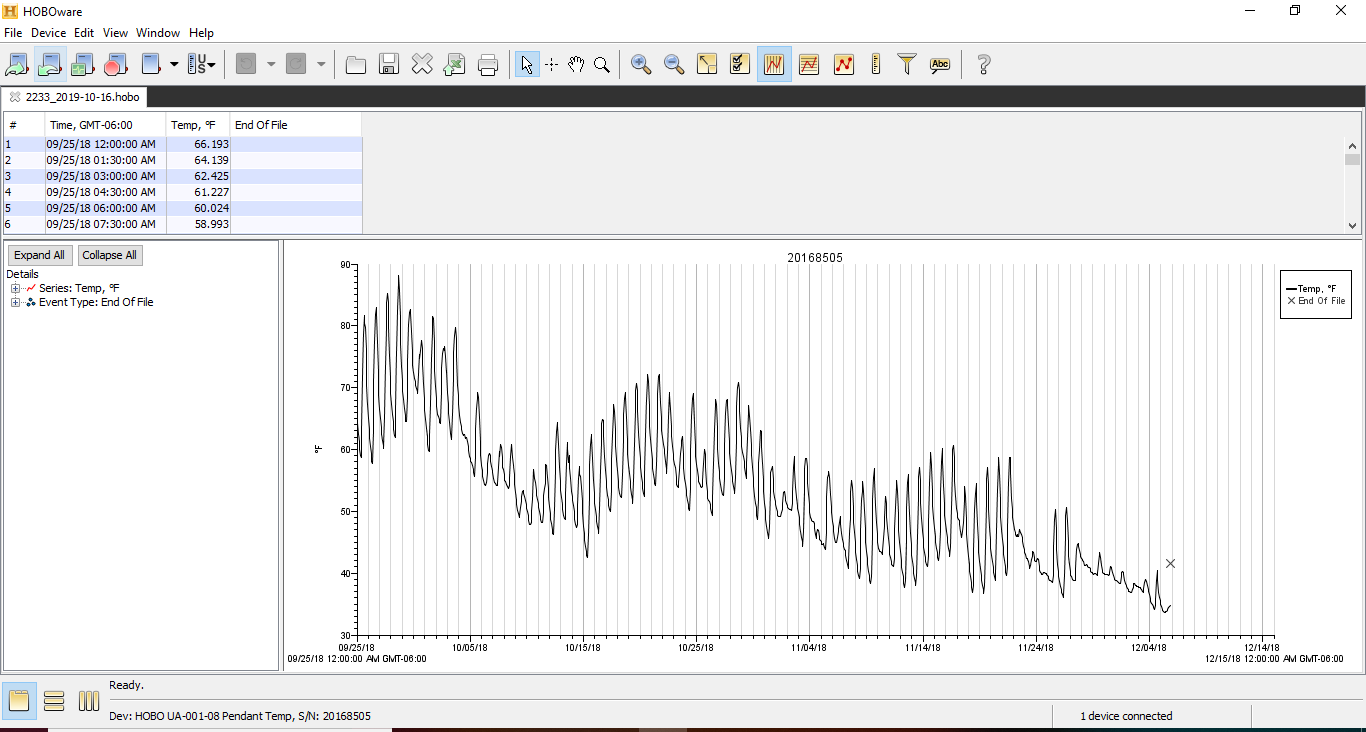

### 2233_2019-10-16_power-reset.PNG

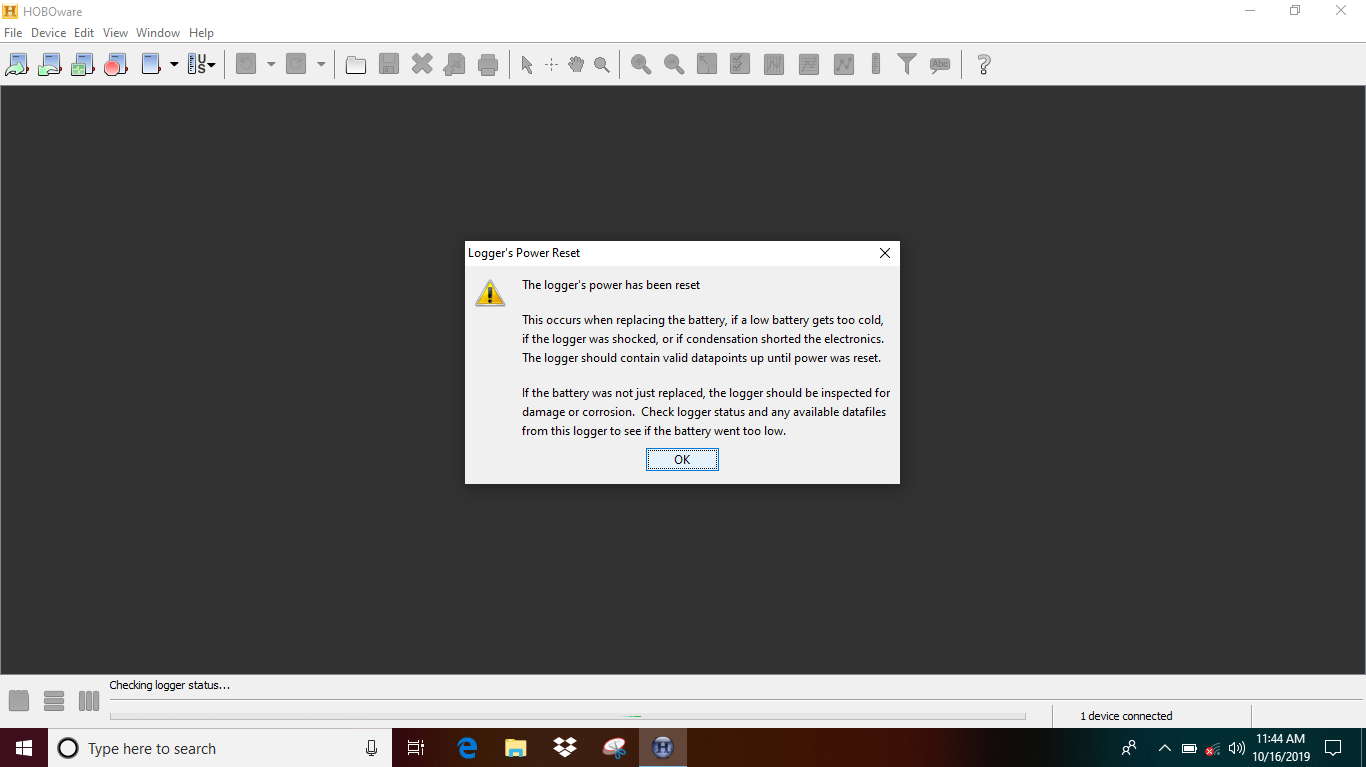

### 2233_2019-10-16_status.PNG

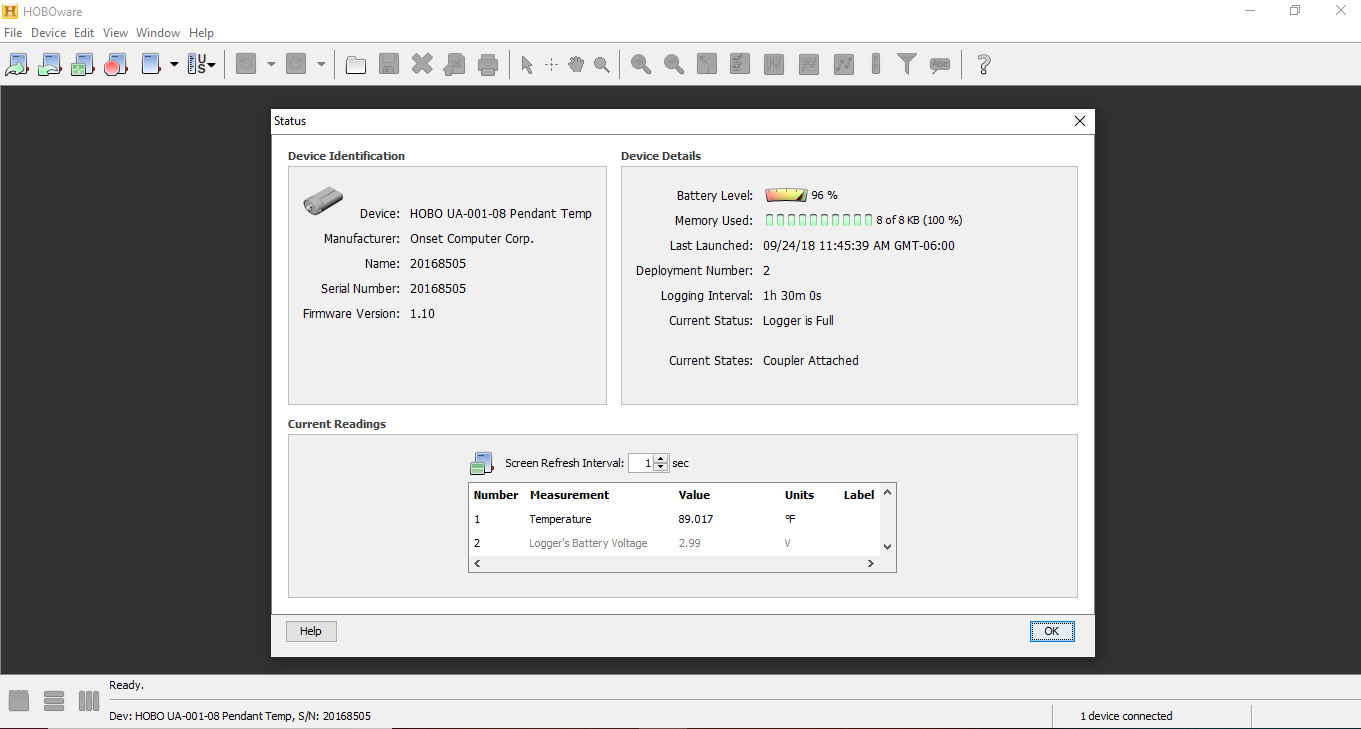

### 2311-2018-09-25.PNG

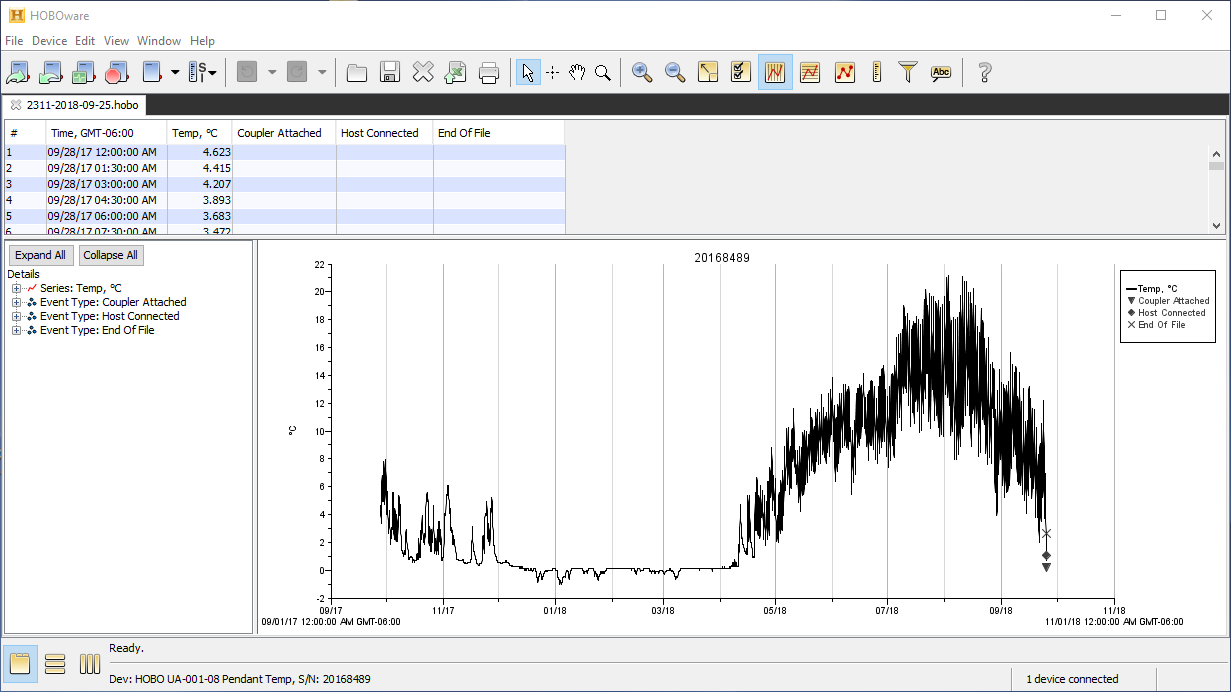

### 2311_2019-10-08_plot.PNG

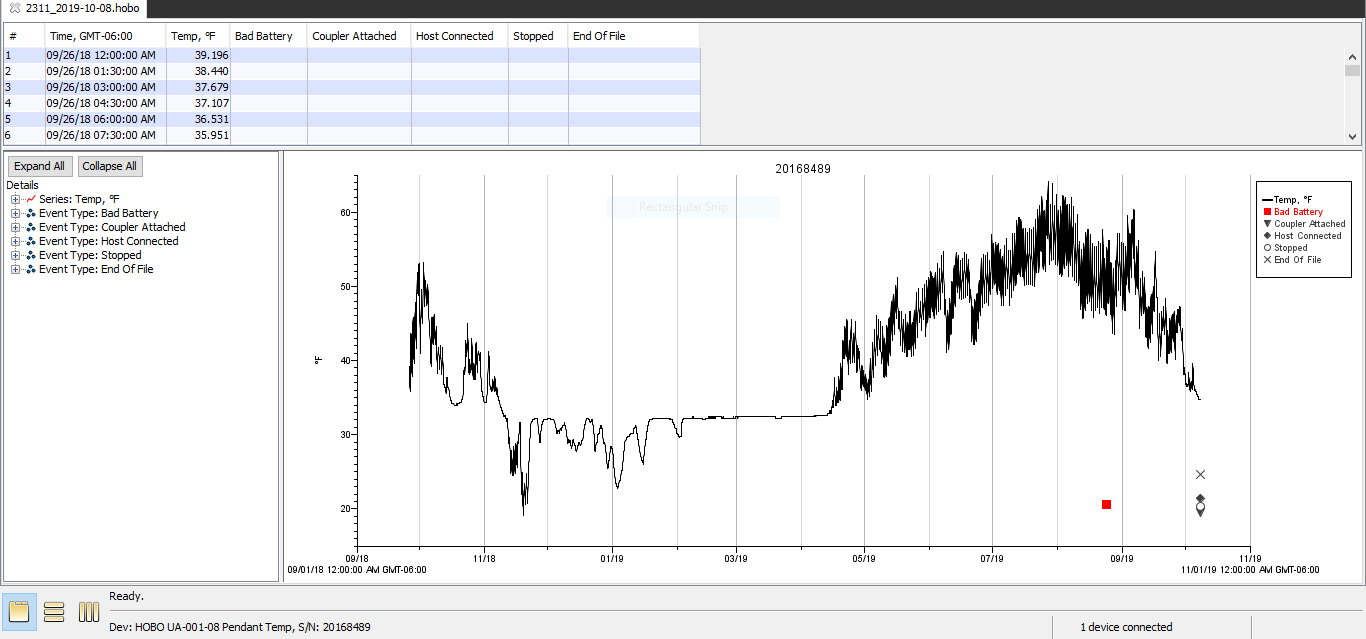

### 2311_2019-10-08_status.PNG

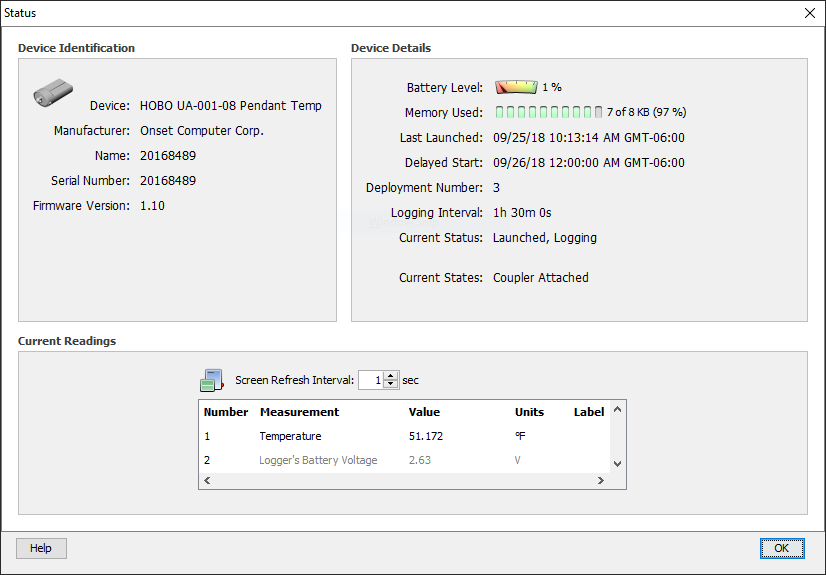

### 2333-2018-09-25.PNG

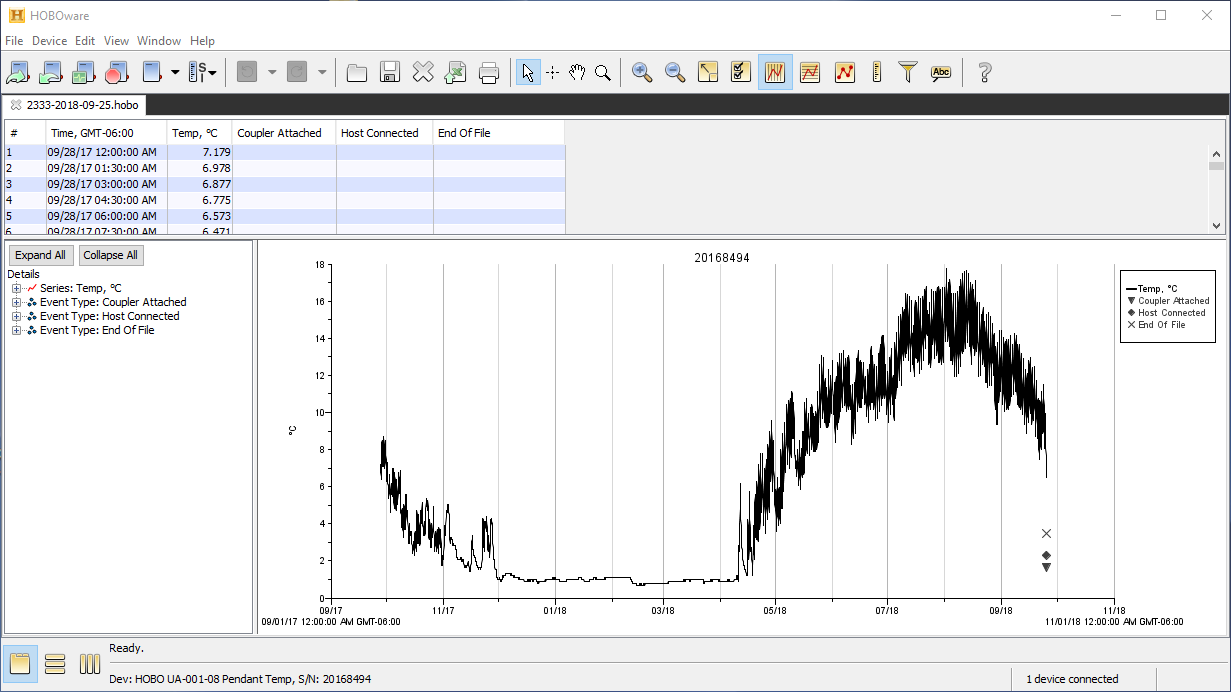

### 2333_2019-10-08_plot.PNG

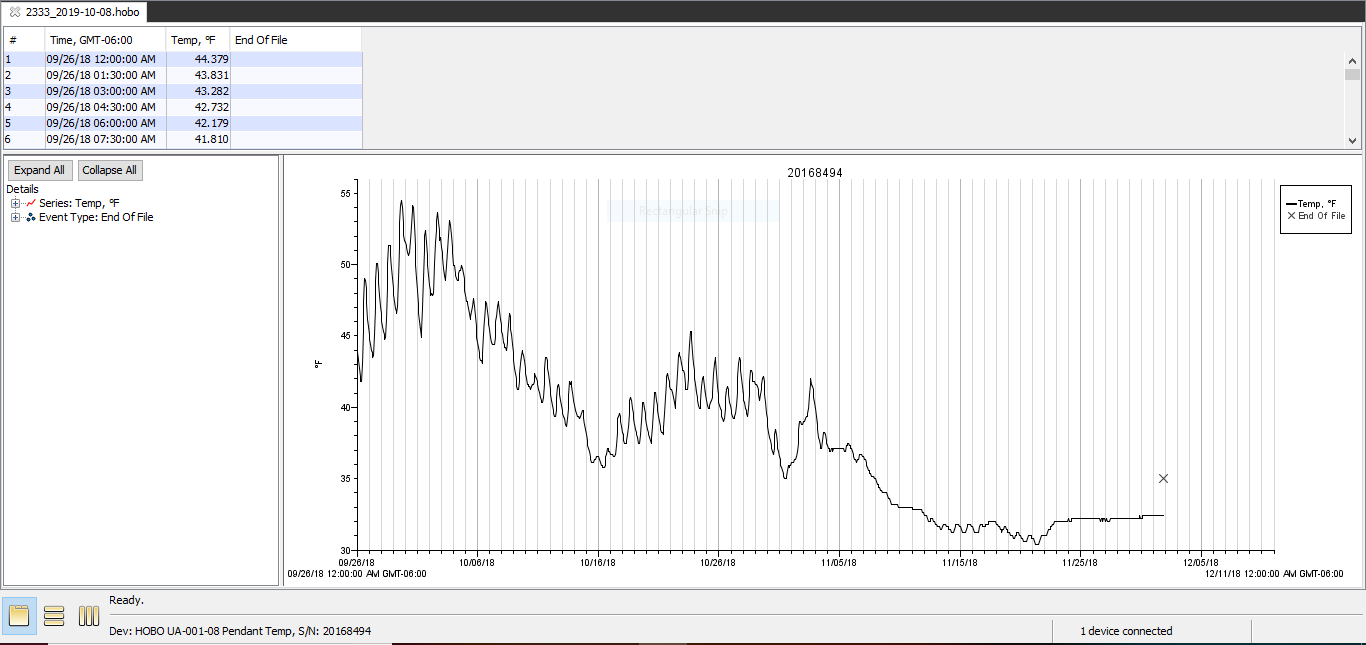

### 2333_2019-10-08_power_reset.PNG

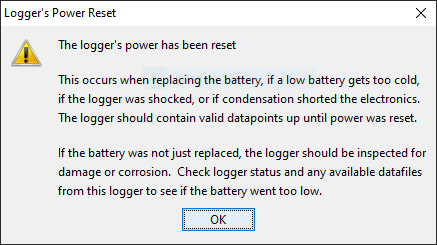

### 2333_2019-10-08_status.PNG

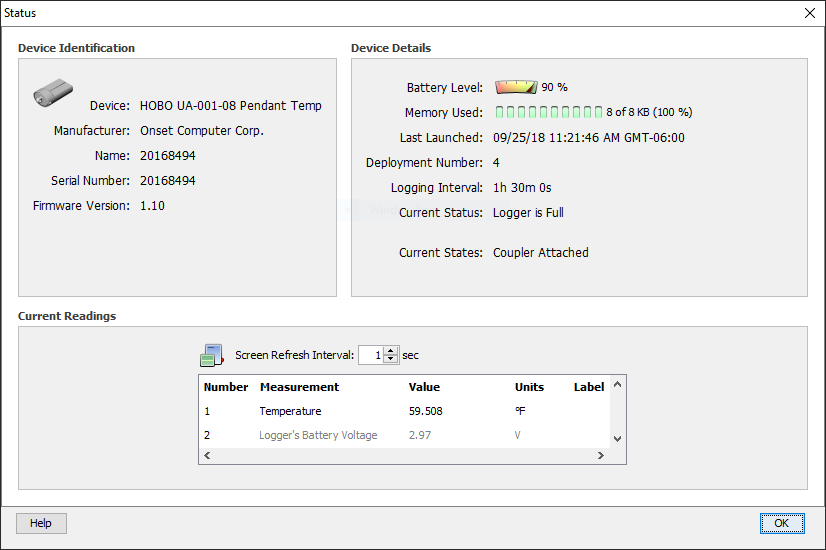

### 3111_2019-10-08_plot.PNG

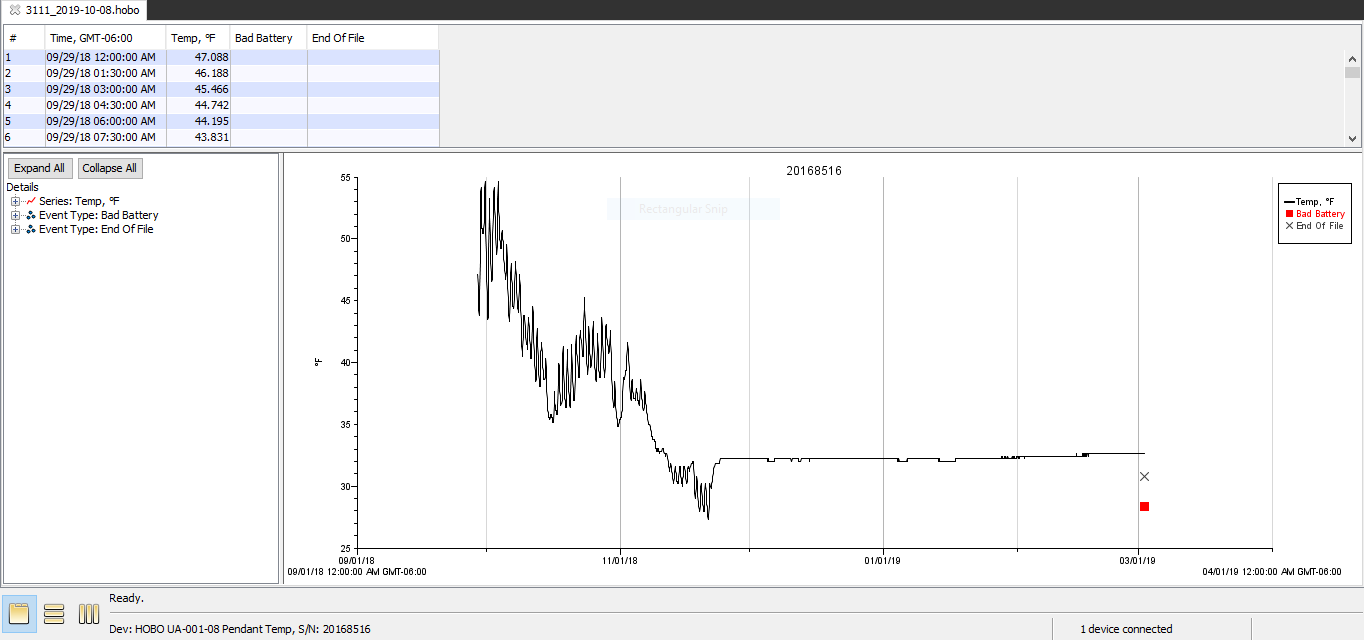

### 3111_2019-10-08_power_reset.PNG

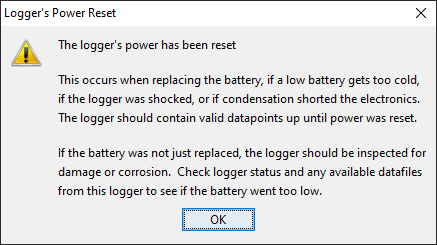

### 3111_2019-10-08_status.PNG

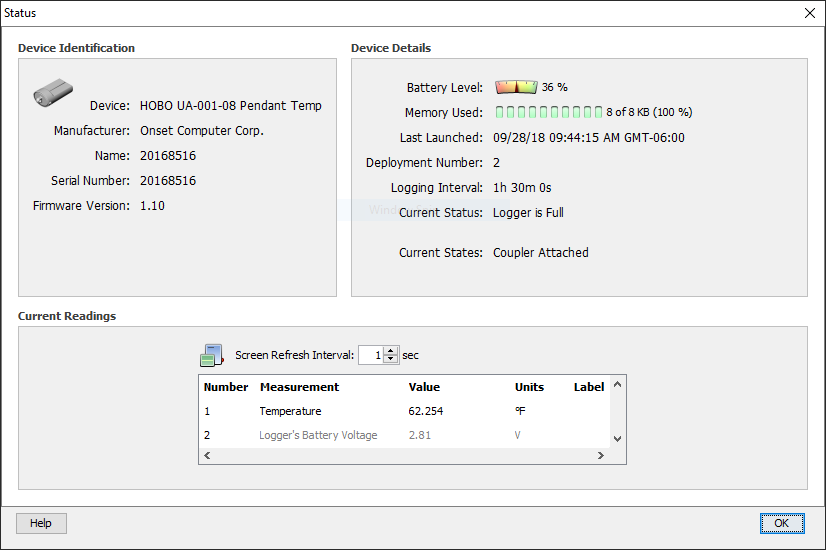

### 3122-2018-09-28.PNG

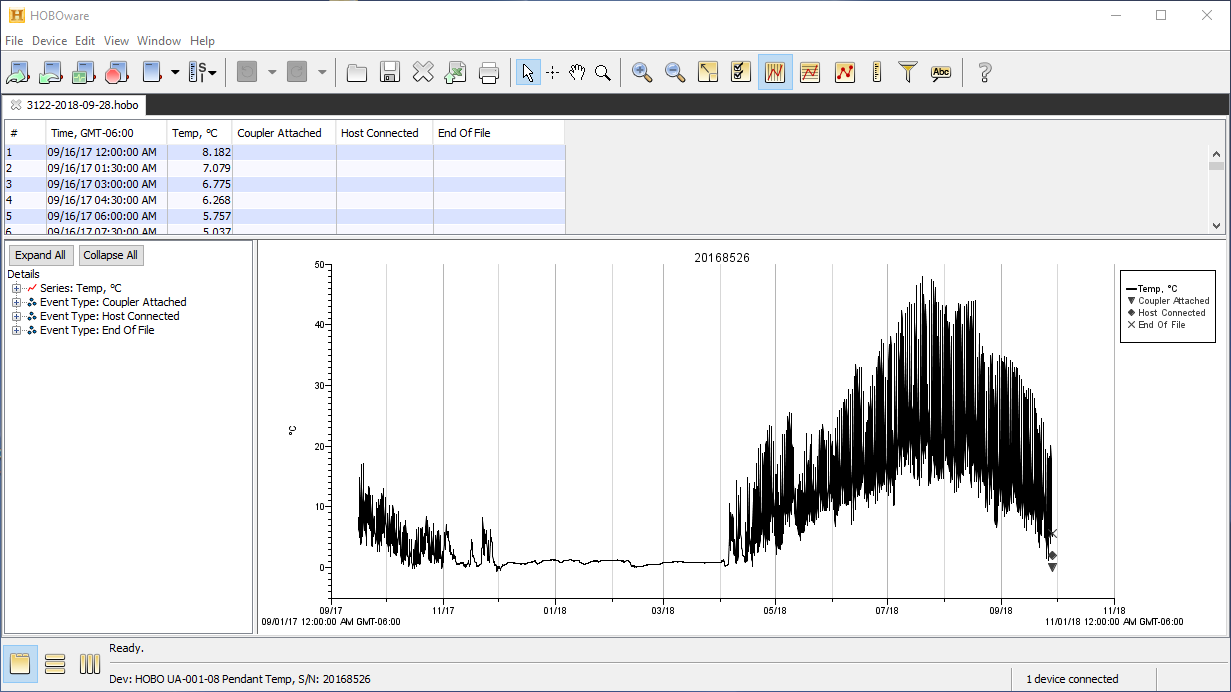

### 3122_2019-09-23_plot.PNG

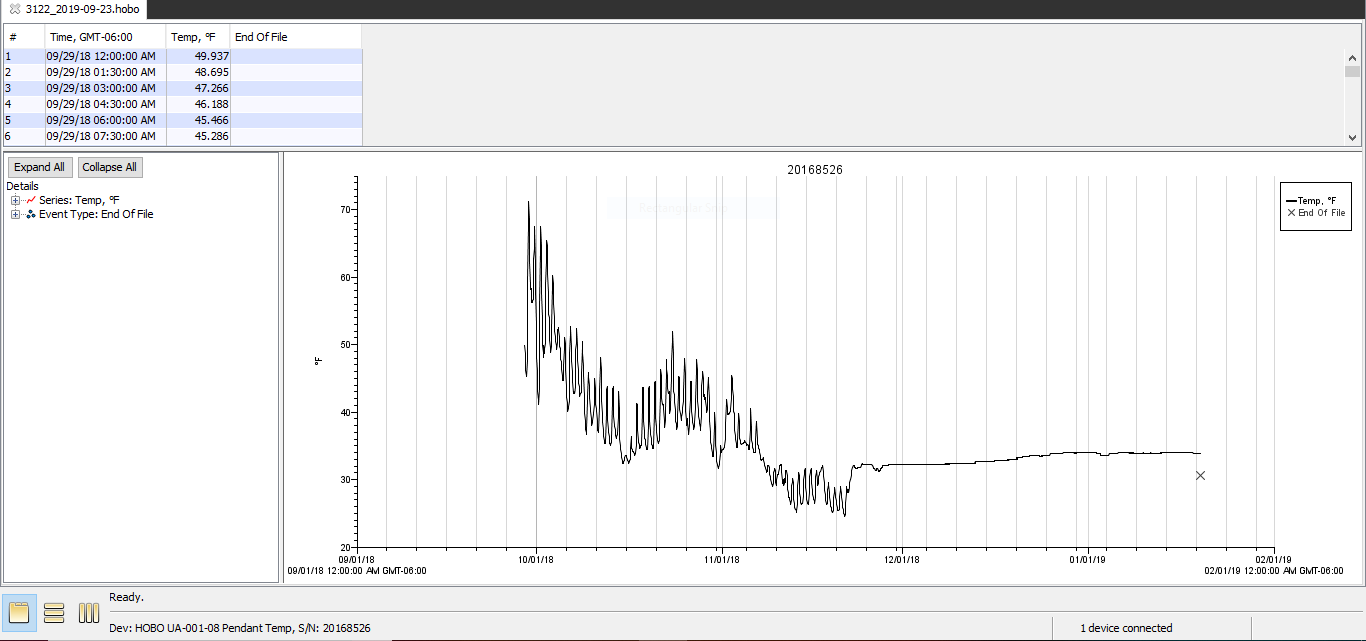

### 3122_2019-09-23_power_reset.PNG

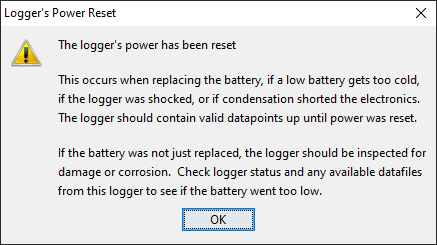

### 3122_2019-09-23_status.PNG

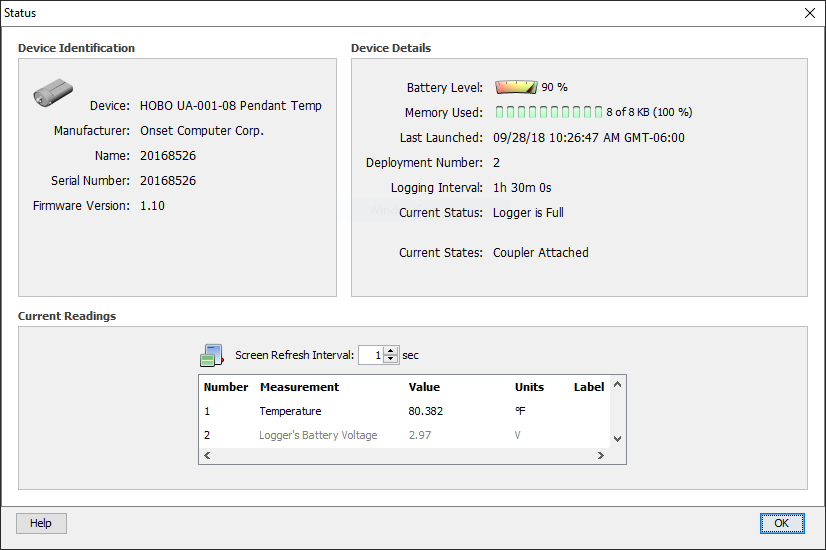

### 3133 HOBO Power Reset.PNG

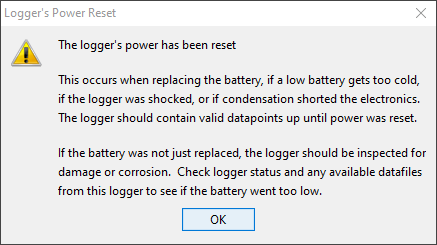

### 3133-2018-09-28.PNG

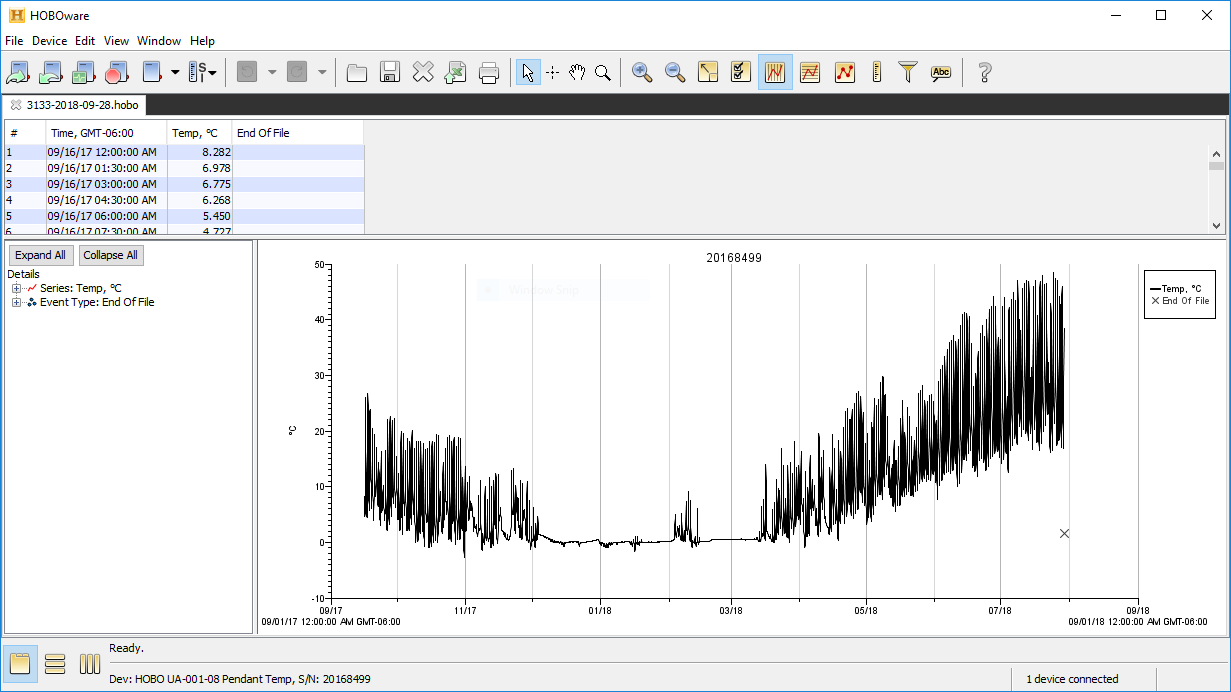

### 3133_2019-09-23_plot.PNG

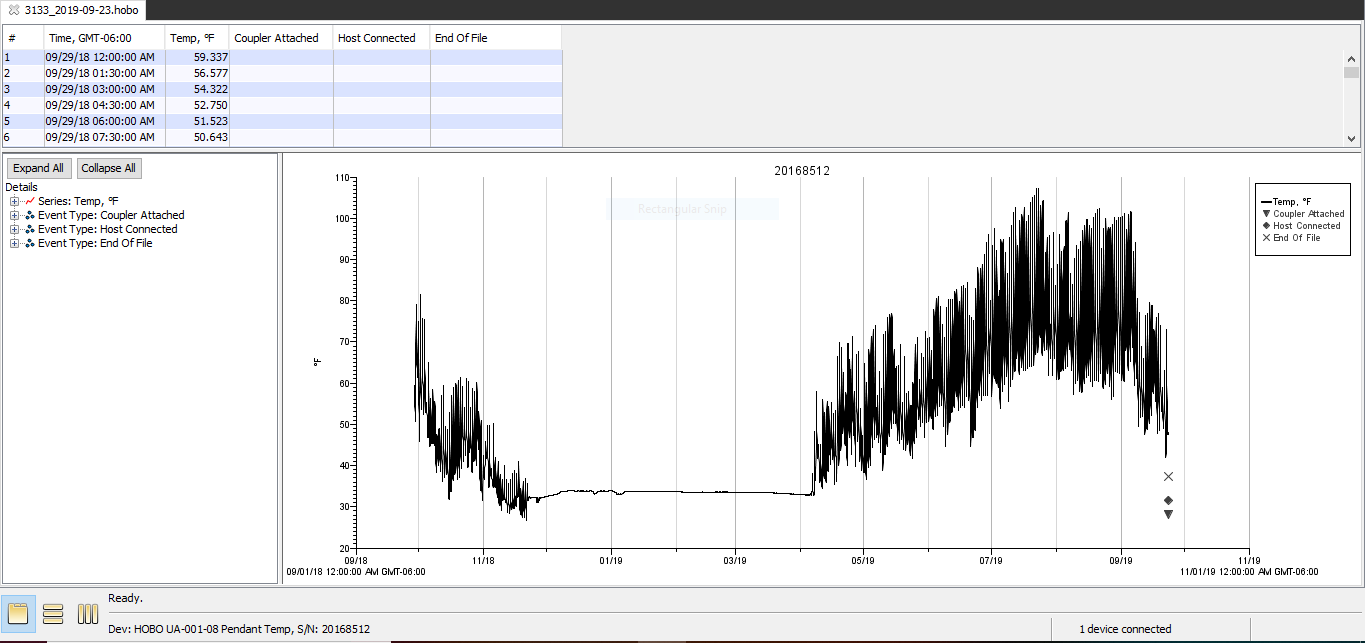

### 3133_2019-09-23_status.PNG

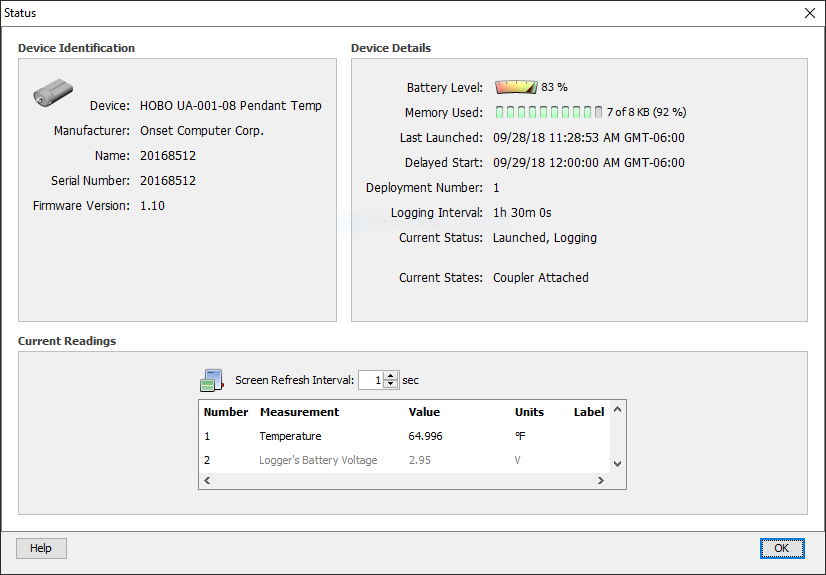

### 3211-2018-10-01.PNG

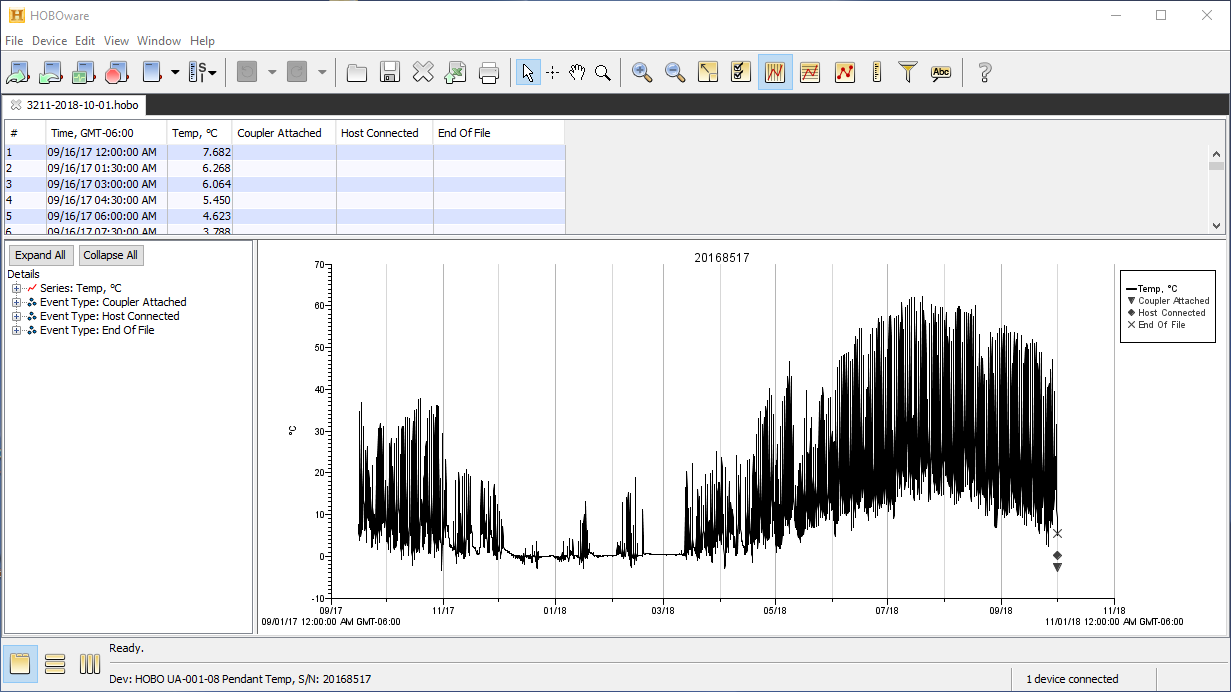

### 3211_2019-09-23_plot.PNG

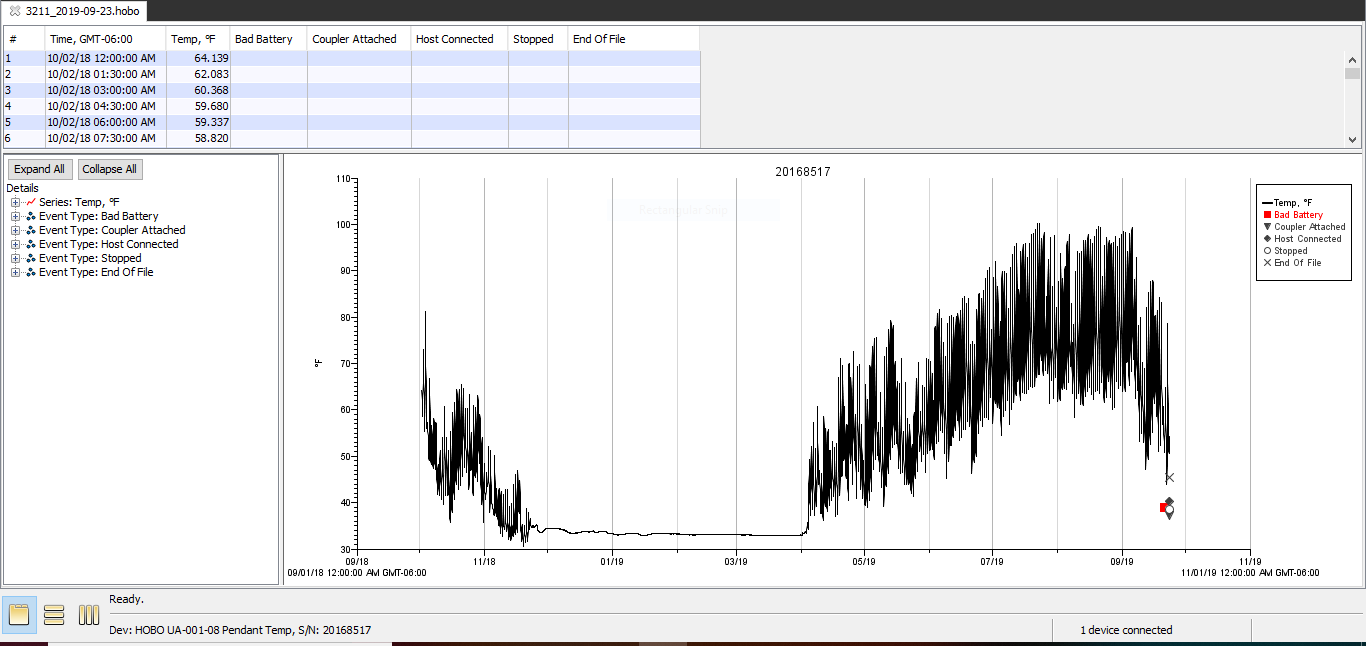

### 3211_2019-09-23_status.PNG

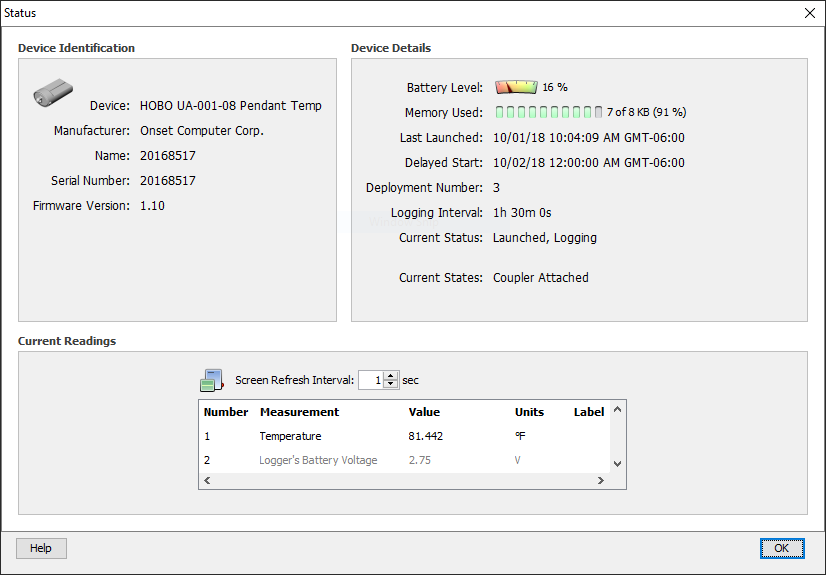

### 3222-2018-10-01.PNG

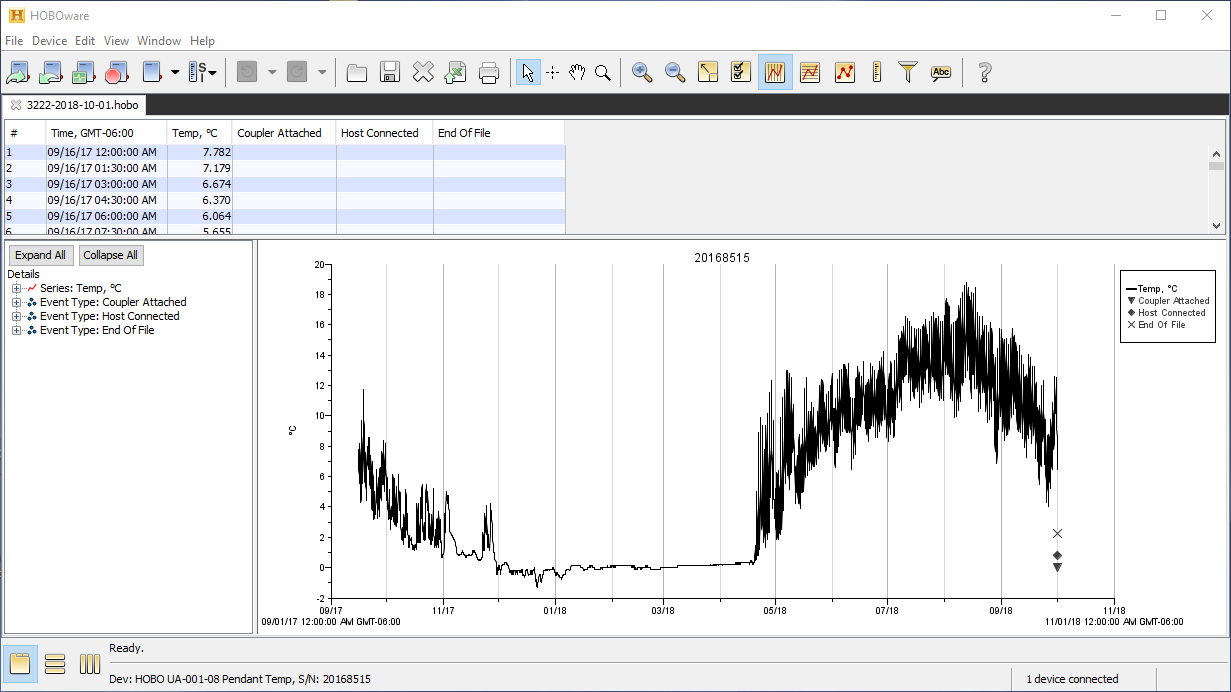

### 3222_2019-09-19_plot.PNG

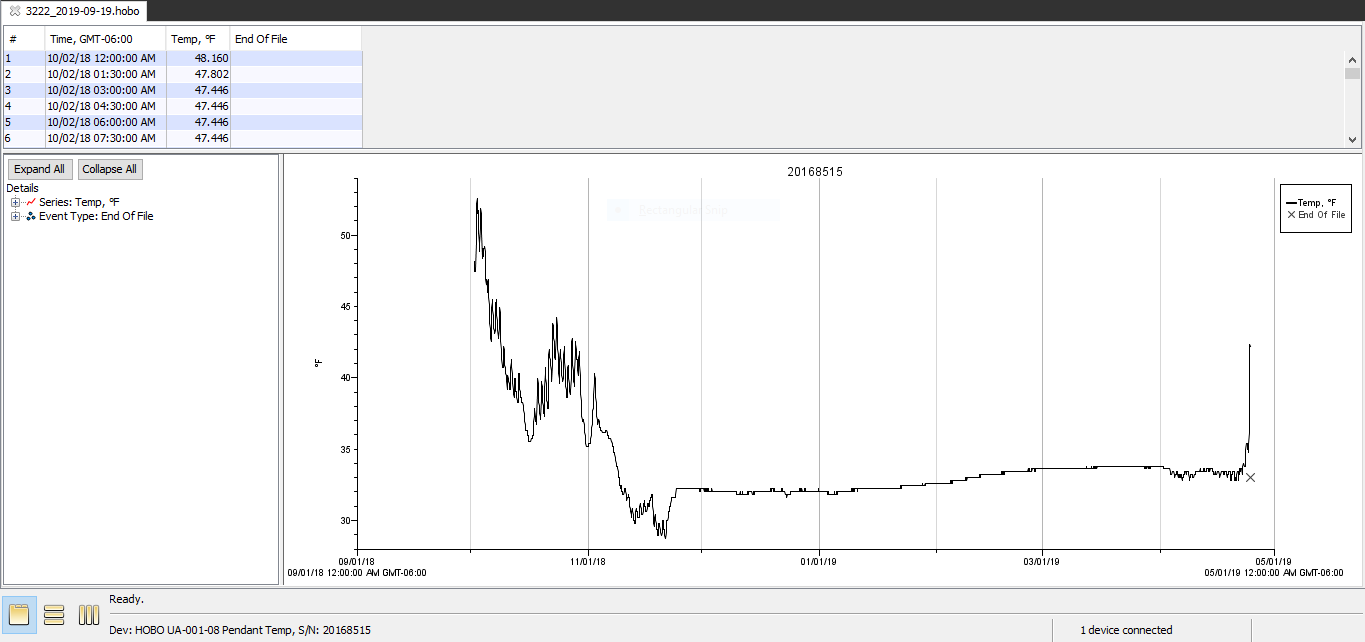

### 3222_2019-09-19_power_reset.PNG

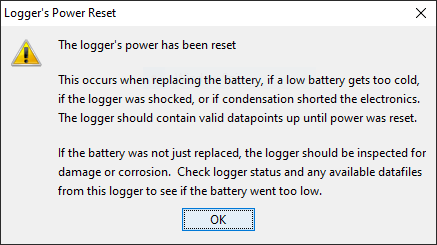

### 3222_2019-09-19_status.PNG

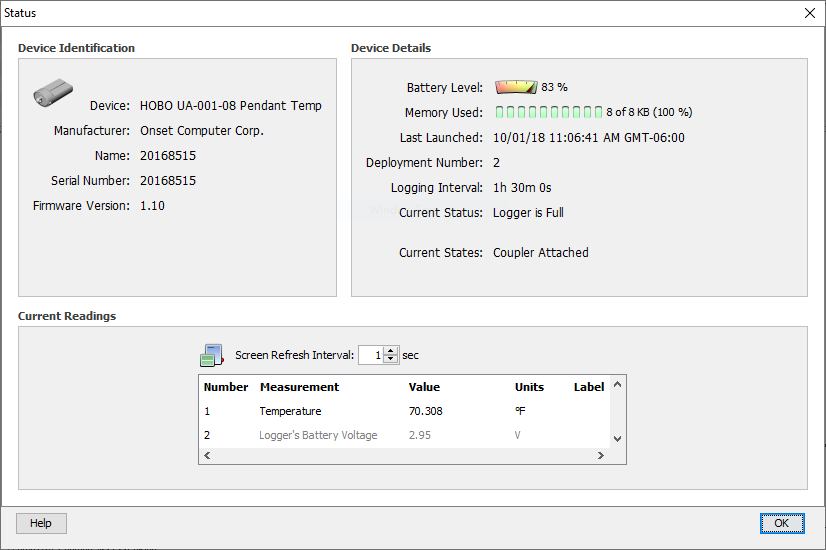

### 3233-2018-10-01.PNG

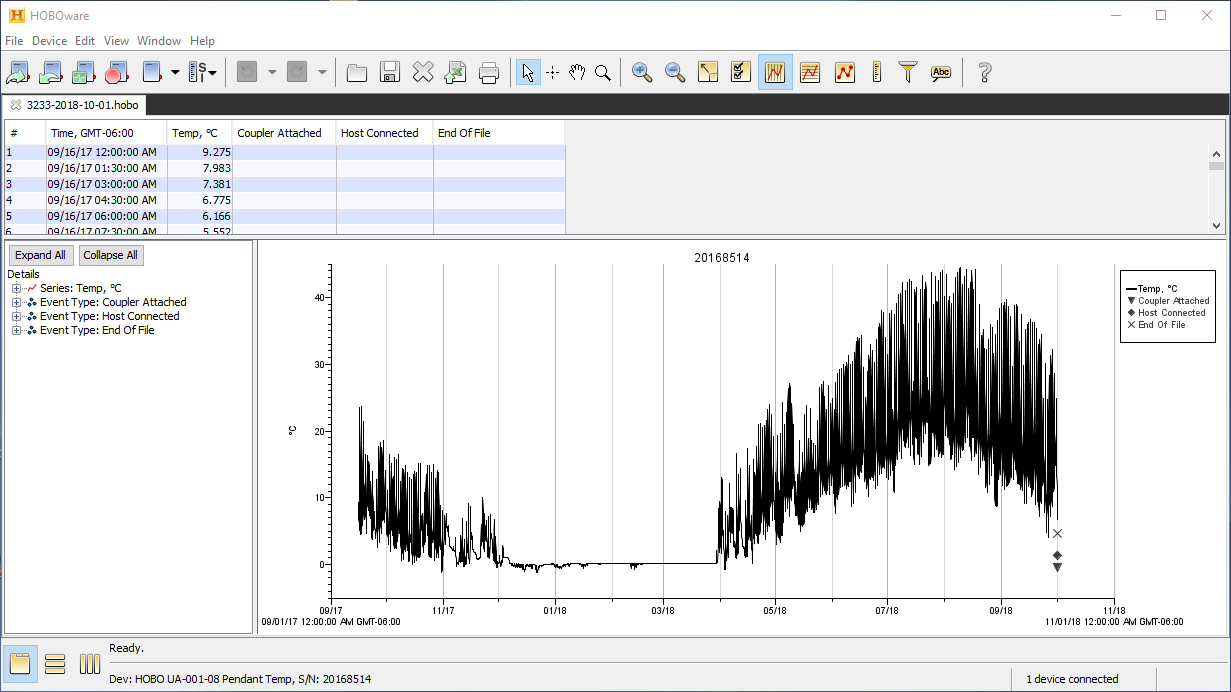

### 3233_2019-09-19_plot.PNG

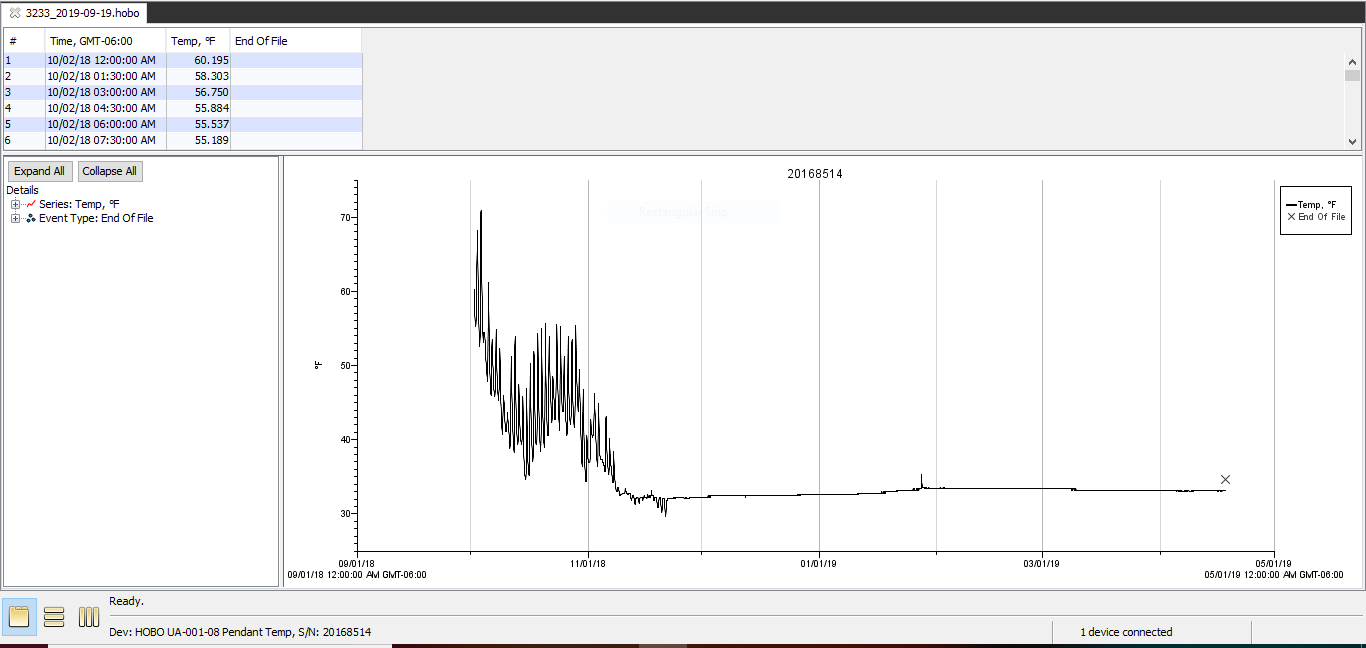

### 3233_2019-09-19_power_reset.PNG

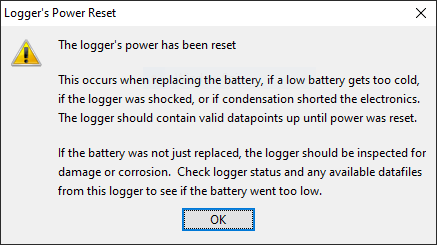

### 3233_2019-09-19_status.PNG

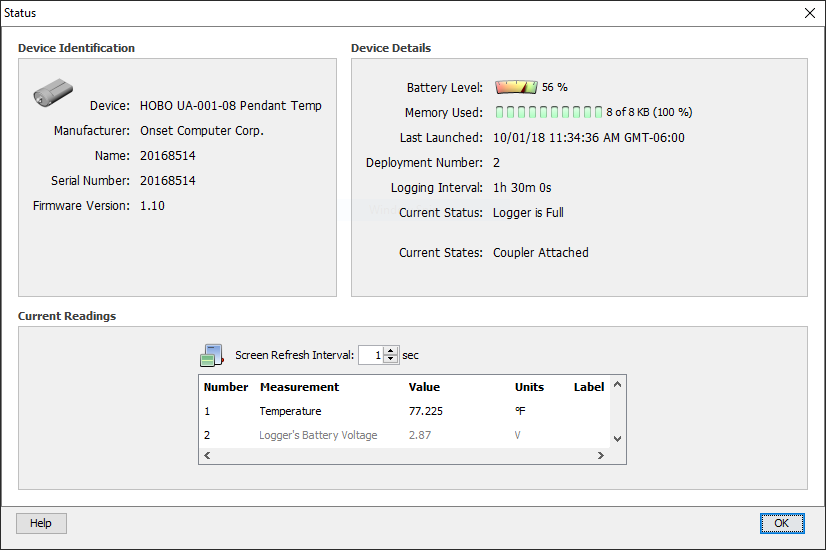

### 3311-2018-10-02.PNG

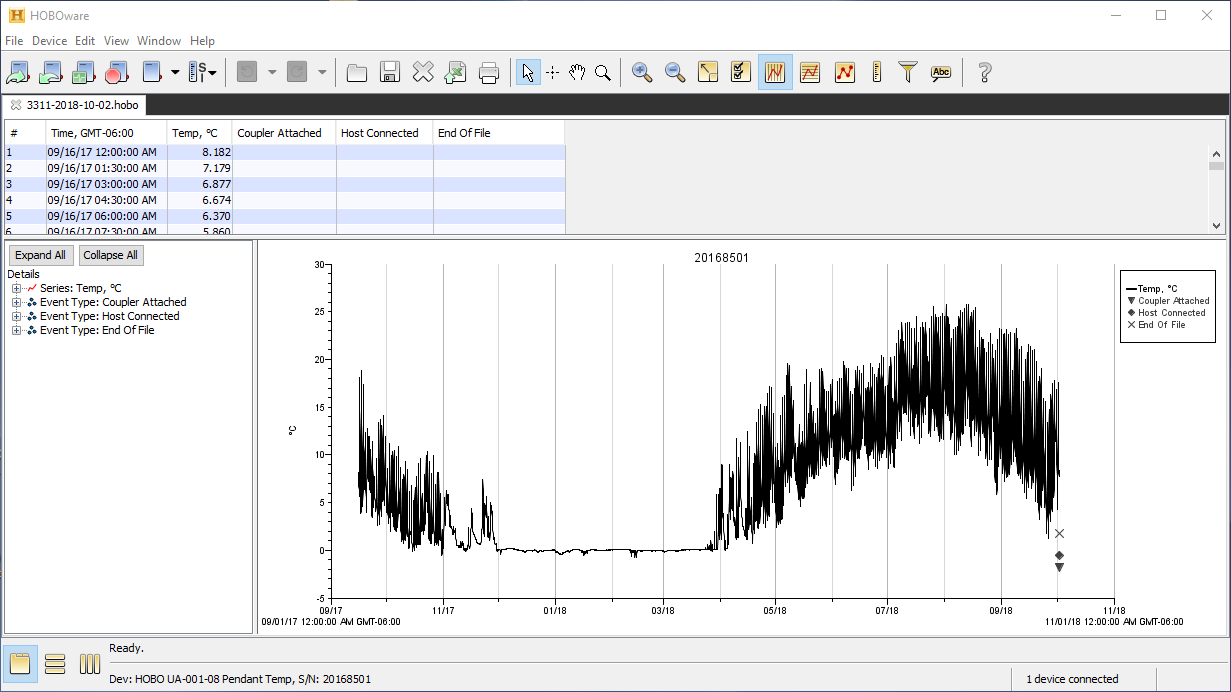

### 3311_2019-09-19_plot.PNG

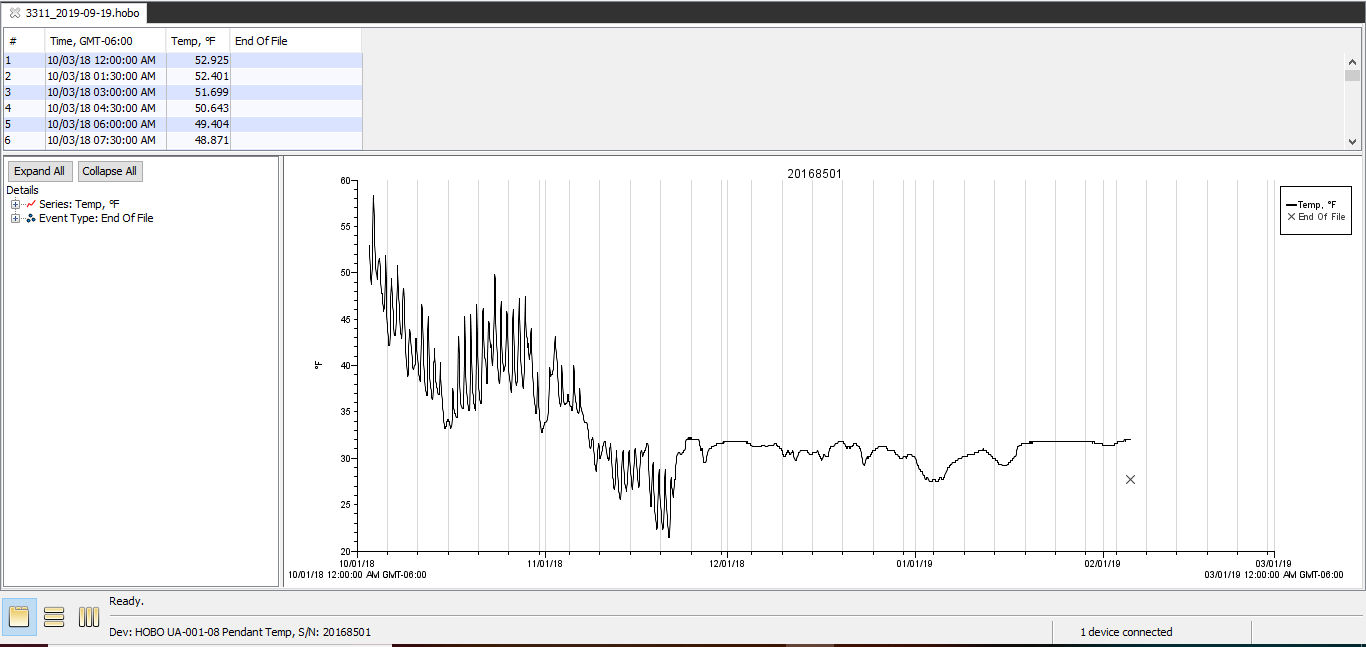

### 3311_2019-09-19_power_reset.PNG

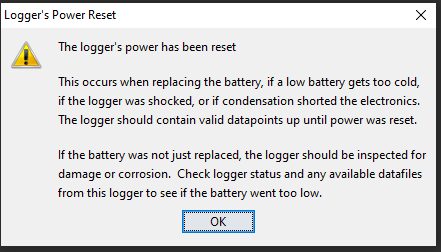

### 3311_2019-09-19_status.PNG

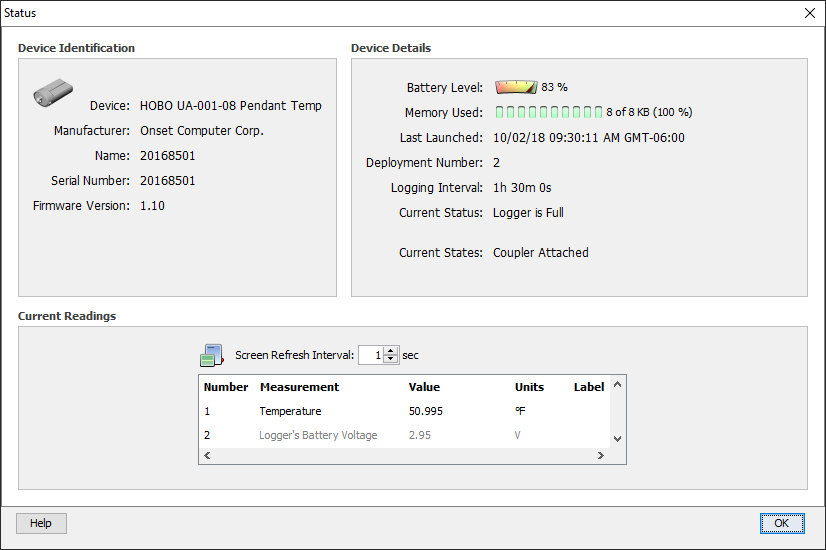

### 3322-2018-10-02.PNG

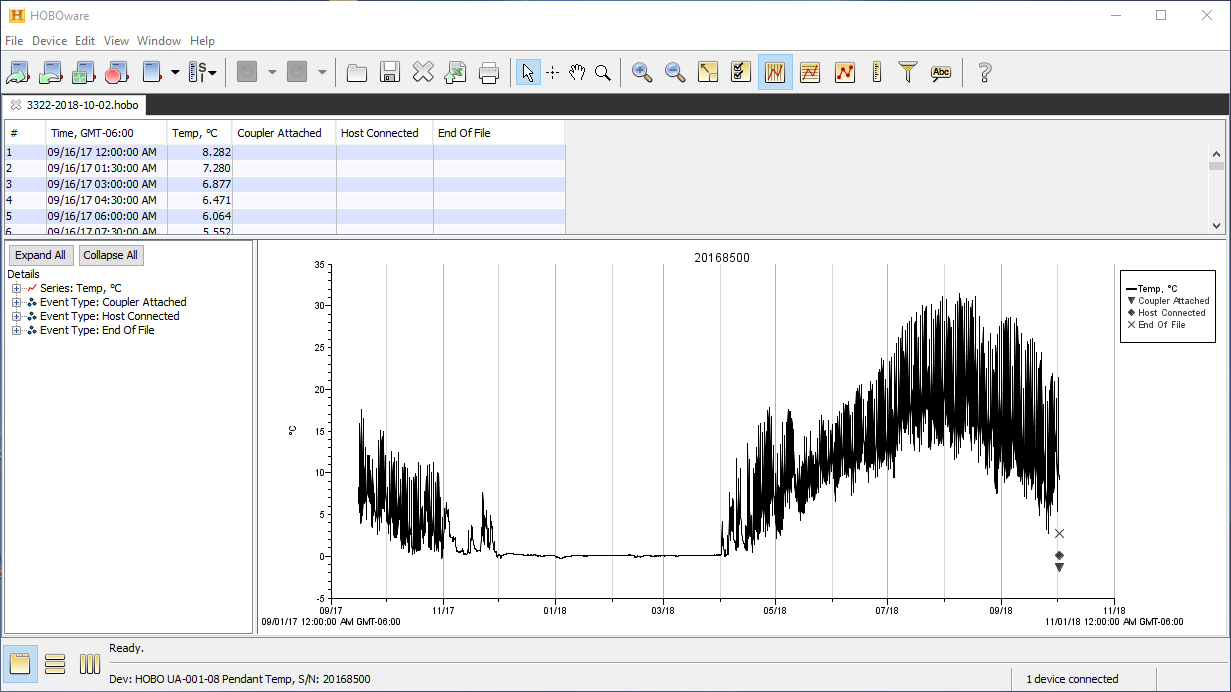

### 3333-2018-10-02.PNG

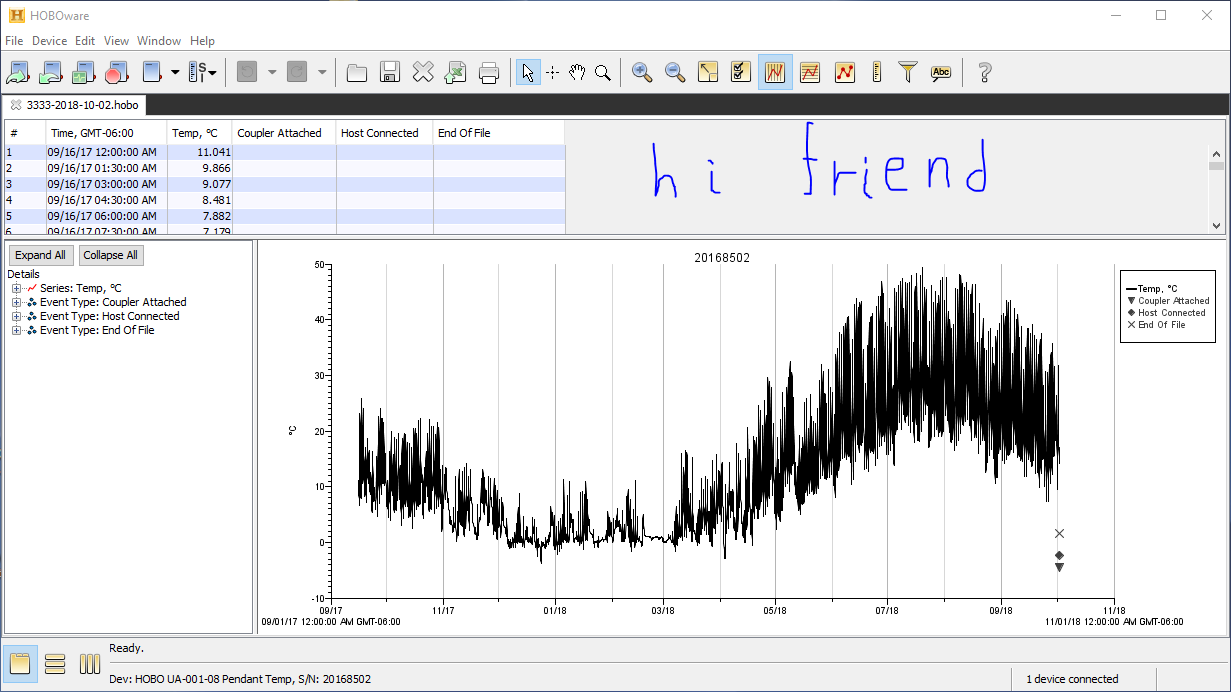

### 3333_2019-09-19_plot.PNG

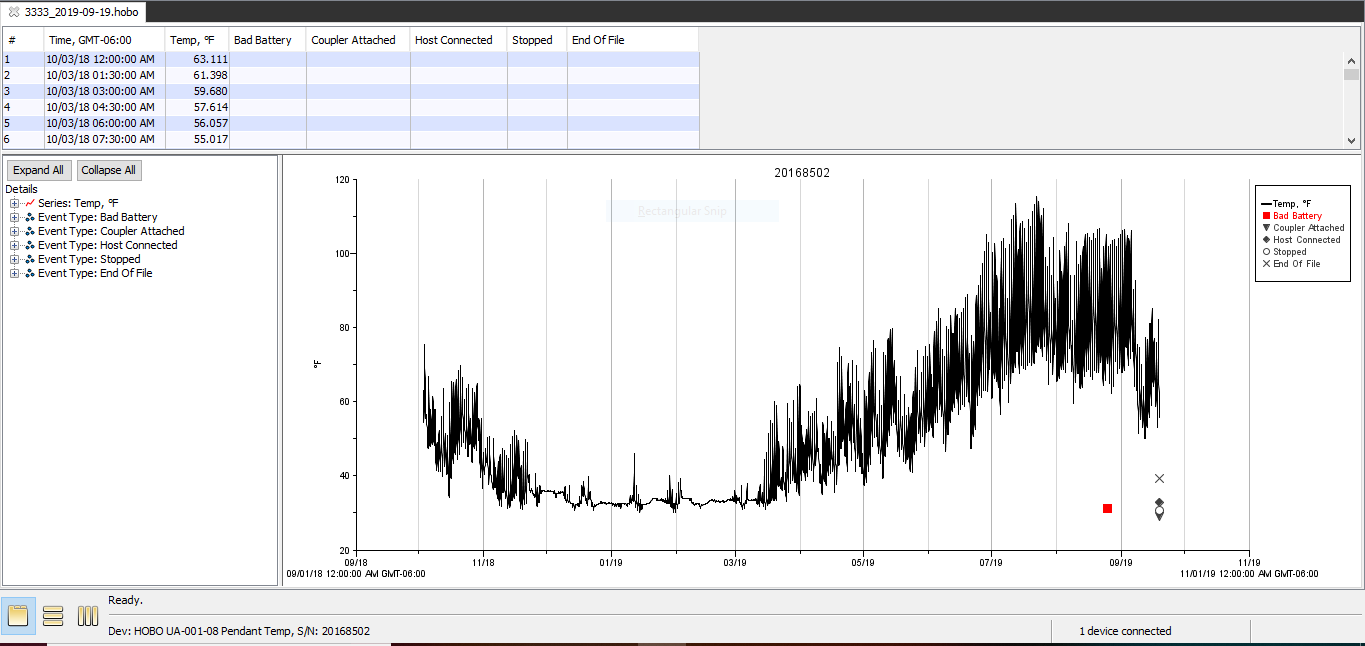

### 3333_2019-09-19_status.PNG

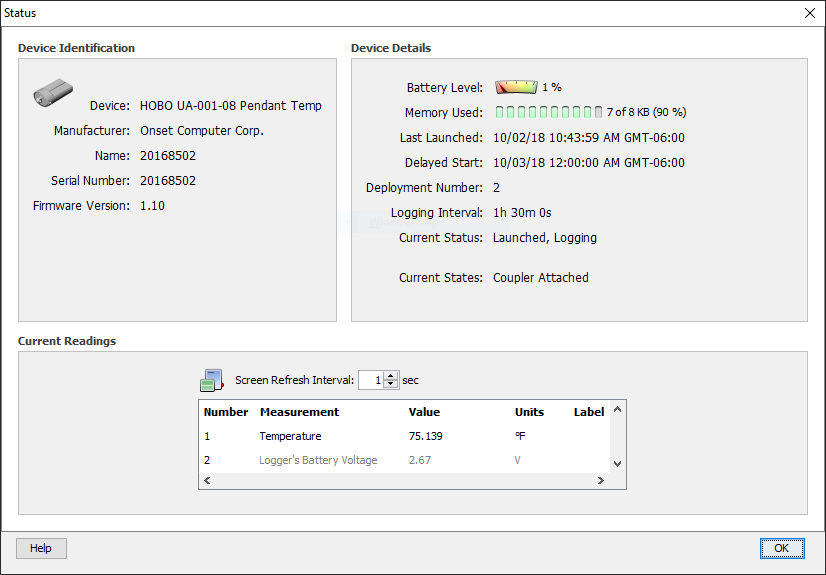

### env-core18.jpeg

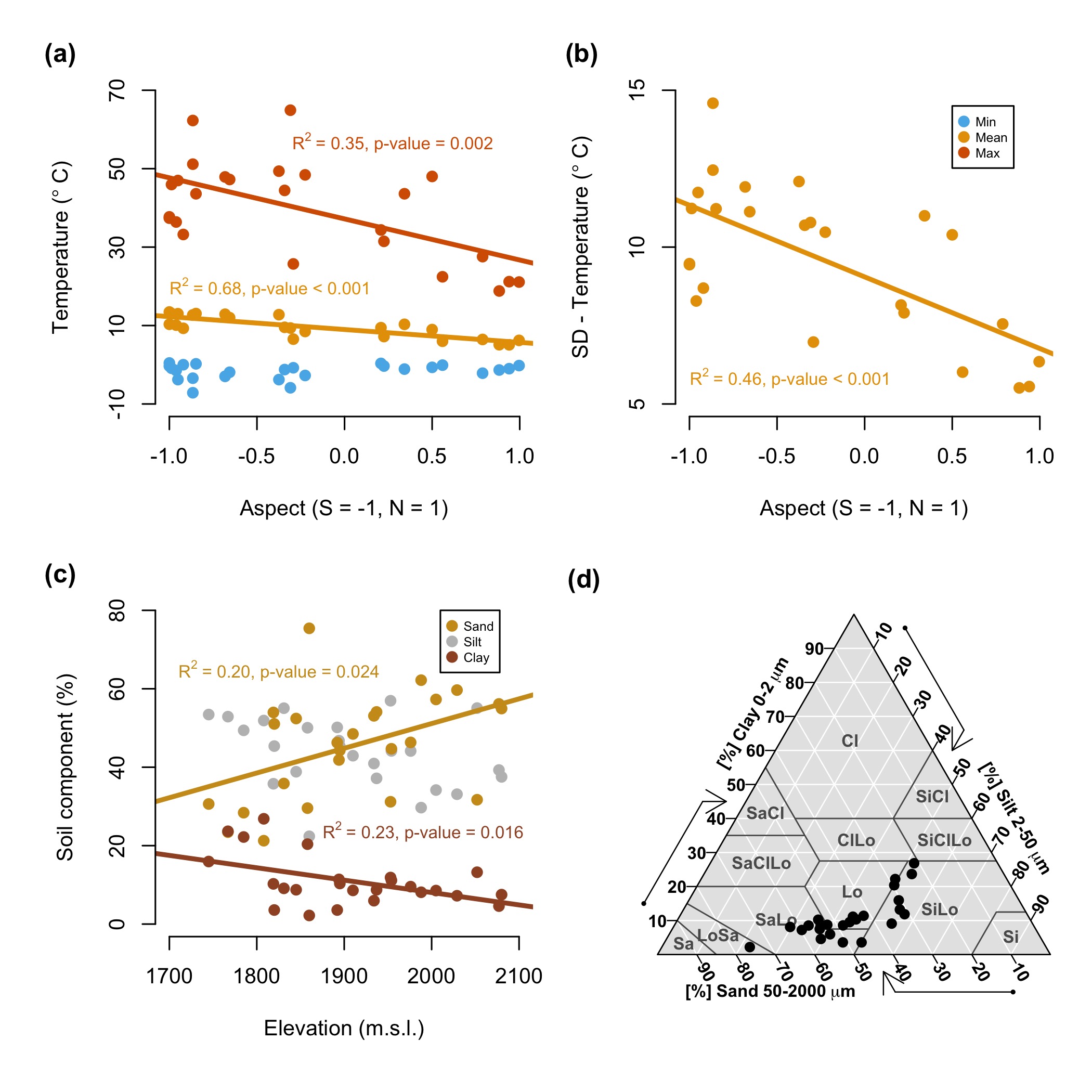

### fdiv-env-core18-wLH.jpeg

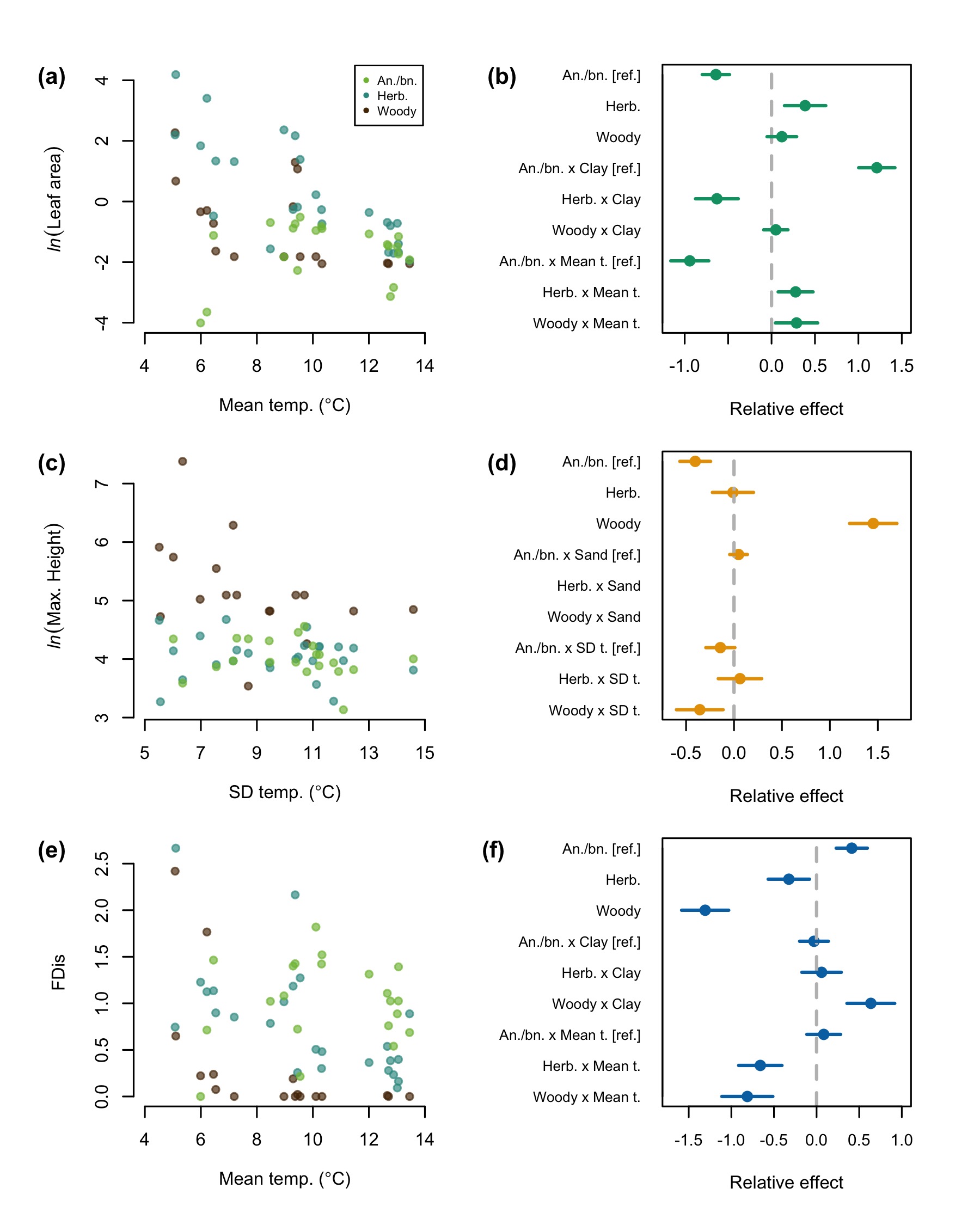

### fdiv-env-core18.jpeg

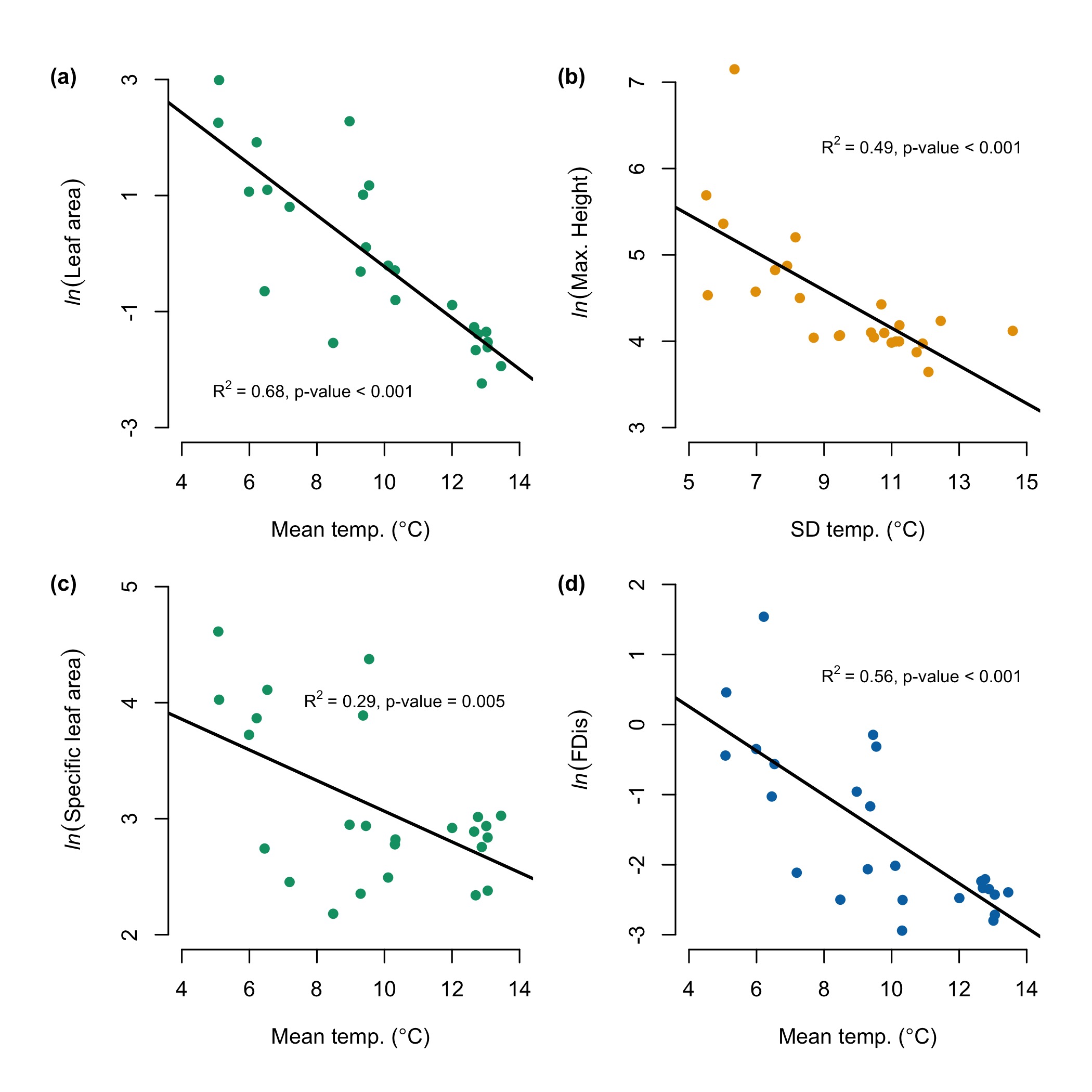

### FT-fig1.jpeg

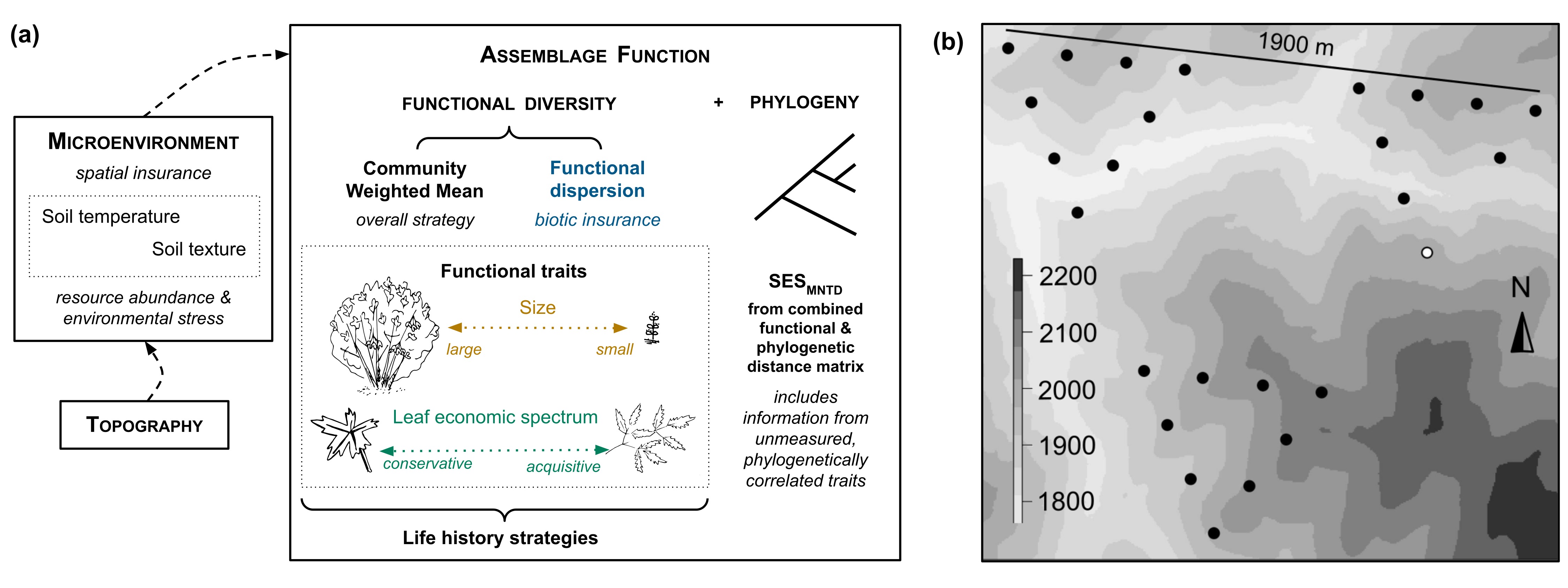

### funct-phy-core18.jpeg

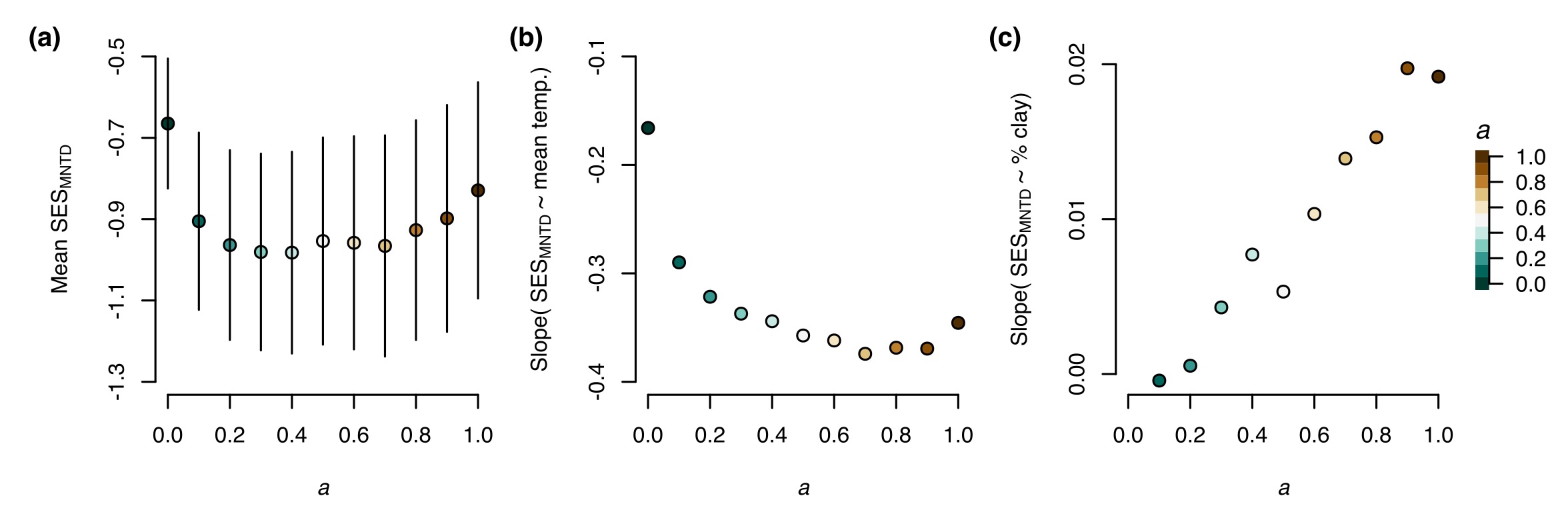

### SUPP-fdiv-env-core18-wLH.jpeg

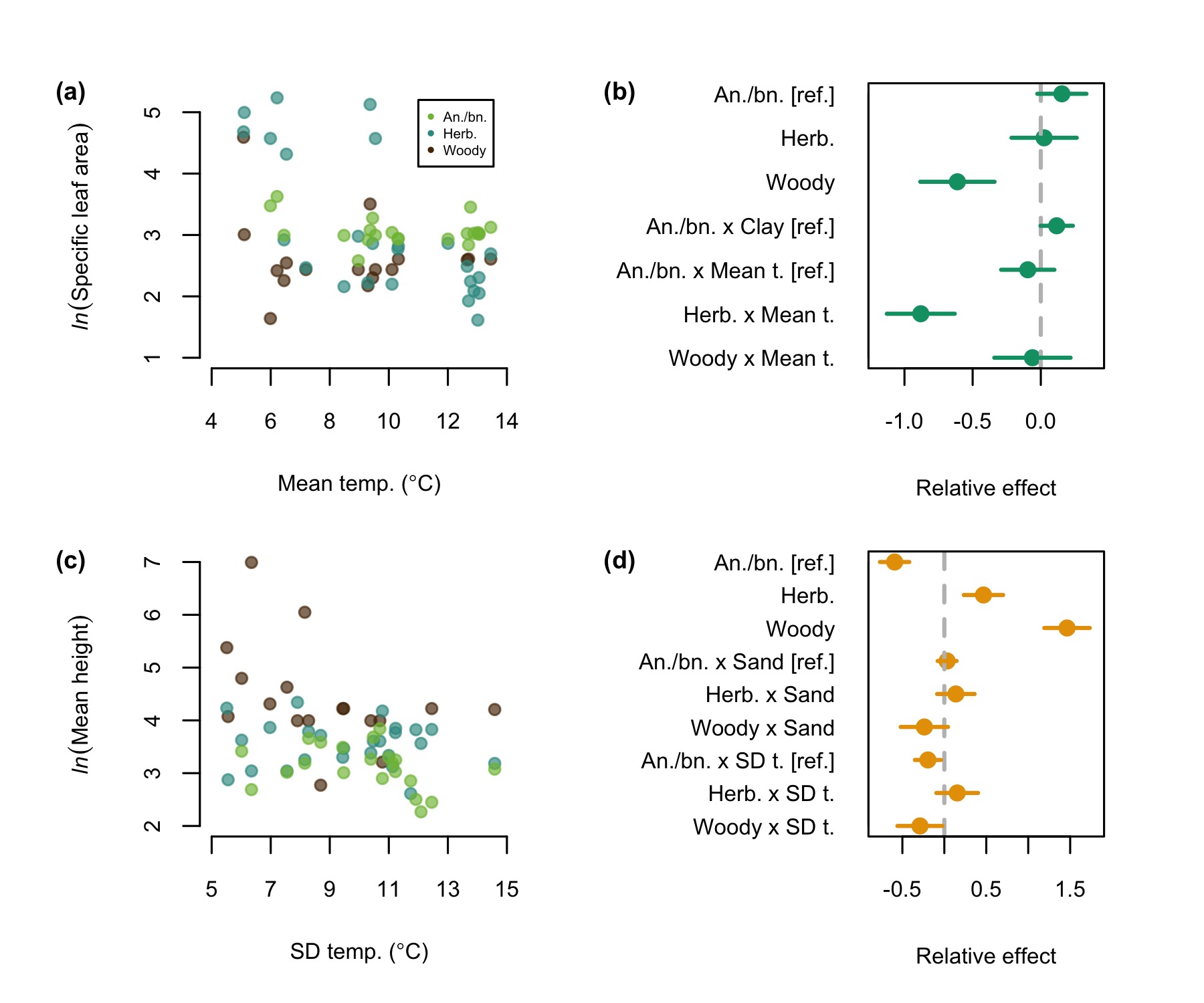

### SUPP-fdiv-env-core18.jpeg

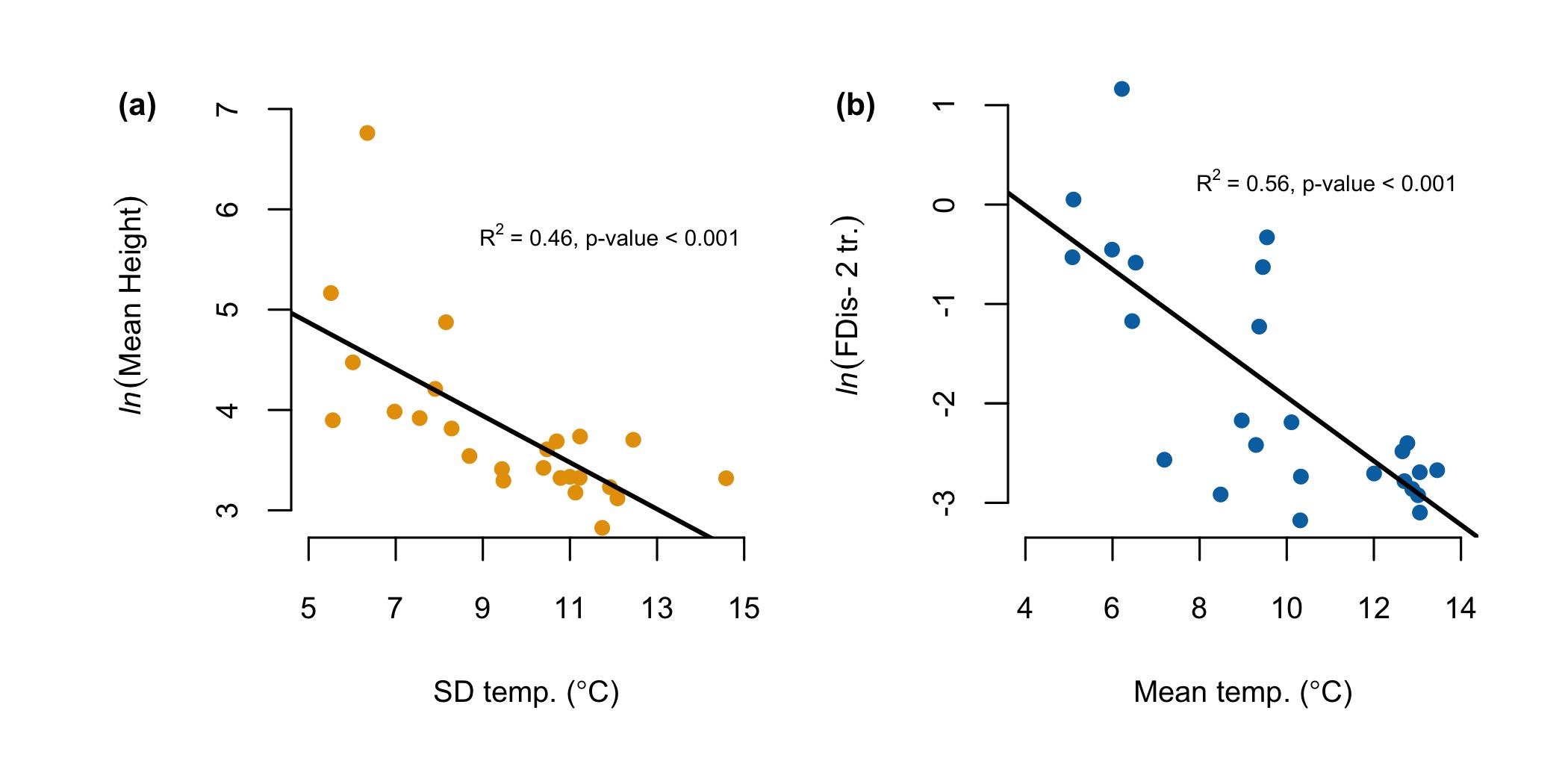
